# Tissue-specific versus pleiotropic enhancers within the *bric-a-brac* tandem gene duplicates display differential regulatory activity and evolutionary conservation

**DOI:** 10.1101/2021.03.25.436949

**Authors:** Henri-Marc G. Bourbon, Mikhail H. Benetah, Emmanuelle Guillou, Luis Humberto Mojica-Vazquez, Aissette Baanannou, Sandra Bernat-Fabre, Vincent Loubiere, Frédéric Bantignies, Giacomo Cavalli, Muriel Boube

## Abstract

During animal evolution, de novo emergence and modifications of pre-existing transcriptional enhancers have contributed to biological innovations, by implementing gene regulatory networks. The *Drosophila melanogaster bric-a-brac* (*bab*) complex, comprising the tandem paralogous genes *bab1*-*2*, provides a paradigm to address how enhancers contribute and co-evolve to regulate jointly or differentially duplicated genes. We previously characterized an intergenic enhancer (named LAE) governing *bab2* expression in leg and antennal tissues. We show here that LAE activity also regulates *bab1*. CRISPR/Cas9-mediated LAE excision reveals its critical role for *bab2*-specific expression along the proximo-distal leg axis, likely through paralog-specific interaction with the *bab2* gene promoter. Furthermore, LAE appears involved but not strictly required for *bab1*-*2* co-expression in leg tissues. Phenotypic rescue experiments, chromatin features and a gene reporter assay reveal a large “pleiotropic” *bab1* enhancer (termed BER) including a series of *cis*-regulatory elements active in the leg, antennal, wing, haltere and gonadal tissues. Phylogenomics analyses indicate that (i) *bab2* originates from *bab1* duplication within the Muscomorpha sublineage, (ii) LAE and *bab1* promoter sequences have been evolutionarily-fixed early on within the Brachycera lineage, while (iii) BER elements have been conserved more recently among muscomorphans. Lastly, we identified conserved binding sites for transcription factors known or prone to regulate directly the paralogous *bab* genes in diverse developmental contexts. This work provides new insights on enhancers, particularly about their emergence, maintenance and functional diversification during evolution.

**Author summary:** Gene duplications and transcriptional enhancer emergence/modifications are thought having greatly contributed to phenotypic innovations during animal evolution. However, how enhancers regulate distinctly gene duplicates and are evolutionary-fixed remain largely unknown. The *Drosophila bric-a-brac* locus, comprising the tandemly-duplicated genes *bab1*-*2*, provides a good paradigm to address these issues. The twin *bab* genes are co-expressed in many tissues. In this study, genetic analyses show a partial co-regulation of both genes in the developing legs depending on tissue-specific transcription factors known to bind a single enhancer. Genome editing and gene reporter assays further show that this shared enhancer is also required for *bab2*-specific expression. Our results also reveal the existence of partly-redundant regulatory functions of a large pleiotropic enhancer which contributes to co-regulate the *bab* genes in distal leg tissues. Phylogenomics analyses indicate that the *Drosophila bab* locus originates from duplication of a dipteran *bab1*-related gene, which occurred within the Brachycera (true flies) lineage. *bab* enhancer and promoter sequences have been differentially-conserved among Diptera suborders. This work illuminates how transcriptional enhancers from tandem gene duplicates (i) differentially interact with distinct cognate promoters and (ii) undergo distinct evolutionary changes to diversifying their respective tissue-specific gene expression pattern.

## Introduction

Gene duplications have largely contributed to create genetic novelties during evolution (1, 2). Intra-species gene duplicates are referred to as “paralogs”, which eventually diverged functionally during evolution in a phylogenetic manner. Gene family expansion has facilitated phenotypic innovation through (i) acquisition of new molecular functions or (ii) the subdivision of the parental gene function between the duplicate copies (3–5). Phenotypic novelties are thought having originated mainly from evolutionary emergence or modifications of genomic *<u>C</u>is*-<u>R</u>egulatory <u>E</u>lements (CREs) or modules, most often dubbed as “enhancer” regions, which regulate gene transcription in a stage-, tissue- and/or cell-type-specific manner (6–10). How CRE (enhancers) within gene complexes (i) are distinctly interacting with their cognate promoters and (ii) are differentially (co-)evolving remain largely unknown.

The *Drosophila melanogaster bric-a-brac* (*bab*) locus comprises two tandemly-duplicated genes (Fig 1A), *bab1*-*2*, which encode paralogous transcription factors sharing two conserved domains: (i) a <u>B</u>ric-a-brac/<u>T</u>ramtrack/<u>B</u>road-complex (BTB) domain involved in protein-protein interactions, and (ii) a specific DNA-binding domain (referred to as BabCD, for <u>B</u>ab <u>C</u>onserved <u>D</u>omain), in their amino(N)- and carboxyl(C)-terminal moieties, respectively (11). Bab1-2 proteins are co-expressed in many tissues (11, 12). In the developing abdominal epidermal cells, within so-called histoblast nests, they jointly regulate directly *yellow* expression in a sexually-dimorphic manner, thus controlling adult male versus female body pigmentation traits (13–16). *bab1*-*2* co-expression in the developing epidermal histoblast nests is partially governed by two CREs which drive reporter gene expression (i) in a monomorphic pattern in the abdominal segments A2-A5 of both sexes (termed AE, for “<u>A</u>nterior <u>E</u>lement”), and (ii) in a female-specific pattern in the A5-A7 segments (DE, for “<u>D</u>imorphic <u>E</u>lement”) (Fig 1A) (14, 17). In addition to controlling male-specific abdominal pigmentation traits, *bab1*-*2* are required, singly, jointly or in a partially-redundant manner, for embryonic cardiac development, sexually-dimorphic larval somatic gonad formation, salivary glue gene repression, female oogenesis, wing development as well as distal leg (tarsal) and antennal segmentation (11, 13, 17–24). In addition to abdominal AE and DE, two other *bab* enhancers, termed CE and LAE (see Fig 1A), have been characterized, which recapitulate *bab2* expression in embryonic cardiac cells and developing tarsal as well as distal antennal cells, respectively (17, 21, 25).

**Fig 1.**
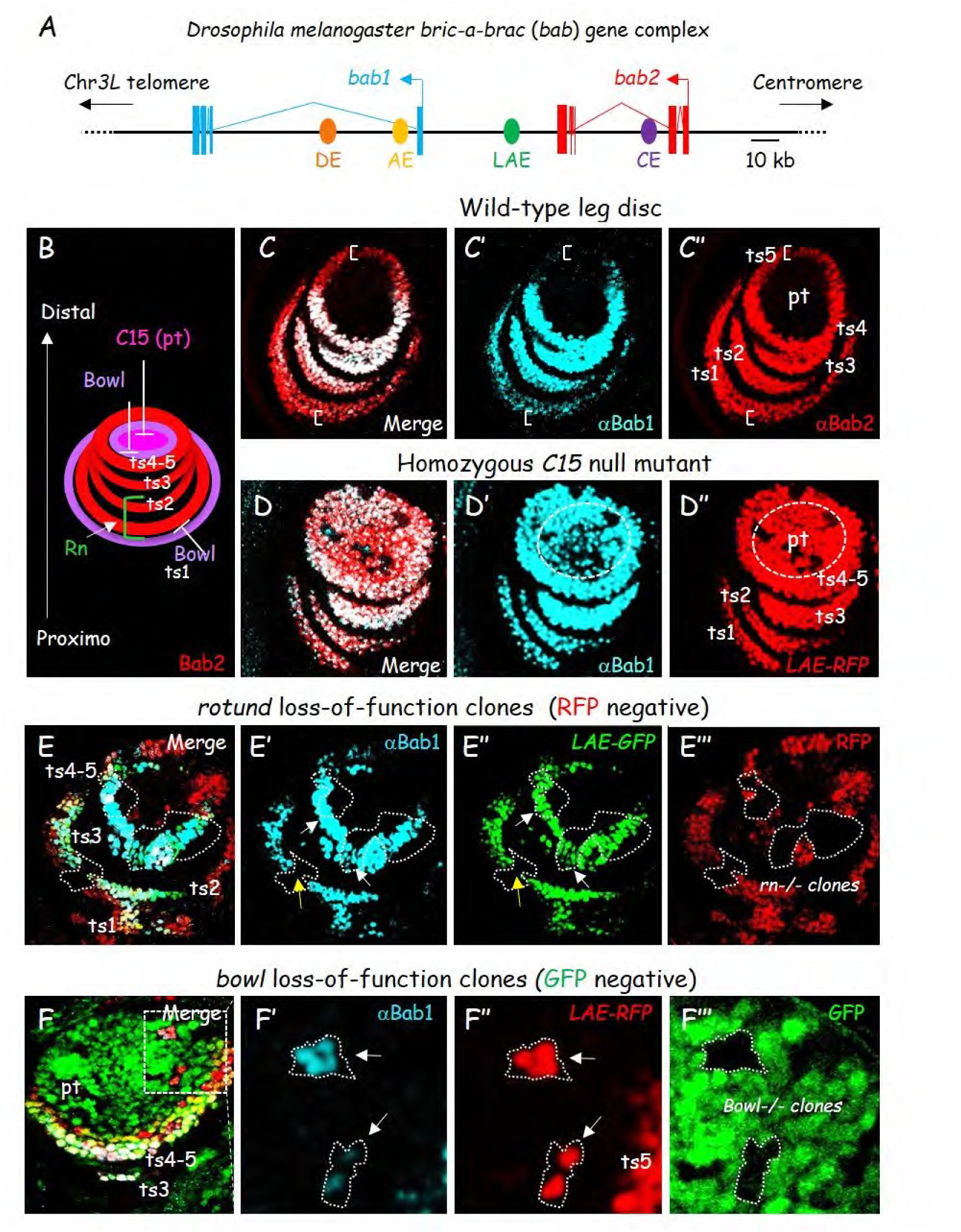
*C15*, r*otund* and *bowl* all regulate both *bab1* and *bab2*. **(A)** Schematic view of the *Dmel bab* locus on the 3L chromosomal arm (Chr*3L*). The tandem *bab1* (blue) and *bab2* (red) transcription units (filled boxes and broken lines represent exons and introns, respectively), the previously known CRE/enhancers are depicted by filled dots (abdominal DE and AE in dark and light orange, respectively; leg/antennal LAE in dark green and cardiac CE in purple), and the telomere and centromere directions are indicated by arrows. **(B)** A scheme depicted C15, Bowl and Rn TF activities in regulating *bab2* expression as a four-ring pattern within the developing distal leg, is shown. **(C)** Medial confocal view of a wild-type L3 leg disc. Merged Bab1 (blue) and Bab2 (red) immunostainings, as well as each marker in isolation in (C’) and (C”), respectively, are shown. Positions of *bab2*-expressing ts1-5 cells and the pretarsal (pt) field are indicated in (C”). Brackets indicate paralog-specific expression in proximalmost and distalmost *bab2*-expressing cells. **(D)** Distal confocal view of a homozygous *C15^2^* mutant L3 leg disc expressing *LAE-RFP*. Merged Bab1 immunostaining (in blue) and RFP fluorescence (red), and each marker in isolation in (D’) and (D”), are shown. Bab2-expressing mutant pt cells are circled with a dashed line in (D’) and (D”). **(E)** Medial confocal view of a mosaic L3 leg disc expressing *LAE-GFP* and harboring *rotund* mutant clones. Merged Bab1 (blue) immunostaining, GFP (green) and RFP (red) fluorescence, as well as each marker in isolation in (E’), (E”) and (E”’), respectively, are shown. Mutant clones are detected as black areas, owing to the loss of RFP. The respective ts1-5 fields are indicated in (E). White and yellow arrows indicate *bab1* (*bab2*) still- and non-expressing *rotund-/-* clones, respectively. **(F)** Distal confocal view of a mosaic L3 leg disc expressing *LAE-RFP* and harboring *bowl* mutant clones. Merged Bab1 (blue) immunostaining, RFP (red) and GFP (green) fluorescence, as well as a higher magnification of the boxed area for each marker in isolation in (F’), (F”) and (F”’), respectively, are shown. Mutant clones are detected as black areas, owing to the loss of GFP. White arrows indicate pretarsal *bowl-/-* clones ectopically expressing both *bab1* and *LAE-RFP* (*bab2*).

Adult T1-3 legs, on the pro-, meso- and meta-thoraces, respectively, are derived from distinct mono-layered epithelial cell sheets, organized as sac-like structures, called leg imaginal discs (hereafter simply referred to as leg discs) (26–28). Upon completion of the third-instar larval stage (L3), each leg disc is already patterned along the proximo-distal (P-D) axis through regionalized expression of the Distal-less (Dll), Dachshund (Dac) and Homothorax (Hth) transcriptional regulators in the distal (center of the disc), medial and proximal (peripheral) regions, respectively (26). The five (ts1-5) tarsal and the single pretarsal (distalmost) segments are patterned through genetic cascades mobilizing transcription factors, notably the distal selector protein Dll and the tarsal Rotund protein as well as nuclear effectors of Notch and <u>E</u>pidermal <u>G</u>rowth <u>F</u>actor <u>R</u>eceptor (EGFR) signaling, i.e., Bowl and C15, respectively (26, 27).

While both *bab* genes are required for dimorphic abdominal pigmentation traits and somatic gonad specification (13, 22), only *bab2* is critical for tarsal segmentation (11). While *bab1* loss-of-function legs are apparently wild-type, a null allele (*bab^AR07^*) removing *bab2* (and *bab1*) activities causes segmental transformation along the P-D leg axis, notably sex comb teeth in tarsal segments ts2-3 of male forelegs, normally only found in ts1, as well as ts2-5 tarsal fusions in both genders (11). While the two *bab* genes are co-expressed within ts1-4 cells, *bab2* is expressed more proximally than *bab1* in ts1, and in a graded manner along the P-D leg axis in ts5 (11, 29). We previously showed that *bab2* expression in distal leg (and antennal) tissues is governed by a 567-bp-long CRE/enhancer (termed LAE for “<u>L</u>eg and <u>A</u>ntennal <u>E</u>nhancer”) which is situated between the *bab1*-*2* transcription units (Fig 1A) (17, 25). However, LAE enhancer contribution to *bab1*-*2* co-regulation in the developing distal legs remains to be investigated in tarsal segments ts3-4 where expression levels of both paralogous BTB-BabCD proteins are the highest (see Fig 1B) (11).

Here, we show that *bab1* expression in the developing distal leg also depends on the Rotund, Bowl and C15 proteins, three transcription factors known to regulate directly *bab2* expression, by binding to dedicated LAE sequences (17, 25). LAE excision by CRISPR/Cas9-mediated genome editing indicates that this enhancer is partly involved in *bab1*-*2* co-regulation and, more unexpectedly, is also required for their differential expression along the P-D leg axis. Additionally, we show that LAE acts redundantly with a large enhancer signature region (termed BER), located within the *bab1* transcription unit, which is bound by dedicated transcription factors involved in diverse developmental processes and thus BER is prone to act as a “pleiotropic” enhancer region. Our phylogenomics analyses indicate that LAE and *bab1* promoter sequences have been fixed early on during dipteran evolution, well before *bab1* duplication. Conversely, BER and *bab2* promoter sequences have been fixed much later. Lastly, within *D. melanogaster* BER, we identified conserved binding sites for many transcriptional regulators known or prone to regulate *bab1* and/or *bab2* expression in the developing leg and antenna, but also in wing, haltere, mesodermal and gonadal tissues. This work illuminates how transcriptional enhancers from tandem gene duplicates (i) differentially interact with distinct cognate promoters and (ii) undergo distinct evolutionary changes to diversifying their respective tissue-specific gene expression pattern.

## Results

### The tandem *bab1*-*2* gene paralogs are co-regulated in the developing distal leg

In addition to the distal selector homeodomain (HD) protein Distal-less, we and others have previously shown that the C15 HD protein (homeoprotein) as well as Rotund and Bowl <u>Z</u>inc-<u>F</u>inger (ZF) transcription factors (TFs) bind dedicated sequences within LAE to ensure precise *bab2* expression in four concentric tarsal rings within the leg discs (Fig 1B) (17, 25). *bab1*-*2* are co-expressed in ts2-4 tarsal segments, while *bab2* is specifically expressed in ts5 and more proximally than *bab1* in ts1, both in a graded manner along the P-D leg axis (Fig 1C and S1A Fig) (11). Given *bab1*-*2* co-expression in ts1-4, we first asked whether *C15*, *rotund* and *bowl* activities are also controlling *bab1* expression in the developing distal leg. To this end, we compared Bab1 expression with that of a X-linked *LAE-GFP* (or *LAE-RFP*) reporter gene faithfully reproducing the *bab2* expression pattern there (17, 25), in homozygous mutant leg discs for a null *C15* allele or in genetically-mosaic leg discs harboring *rotund* or *bowl* loss-of-function mutant cells (Fig 1D-F).

*C15* is specifically activated in the distalmost (center) part of the leg disc giving rise to the pretarsal (pt) segment (see Fig 1B) (30, 31). We have previously shown that the C15 homeoprotein down-regulates directly *bab2* to restrict its initially broad distal expression to the tarsal segments (25). Bab1 expression analysis in a homozygous *C15* mutant leg disc revealed that both *bab1* and *LAE-RFP* (*bab2*) are similarly de-repressed in the pretarsus (Fig 1, compare panels C-D).

In contrast to *C15*, *rotund* expression is restricted to the developing tarsal segments (32) and the transiently-expressed Rotund ZF protein contributes directly to *bab2* up-regulation in proximal (ts1-2) but has no functional implication in distal (ts3-5) tarsal cells (17). Immunostaining of genetically-mosaic leg discs at the L3 stage revealed that *bab1* is cell-autonomously down-regulated in large *rotund* mutant clones in ts1-2, but not in ts3-4 segments (Fig 1E), as it is the case for *LAE-GFP* reflecting *bab2* expression. Lastly, we examined whether the Bowl ZF protein, a repressive TF active in pretarsal but not in most tarsal cells, is down-regulating *bab1* expression there (33), like *bab2* (25). Both *bab1* and *LAE-RFP* (*bab2*) appeared cell-autonomously de-repressed in *bowl* loss-of-function pretarsal clones (Fig 1F).

In addition to loss-of-function, we also conducted gain-of-function experiments for *bowl* and *rotund*. Bowl TF gain-of-function was achieved by down-regulating *lines* which encodes a related but antagonistic ZF protein (i) destabilizing nuclear Bowl and is specifically expressed in the tarsal territory (33). As previously shown for *LAE-GFP* (and *bab2*) expression, nuclear Bowl stabilization in the developing tarsal region appears sufficient to down-regulate cell-autonomously *bab1* (S1C Fig). Prolonged expression of the Rotund protein in the entire distal part of the developing leg disc, i.e., tarsal in addition to pretarsal primordia, induces ectopic *bab1* expression in the presumptive pretarsal territory, albeit with some differences with *bab2* expression (S1B Fig, differentially-expressing cells are indicated with arrows), thus suggesting differential sensitivity of the two gene duplicates to Rotund TF levels (see discussion).

Taken together, these data indicate that the C15, Bowl and Rotund transcription factors, previously shown to interact physically with specific LAE sequences and thus to regulate directly *bab2* expression in the developing distal leg, are also regulating *bab1* expression there. These results suggest that the limb-specific intergenic LAE enhancer activity regulates directly both *bab* genes.

### LAE activity regulates both *bab1* and *bab2* gene paralogs along the proximo-distal leg axis

To test the role of LAE in regulating *bab1-2,* we deleted precisely the LAE sequence through CRISPR/Cas9 genome editing (see Materials and Methods) (Fig 2A). Two independent deletion events (termed *ΔLAE-M1* and *-M2*; see S2 Fig for deleted DNA sequences) were selected for phenotypic analysis. Both are homozygous viable and give rise to fertile adults with identical fully-penetrant distal leg phenotypes, namely ectopic sex-comb teeth on ts2 (normally only found on ts1) tarsal segment in the male prothoracic (T1) legs (Fig 2B), which are typical of *bab2* hypomorphic alleles (11). The *ΔLAE-M1* allele was selected for detailed phenotypic analyses and is below referred to as *bab^ΔLAE^*.

**Fig 2.**
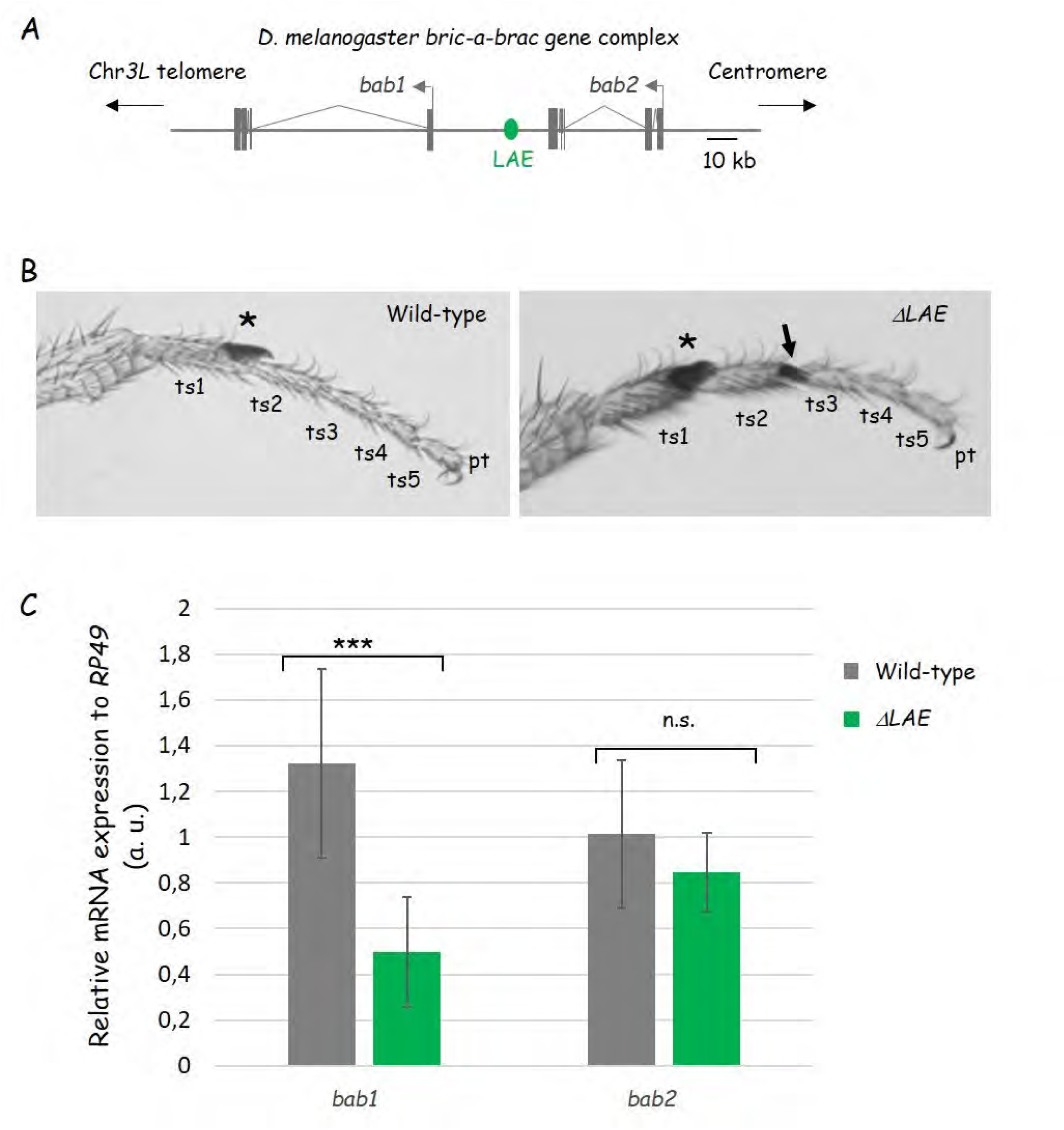
LAE is not critically required for tarsal segmentation and for overall *bab2* expression in the leg disc. **(A)** Schematic view of the *Dmel bab* locus on the 3L chromosomal arm (Chr*3L*). The tandem *bab1* and *bab2* transcription units (filled boxes and broken lines represent exons and introns, respectively), the intergenic LAE enhancer (in green), as well as the telomere and centromere directions, are depicted as in Fig 1A, except that both genes are depicted in grey. The small CRISPR/Cas9-mediated chromosomal deficiency (*bab^ΔLAE^*) is shown in beneath (deleted LAE is depicted as a broken line). **(B)** Photographs of wild-type and homozygous *bab^ΔLAE^* T1 distal legs from adult males. The regular sex-comb (an array of about 10 specialized bristles on the male forelegs) on distal ts1 is indicated with asterisks, while ectopic sex-comb bristles on distal ts2 from the mutant leg (right) is indicated by an arrow. Note that the five tarsal segments remain individualized in homozygous *bab^ΔLAE^* mutant legs. **(C)** Overall *bab1-2* expression from wild-type and homozygous *bab^ΔLAE^* L3 leg discs, as determined from reverse transcription quantitative PCR analyses. The *bab1-2* expression levels were quantified relative to *rp49* mRNA abundance.

First, we quantified *bab1* and *bab2* mRNAs prepared from dissected wild-type and homozygous *bab^ΔLAE^* mutant leg discs. As shown in Fig 2C, both mRNAs were detected in mutant discs, although *bab1* levels were two times lower than wild-type. Second, Bab1-2 expression patterns were analyzed in homozygous *bab^ΔLAE^* leg discs (Fig 3). To identify leg cells that should normally express *bab2*, we used the X-linked *LAE-GFP* reporter. In homozygous *bab^ΔLAE^* mutant leg discs, *bab2* specific expression (see Fig. 1B) is no longer observed (Fig 3B-C), while Bab1-2 shared expression is very low in ts3-4 to undetectable in ts1-2. Nevertheless, residual *bab1*-*2* co-expression in homozygous *bab^ΔLAE^* mutant discs indicates that additional *cis*-regulatory region(s) within the *bab* locus act(s), at least partly, redundantly with the LAE enhancer.

**Fig 3.**
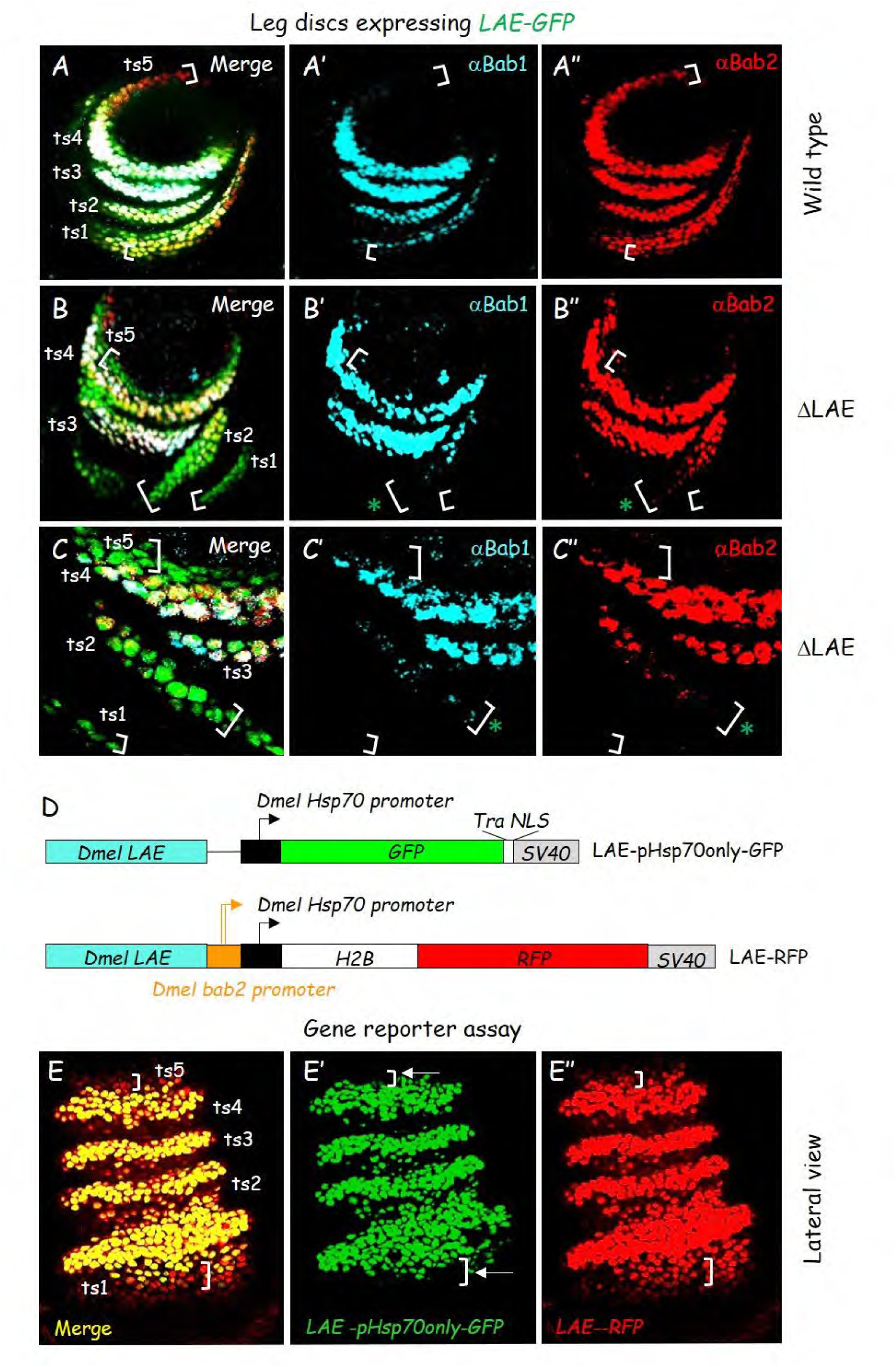
LAE is mostly critically required for paralog-specific *bab2* expression in the developing distal legs. **(A-C)** Medial (A-B) and distal (C) confocal views of wild-type (A) and homozygous *bab^DLAE^* mutant (B and C) L3 leg discs expressing *LAE-GFP*. Merged GFP fluorescence (green), Bab1 (blue) and Bab2 (red) immunostainings, as well as the two latter in isolation in (A’-C’) and (A”-C”), respectively, are shown. Brackets indicate positions of paralog-specific expression in proximalmost (ts1) and distalmost (ts5) *bab2*-expressing (GFP+) cells. Green asterisks in (B’-B”) and (C’-C”) indicate weaker expression of both *bab* paralogs in GFP+ ts2 cells. **(D)** Modular structures of the *LAE-pHsp70onlyGFP* and *LAE-RFP* reporter constructs. GFP (green box) and RFP (red box) open-reading frames (ORFs) have been fused with ORFs for the Transformer (Tra) nuclear localization signal (NLS) and the histone H2B, respectively (see white boxes). The *SV40* polyadenylation signal region is boxed in grey. The *Dmel* LAE sequence is boxed in blue. The classical non-heat-inducible basal *Hsp70* promoter sequence is boxed in black, while the *bab2* core promoter sequence is depicted in orange. Note that both promoters are juxtaposed in the *LAE-RFP* construct. **(E)** LAE activity requires functionally the *bab2* promoter to ensure paralog-specific expression in the developing legs. A lateral confocal view of merged GFP (green) and RFP (red) fluorescence, as well as each marker in isolation in (E’) and (E”), respectively, of the distal part of an early pupal leg expressing both the *LAE-pHsp70onlyGFP^ZH2A^* and *LAE-RFP^ZH86Fb^* reporter constructs (depicted in (D)), are shown. Brackets indicate tarsal RFP+ cells expressing *bab2* in a paralog-specific manner, which never express the *LAE-pHsp70onlyGFP^ZH2A^* reporter which lacks *bab2* core promoter sequences (see white arrows).

Taken together, our data indicate that LAE enhancer activity is (i) required for *bab1*-*2* co-expression in the two proximal-most tarsal segments, particularly ts1, (ii) dispensable for their co-expression in ts3-4, suggesting the presence of redundant *cis*-regulatory information and (iii) critically required for *bab2*-specific tarsal expression both proximally and distally. Thus, LAE activity governs both shared and paralog-specific expression of the *bab1*-*2* gene duplicates.

### LAE paralog-specific activity requires the *bab2* core promoter

Whereas enhancer emergence has been proposed to account for acquisition of novel tissue- or paralog-specific functions for gene duplicates (34–36), LAE regulatory function provides an example of a single enhancer responsible both for shared and differential expression of two tandemly-repeated gene paralogs. Previously tested LAE reporter constructs fused *bab2* core promoter sequences to the minimal *Hsp70* promoter region (*pHsp70*) (17, 25). To examine the contribution of the *bab2* promoter to LAE activity we compared the expression of two LAE reporters containing (*LAE-RFP*) or not (*LAE-pHsp70only-GFP*) the *bab2* promoter sequence (Fig 3D). Strikingly, the *LAE-pHsp70only-GFP* reporter was no longer activated in *RFP+* (*bab2*-expressing) ts1 and ts5 cells (Fig 3E; see white brackets and arrows). These data indicate that *bab2*-specific regulation by LAE activity requires the *bab2* core promoter sequences.

### In addition to the intergenic LAE, other leg-specific enhancer elements are present within the *bab1* first intron

Since LAE appeared dispensable for *bab1-2* co-expression in ts3-4 cells, our data suggested the existence of other redundant *cis*-regulatory elements, presumably located also within the *bab* locus. On one side, a X-linked <u>B</u>acterial <u>A</u>rtificial Chromosome (BAC) construct, *BAC26B15^ZH2A^*, encompassing the *bab2* gene and downstream intergenic sequence including LAE (see Fig 4A), could rescue (i) Bab2 expression in the tarsal primordium and (ii), distal leg phenotypes detected in homozygous animals for the null allele *bab^AR07^* (17). On the other side, a *BAC26B15* construct (*BAC26B15ΔLAE ^ZH2A^*) inserted at the same genomic landing site and specifically lacking LAE sequence did not (Fig 4B-D). These results confirmed that (i) in absence of redundant *cis*-regulatory information, LAE is essential for *bab1-2* expression in the tarsal segments and (ii) the *cis*-information redundant with LAE is located outside the genomic region covered by *BAC26B15*.

**Fig 4.**
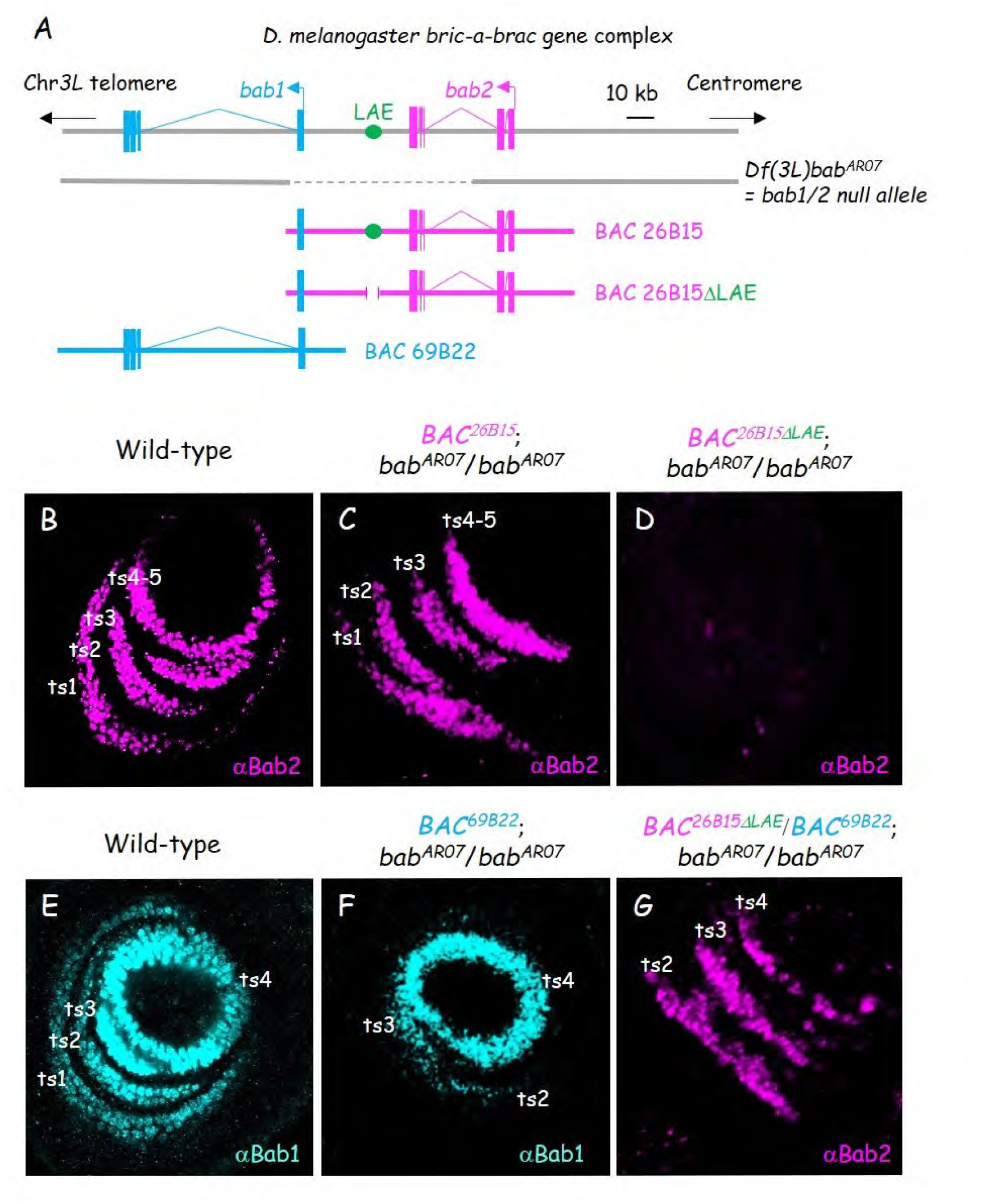
*bab1* includes partially-redundant limb-specific *cis*-regulatory information. **(A)** Chromosomal deficiency and BAC constructs covering the *bab* locus. The tandem gene paralogs and intergenic LAE are depicted as shown in Fig. 1A, except that *bab2* is depicted in pink instead of red. The *bab^AR07^ 3L* chromosomal deficiency is shown in beneath, with known deleted portion indicated by a dashed line. Note that the breakpoints have not been precisely mapped. The two overlapping BAC constructs *69B22* and *26B15*, as well as a mutant derivative of the latter specifically-deleted for LAE, are shown further in beneath. **(B-G)** Medial confocal views of wild-type (B-E) and homozygous *bab^AR07^* mutant (C-D and F-G) L3 leg discs, harboring singly or combined X-linked BAC construct(s) shown in (A), as indicated above each panel. Bab2 (pink) and Bab1 (blue) immunostainings are shown. Positions of *bab1*- and *bab2*-expressing ts1-4 cells are indicated. Note stochastic *bab2* expression in (G).

To identify limb-specific redundant *cis*-regulatory information within the *bab* complex, we first tested the capacity of another BAC, *BAC69B22*, which overlaps *bab1* and lacks LAE (see Fig 4A), to restore Bab1 expression in *bab^AR07^* mutant leg discs. As shown in Fig 4E-F, the X-linked *BAC69B22^ZH2A^* could restore *bab1* expression in ts2-4, indicating that it contains *cis*-regulatory information redundant with LAE activity in these segments. To test the capacity of *BAC69B22* sequences to also regulate *bab2* expression in ts2-4, we placed *BAC69B22^ZH2A^* across *BAC26B15ΔLAE^ZH2A^,* to allow pairing-dependent *trans*-interactions (i.e., transvection) between the two X chromosomes in females. This configuration partially restored Bab2 expression in ts2-4 cells from *bab^AR07^* mutant L3 leg discs, albeit in salt and pepper patterns (Fig 4G), diagnostic of transvection effects (37).

From these data, we predicted the existence of *cis-*regulatory information within the *69B22* chromosomal interval capable to drive some *bab1-2* expression in distal leg tissues and acting redundantly with the LAE enhancer.

### Chromatin features predict limb-specific *cis*-regulatory elements within *bab1*

Next, we sought to identify *cis*-regulatory information acting redundantly with LAE by taking advantage of available genome-wide chromatin features and <u>Hi</u>gh-throughput chromosome conformation <u>C</u>apture (Hi-C) experiments performed from L3 eye-antennal and/or leg discs (Fig 5). *bab1-2* are indeed co-expressed in distal antennal cells within the composite eye-antennal imaginal disc (11). A topologically-associating domain covering the entire *bab* locus was detected in Hi-C data from eye-antennal discs (Fig 5A and S3A Fig) (38), revealing particularly strong interactions between *bab1*-*2* promoter regions.

**Fig 5.**
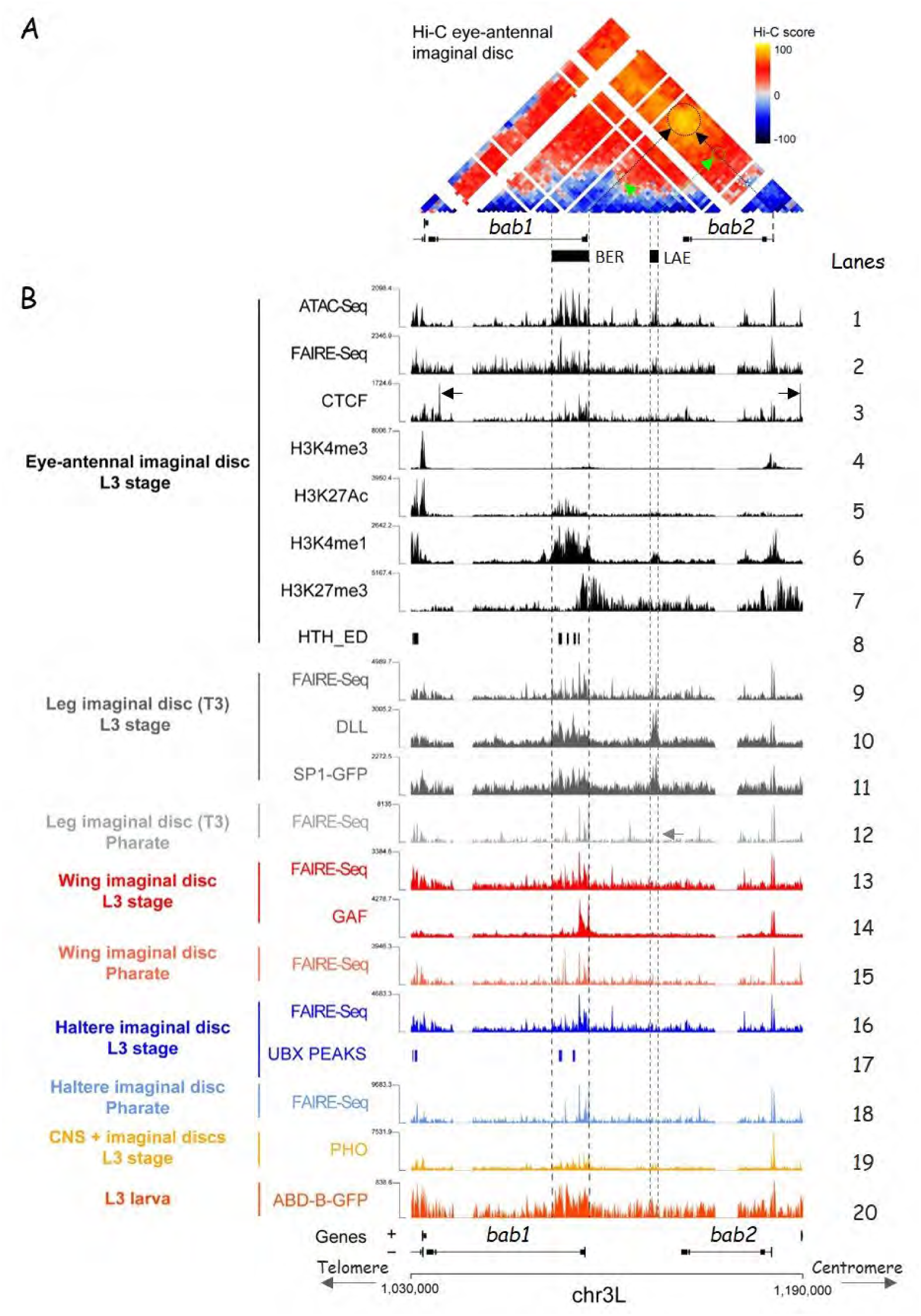
A topologically-associating domain encompasses the *bab* locus in the eye-antennal disc and genome-wide chromatin features identify an enhancer signature region (BER) within *bab1*. **(A)** Hi-C screenshot of a 160 kb region covering the *Dmel bab* gene complex. Score scale is indicated on the right (yellow to dark blue from positive to negative). **(B)** ATAC-, FAIRE- and/or ChIP-Seq profiles from L3 eye-antennal (ED), leg, wing and haltere discs as well as from adult pharate appendages (leg, wing and haltere) and from whole larval tissues, as indicated on the left side. As referred in the main text, lanes are numbered on the right side. ChIP-Seq peak calling data are shown in lanes 8, 17-18. Otherwise, normalized open chromatin, histone H3 post-translational modifications and TF binding profiles are shown. Positions of the tandem *bab1*-*2* genes are indicated on the bottom. The respective locations of the BER and LAE sequences are highlighted with vertical dashed lines. Of note, according to normalized FAIRE-Seq signals, LAE is not fully accessible in the pharate T3 leg (see grey arrow in lane 12). Strongest CTCF ChIP-Seq signals are indicated by horizontal black arrows (lane 3).

We then used published genome-wide data from <u>Ch</u>romatin Immuno-<u>P</u>recipitation (ChIP-Seq), <u>F</u>ormaldehyde-<u>A</u>ssisted Isolation of <u>R</u>egulatory <u>E</u>lements (FAIRE-Seq) and <u>A</u>ssay for <u>T</u>ransposase-<u>A</u>ccessible <u>C</u>hromatin (ATAC-Seq) experiments (38–41), looking for active enhancer marks (H3K4me1 and H3K27Ac) and nucleosome-depleted chromatin regions (thus accessible to transcription factors). Active enhancer signatures are mainly associated with a ∼15-kb-long genomic region that we termed BER, for “*<u>b</u>ab1* <u>E</u>nhancer <u>R</u>egion”, encompassing the *bab1* promoter, first exon and part of its first intron (Fig 5B, lanes 1-2 and 5-6, respectively; see also S3B Fig for peak calling data). Note that LAE is also accessible to transcription factors and carries H3K4me1 marks, consistently with enhancer activity in distal antennal cells (17).

To more precisely locate putative enhancer element(s) within BER, we analyzed previously-published ChIP-Seq data from L3 leg discs (42) for binding sites for Dll, Sp1 and Hth proteins, known to regulate *bab1* and/or *bab2* expression in the developing legs (17, 42–44). Strong Dll binding is detected throughout BER, including over the *bab1* promoter (Fig 5B, lane 10; see also S3B Fig). In leg discs, Dll binding is detected over 8 out of 10 <u>O</u>pen <u>C</u>hromatin <u>S</u>ubregions (OCS) within BER (Fig 6C and S3B Fig) and six of those eight are also bound by Sp1 ZF protein. Of note, all nucleosome-depleted (i.e., OCS) BER subregions in the leg are also accessible in the eye-antennal discs (40) (Fig 5B, compare lanes 2 and 9, and Fig 6C, two upper lanes). Lastly, of six OCS sequences co-bound by Dll and Sp1, four are also bound by Hth protein (Fig 5B, lane 8, and Fig 6C, bottom lane). FAIRE-Seq data indicate that, in addition to LAE, only OCS7 is nucleosome-depleted in the leg and eye-antennal discs but not the wing and/or haltere discs (Fig 5B, lanes 13, 16, 18, and Fig 6C, four upper lines; see also S3B Fig). Importantly, ventral limb-specific OCS7 is co-bound by Dll, Sp1 and Hth transcription factors (Fig 6C). Thus, within the entire *bab* locus, only LAE and BER OCS7 are specifically bound by transcriptional regulators in the developing leg and antenna.

**Fig 6.**
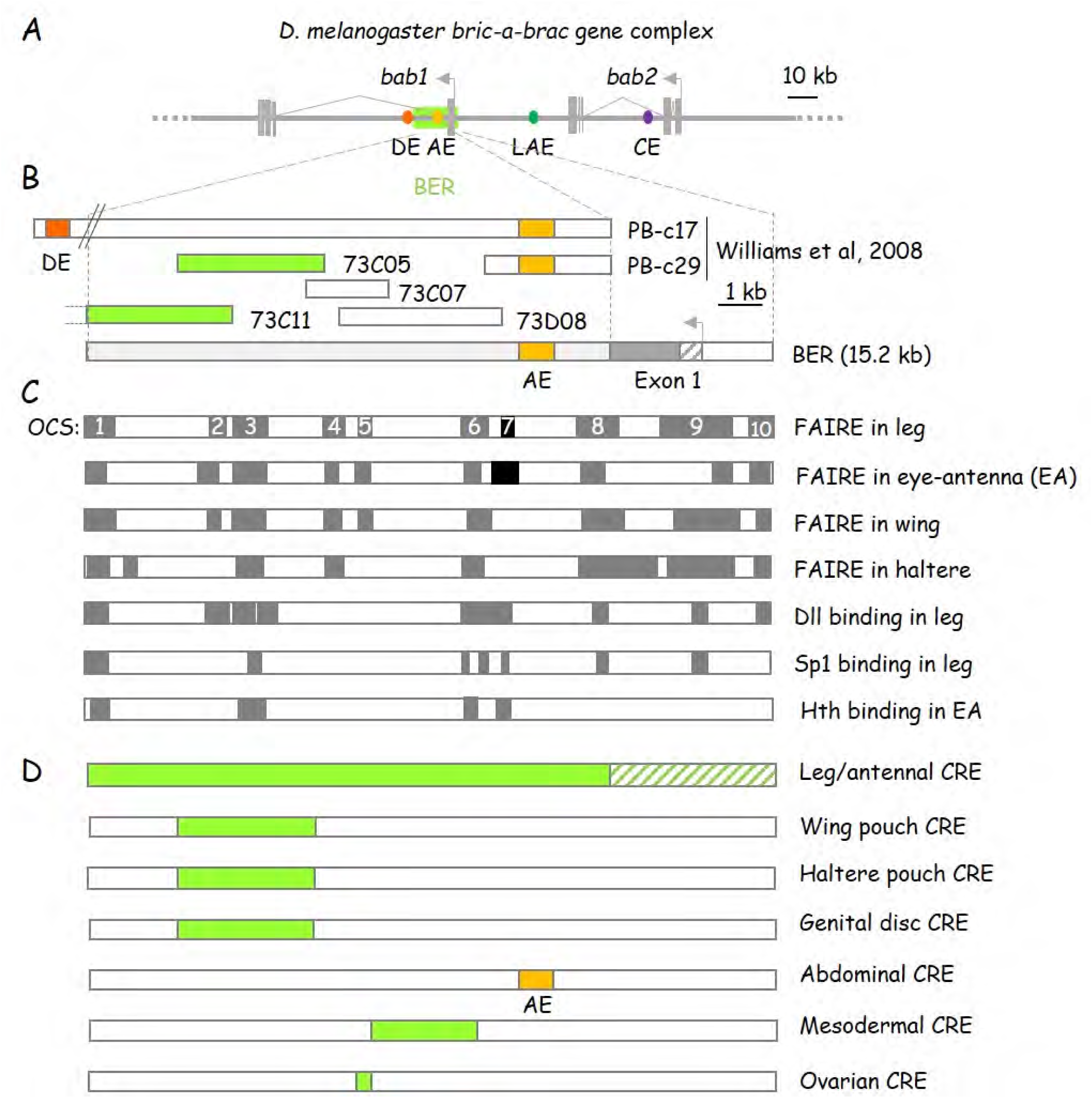
BER behaves as a composite pleiotropic enhancer. **(A)** The BER enhancer signature region includes the abdominal AE enhancer. Organization of the *Dmel bab* locus, with the tandem gene paralogs as depicted as in Fig. 2A. The characterized enhancers are depicted by filled dots (abdominal DE and AE in dark and light orange, respectively; leg/antennal LAE in dark green and cardiac CE in purple). BER is boxed in light green. The genomic portions of the overlapping *69B22* and *26B15* BAC constructs are shown in beneath. **(B)** Transgenic lines covering BER identify *cis*-regulatory elements driving reporter gene expression in diverse larval imaginal tissues. Genomic fragments covered by relevant Janelia Farm FlyLight reporter lines [77] and the DE- and/or AE-containing PB-c17 and PB-c29 genomic constructs, described in [14], are shown above a scheme of the BER region. *bab1* protein coding and 5’-untranslated sequences within the first exon are filled or hatched in dark grey, respectively, while the intronic region is in light gray. The AE sequence is in orange, as depicted in (A). FlyLight reporter lines driving reporter expression in diverse imaginal discs (see S4 Fig) are filled in light green. **(C)** BER includes open chromatin sequences (OCS) and is bound by Dll, Sp1, Hth TFs in diverse developing appendages (leg, eye-antenna, wing and haltere). OCS and TF-bound sequences are depicted by filled grey/black boxes. Numbers refer to OCSs detected in the leg discs (see main text). The black boxes represent OCSs detected in the eye-antennal (EA) and leg but not in wing and haltere discs. OCS and Dll or Sp1-bound regions, as determined from peak calling (FAIRE-Seq GSE38727 and ChIP-Seq GSE113574 GEO dataset series, respectively), are from [40] and [42], respectively. ChIP-Seq data for Hth are from [78]. **(D)** BER includes pleiotropic *cis*-regulatory elements (CREs). Locations of predicted CREs (see text) are indicated by light green boxes. The hatched part of the predicted leg/antennal CRE is inferred from data obtained with the PB-c17 construct reported in [14].

In summary, data mining indicates that BER includes a cluster of enhancer elements bound by Dll and Sp1 in leg discs and thus are good candidates for acting redundantly with LAE in regulating *bab* genes in a ventral limb-specific manner.

### BER includes multiple *cis*-regulatory elements active in diverse developmental contexts

To further ascribe regulatory roles to BER subregions, we took advantage of a systematic analysis of Gal4 reporter lines (45). Out of six lines containing BER fragments (Fig 6B), only two, *73C11* and *73C05*, overlapping OCS1-3, are active in the leg and eye-antennal discs (FlyLight database; (http://flweb.janelia.org/cgi-bin/flew.cgi; S4B Fig). Nevertheless, none reproduce the *bab1/2* leg or antennal expression patterns in four or two concentric distal rings, respectively. These and other published data from reporter constructs including *bab*1 first intron sequences (14) indicate that OCS1-7 (see Fig 6B) are not sufficient to properly drive *bab1/2* expression in the developing legs and suggest the requirement of additional BER elements, particularly the *bab1* promoter region (i.e., OCS9). This hypothesis is consistent with binding of the known *bab1-2* leg regulators Dll and Sp1 throughout BER, in addition to LAE (Fig 5, lane 10; and S3B Fig).

The *73C05* BER fragment also drives reporter gene expression in the wing, haltere and genital discs (S4 Fig, panels E and H-J) in patterns strikingly similar to those described for *bab2* (11, 44). Consistently, FAIRE- and ChIP-Seq data from haltere discs indicate respectively that OCS1-3 are nucleosome-free and bind the Ultrabithorax (Ubx) Hox-type homeoprotein known to activate directly *bab2* expression in haltere tissues (Fig 5B, lanes 16-17, and Fig 6C) (46, 47). Furthermore, ChIP-Seq data from whole L3 larvae (modENCODE), showed binding over the entire BER region of the Hox Abd-B genital selector (Fig5B, lane 20) (48). Lastly, BER includes (i) nucleosome-depleted sequences governing expression in adult muscles and bound by the mesodermal transcription factors Mef2, Slp1 and Tinman in late embryos (14, 49) as well as (ii) a sequence element (overlapping OCS5) which confers enhancer activity in ovarian somatic cells (50) (see Fig 6D).

Taken together, our data indicate that BER sequences drive *bab1*-*2* expression in developing limbs but also in other tissues such as wing, haltere, genitalia and mesoderm. Moreover, owing to the presence of binding sites for transcription factors known to regulate *bab* gene expression in these respective tissues, spread out over the whole BER sequence, the latter is thus proposed to act as a pleiotropic enhancer region.

### Cross- and auto-regulations among the *bab* genes

Bab proteins interact with A/T-rich DNA sequences through their BabCD DNA-binding domain, including binding sites within their own locus (51). We therefore tested whether the Bab1-2 proteins autoregulate and/or cross-regulate their own expression. Previous data indicated that Bab2 protein expression is unaffected in *bab1* loss-of-function mutants (11). Given that protein null *bab2* alleles are not available, we used RNA interference coupled to flip-out (FO) Gal4 expression to down-regulate clonally *bab2* expression within developing legs, and examine *LAE-RFP* and *bab1* expression in mosaic L3 leg discs (Fig 7). Strikingly, both *LAE-RFP* and *bab1* were up-regulated cell-autonomously in most tarsal mitotic clones (n=17/20) (Fig 7, panels A-A”). Moreover, *bab2* down-regulation in proximal-most RFP+ ts1 cells (expressing only *bab2*) activated cell-autonomously *bab1*, in addition to up-regulating *LAE-RFP* expression (Fig 7, white arrows in panels B-B”). These results suggested to us that the *bab2* paralog specifically down-regulates its own expression through partial repression of LAE enhancer activity. To confirm these observations, we generated mutant clones for the *bab^AR07^* null allele, lacking both *bab2* and *bab1* activities. A slight cell-autonomous *LAE-GFP* reporter up-regulation could be observed in all examined *bab^AR07^* clones (detected with anti-Bab2 antibodies; n>20) (Fig 7, panels C-C”), independently of their size and position within *bab2*-expressing tarsal cells (see white arrows and arrowheads).

**Fig 7.**
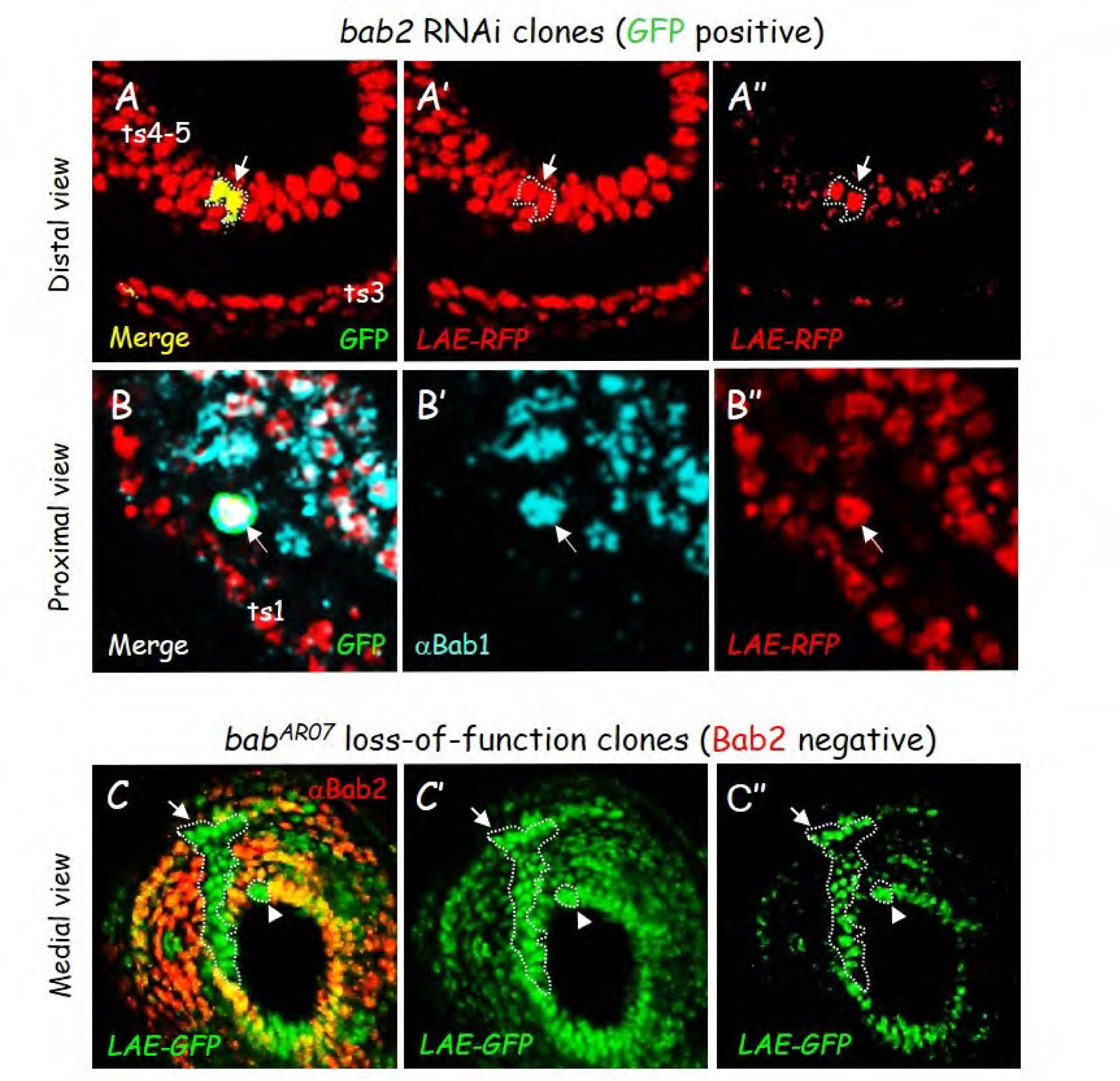
Auto- and cross-regulation among the two *bab* paralogs in the developing legs. **(A)** Distal confocal view of a L3 leg disc expressing *LAE-RFP* and harboring flip-out (FO) clones expressing interfering RNA against *bab2*. Merged RFP (red) and GFP (green) fluorescence, as well as the former in isolation and under two distinct signal magnification (A’ and A”), are shown. A FO clone (GFP+) within *LAE-RFP* (*bab2*)-expressing ts4 cells is circled with a dashed line (see arrow). **(B)** Proximal confocal view of a L3 leg disc expressing *LAE-RFP* and harboring FO clones expressing interfering RNA against *bab2*. Merged Bab1 (blue) immunostaining, RFP (red) and GFP (green) fluorescence, as well as the two formers in isolation, in (B’) and (B”), respectively, are shown. Note that a single-cell FO clone (GFP+), within *LAE-RFP* (*bab2*)-expressing ts1 cells, is sufficient to upregulate *bab1* (see arrows). **(C-D)** L3 leg (C) and eye-antennal (D) discs expressing *LAE-GFP* and harboring mutant clones for the protein null allele *bab^AR07^*. Merged Bab2 (red) immunostaining and GFP (green) fluorescence as well as the latter in isolation under two distinct signal magnifications (C’-C” and D’-D”), are shown. Tiny and larger clones (Bab2 negative) are circled with dashed lines (arrowheads and arrows, respectively).

Altogether, we conclude that *bab2* down-regulates its own expression, likely via partial repression of LAE activity, thus ensuring appropriate levels of both paralogous BTB-BabCD transcription factors in distal leg tissues, and most likely in other appendages as well.

### The *bab* gene complex arose from *bab1* duplication in the Muscomorpha infraorder

The different levels of *cis*-regulatory element redundancy within the *bab* locus led us to trace back the evolutionary origin of the *bab* duplication found in *D. melanogaster* (*Dmel*). To start, we identified proteins orthologous to *Dmel* Bab1 or Bab2, i.e., displaying an N-terminal BTB associated to a C-terminal BabCD domain (collectively referred to as BTB-BabCD proteins) (11) within highly diverse dipteran species (see Fig 8A). Two distinct BTB-BabCD proteins strongly related to *Dmel* Bab1 and Bab2, respectively, were identified in the Muscomorpha (higher flies, also known as Cyclorrhapha) superfamily, both within the Schizophora (in Calyptratae, such as *Musca domestica* and *Glossina morsitans*, and in Acalyptratae, particularly among Drosophilidae) and Aschiza subsections (Fig 8A-B and Supplementary data). In contrast, a single BTB-BabCD protein could be identified in evolutionarily-distant dipteran species within (i) the brachyceran Asilomorpha and Stratiomyomorpha superfamilies (such as *Proctacanthus coquilletti* and *Hermetia illucens*, respectively), collectively referred to as Orthorrhapha; (ii) the Nematocera suborder families (with rare exceptions, in Psychodomorpha and Bibionomorpha, see below); (iii) other Insecta orders (e.g., Coleoptera, Hymenoptera and Lepidoptera), and in crustaceans (e.g., *Daphnia pulex*) (see Supplementary data).

**Fig 8.**
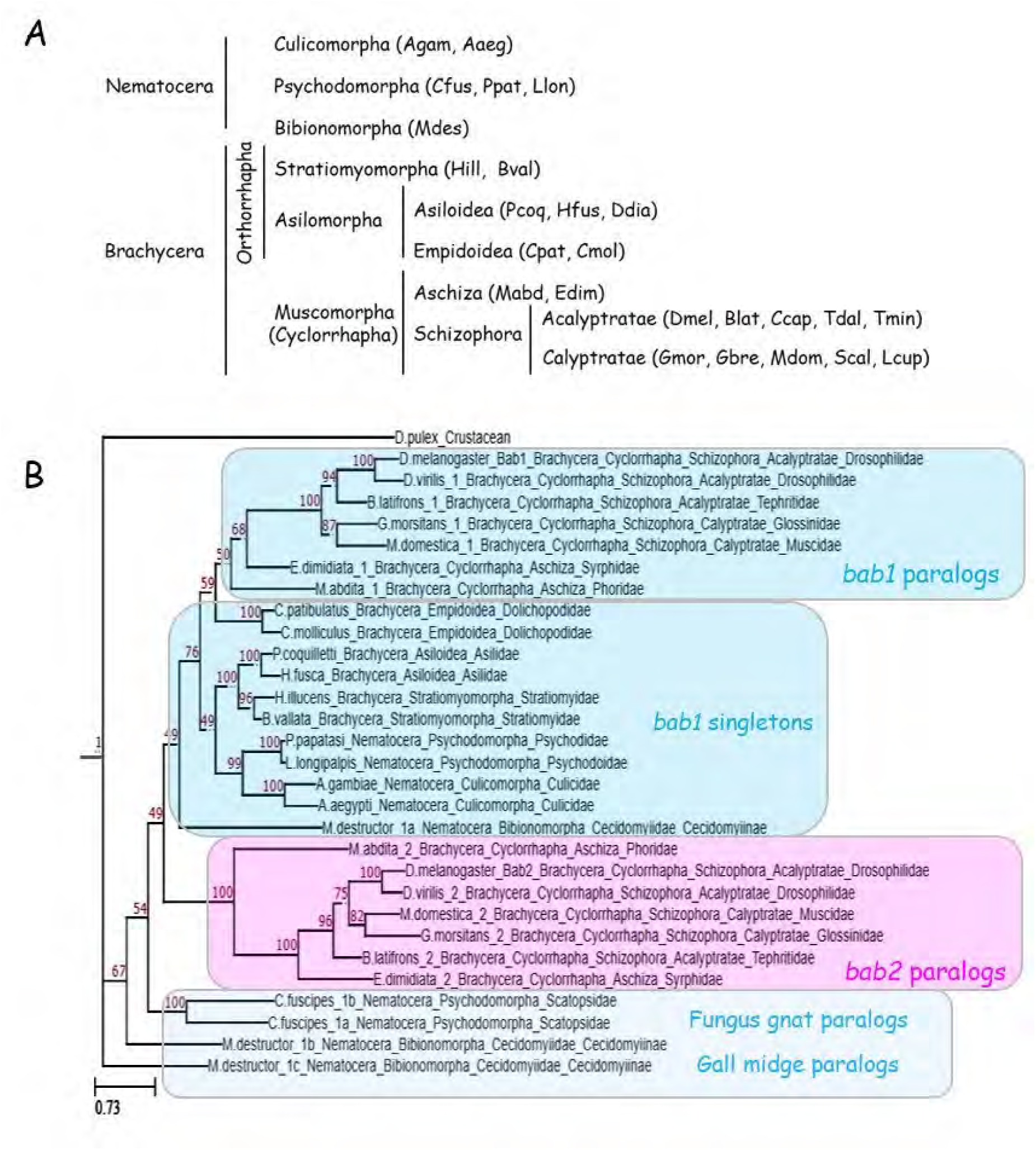
Phylogenetic relationships among dipteran *bab* paralogs and orthologs. **(A)** Dipteran families studied in this work. Species abbreviations are described in Supplementary data. **(B)** Phylogenetic relationships of the *bab* paralogs and orthologs inferred from a maximum likelihood consensus tree constructed from 1000 bootstrap replicates. Support values (percentage of replicate trees) are shown in red. Scale bar represents substitution per site. Clustered positions of *bab2* paralogs and *bab1* paralogs/orthologs are shown in pink and light blue, respectively.

To analyze the phylogenetic relationships between these different Bab-related proteins, their primary sequences were aligned and their degree of structural relatedness examined through a maximum likelihood analysis. As expected from an ancient duplication, muscomorphan Bab1-2 paralogs cluster separately, while singleton asilomorphan BTB-BabCD proteins are more related to muscomorphan Bab1 than Bab2 (Fig 8B and S5 Fig), indicating that muscomorphan *bab2* originated from *bab1* duplication.

Interestingly, contrary to most nematocerans, two or even three *bab1* paralogs are present in the fungus gnat *Coboldia fuscipes* (Psychodomorpha) and the gall midge *Mayetiola destructor* (Bibionomorpha), respectively. Significantly, *M. destructor* and *C. fuscipes bab1* paralogs (i) cluster separately in our phylogenetic analysis (Fig 8B and S5 Fig) and (ii) two are arrayed in the same chromosomal contexts for both species (S6 Fig), indicating that they have likely been generated through independent gene duplication processes in the Bibionomorpha and Psychodomorpha, respectively.

Taken together, and updating a previous work (13), our phylogenomics analysis (summarized in Fig 9B-C) indicates that a single ancestral *bab* gene related to *bab1* has been duplicated to give rise to *bab2* within the Muscomorpha (Cyclorrhapha) infraorder.

**Fig 9.**
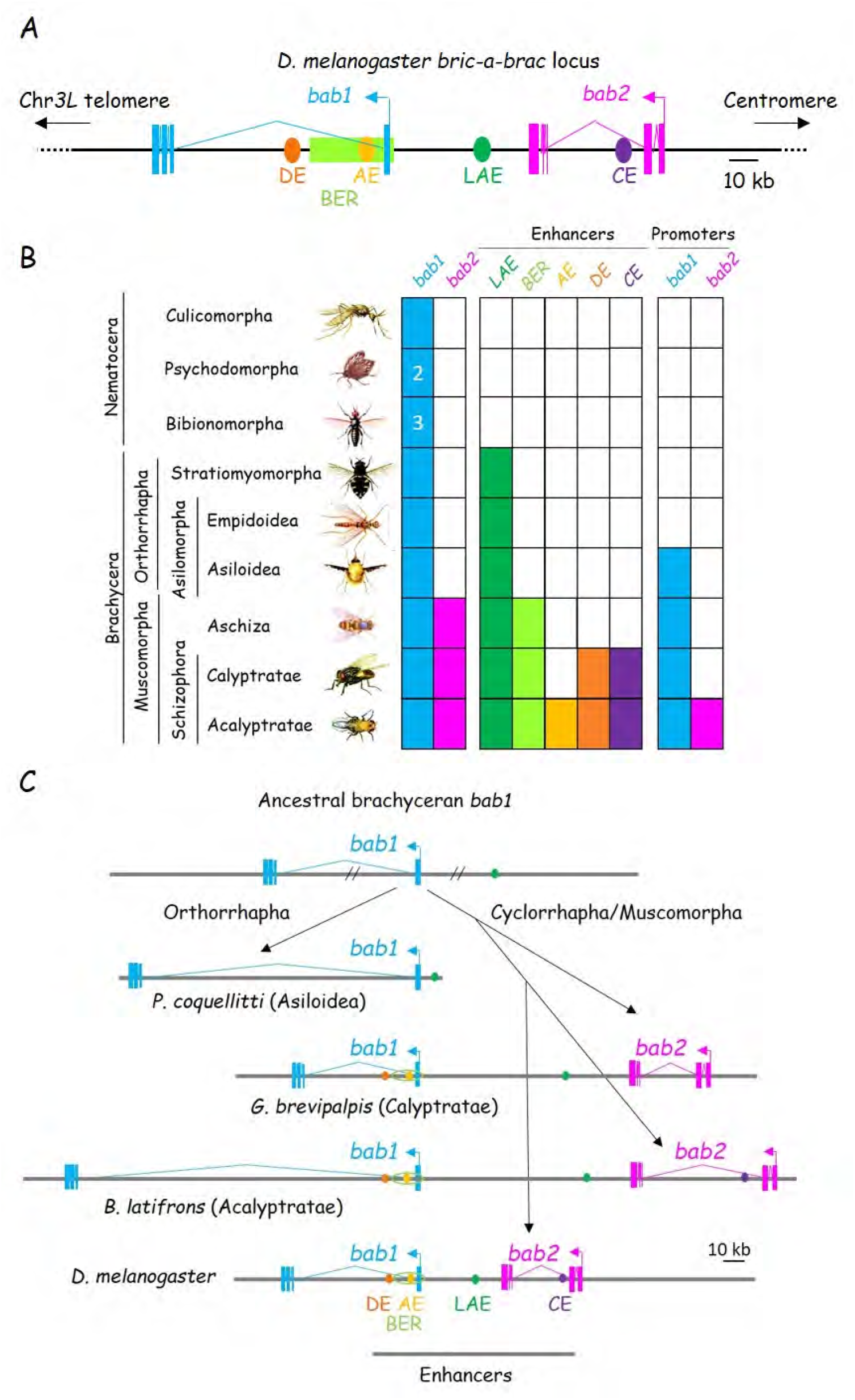
Conservation of enhancer/promoter sequences and evolutionary history of the *bab* locus among the Brachycera. **(A)** Organization of the *Dmel bab* gene paralogs and enhancers. The locus is depicted as in Fig 1A, except that *bab2* is represented in pink instead of red. **(B)** Evolutionary conservation of the *bab* gene paralogs, enhancers and promoters among diverse dipterans. Infraorders, sections, subsections and superfamilies are indicated on the left, arranged in a phylogenetic series from the “lower” Nematocera to the “higher” Brachycera suborders. Presence of *bab1* and/or *bab2* paralogs and conservation of enhancer and promoter sequences are indicated by filled or hatched boxes colored as depicted in (A). **(C)** Evolutionary scenario for the *bab* locus within the Brachycera suborders. A scheme depicting chromosomal fate of an ancestral *bab1*-like gene which gives rise to derived extant orthorrhaphan singletons (Asilomorpha) and Muscomorpha-specific paralogous (Calyptratae and Acalyptratae) genes. Locations of conserved enhancer sequences are shown, as depicted in (A).

### LAE sequences have been fixed in the Brachycera, thus predating *bab1* duplication

Having traced back the *bab* gene duplication raised the question of the evolutionary origin of the LAE enhancer, which regulates both *bab1* and *bab2* expression (17) (this work). We have previously shown that LAE includes three subsequences highly-conserved among twelve reference Drosophilidae genomes (52), termed CR1-3 (for <u>C</u>onserved <u>R</u>egions 1 to 3; see S7A Fig and Supplementary data), of which only two, CR1 and 2, are critical for tissue-specificity (17, 25). The 68 bp CR1 includes contiguous binding sites for Dll and C15 homeoproteins, while the 41 bp CR2 comprises contiguous binding sites for Dll as well as the ZF protein Bowl (S7 Fig, panels B and C, respectively) (17, 25).

To trace back the LAE evolutionary origin, we then systematically searched for homologous CR1-3 sequences (>50% identity) in dipteran genomes. Importantly, conserved LAE sequences have not been yet reported outside drosophilids. Small genomic regions with partial or extensive homologies to the CR1 (encompassing the C15 and Dll binding sites) and CR2 (particularly the Dll and Bowl binding sites) could be detected in all examined Brachycera families but not in any nematoceran (Fig 9B and S7B-C Fig). Contrary to closely-associated CR1-2 homologous sequences, no CR3-related sequence could be identified nearby, in any non-Drosophilidae species. Significantly, homologous LAE sequences are situated (i) in between the tandemly-duplicated paralogs in muscomorphan species for which the entire *bab* locus sequence was available to us, suggesting an evolutionarily-conserved enhancer role, or (ii) 20 kb upstream of the *bab1*-related singleton in the asilomorphan *P. coquilletti* (see Fig 9C).

Taken together, as summarized in Fig 9A-C, these data suggest that a LAE-like enhancer with CR1- and CR2-related elements emerged early on in the Brachycera suborder, 180-200 million years ago, and has been since fixed within or upstream the *bab* locus in the Muscomorpha and Asilomorpha infraorders, respectively.

### Like LAE elements, *bab1* promoter sequences have been fixed early on in the Brachycera

Given their differential interplay with the long-lasting LAE enhancer, we next analyzed the evolutionary conservation of *Dmel bab1*-*2* promoter core sequences (Fig 9B and S8 Fig). Both *bab* promoters are TATA-less. Whereas *bab1* has a single transcriptional initiator (Inr) element (TTC<u>A</u>GTC), its *bab2* paralog displays tandemly-duplicated Inr sequences (ATTC<u>A</u>GTTCGT) (53, 54) (S8 Fig). Both promoters display 64% sequence identity over 28 base pairs, including Inr (TTC<u>A</u>GT) and downstream putative <u>P</u>ause <u>B</u>utton (PB; consensus CGNNCG) sequences (55) (see S8A Fig). These data suggested that (i) the duplication process having yielded *bab2* included the ancestral *bab1* promoter and (ii) PolII pausing ability previously shown for *bab2* promoter (56–58) probably also occurs for *bab1* promoter.

Homology searches revealed that *bab1* promoter sequences have been strongly conserved in the three extant Muscomorpha families and even partially in some asilomorphans (e.g., *P. coquellitti*), for which a *bab1*-related singleton gene is present (Fig 9B and S8B Fig). In striking contrast to *bab1*, sequence conservation of the *bab2* promoter could only be detected among some Acalyptratae drosophilids (Fig 9B and S8C Fig). In agreement with a fast-evolutionary drift for *bab2* promoter sequences, the duplicated Inr is even only detected in Drosophila group species.

Taken together, these evolutionary data (summarized in Fig 9B) indicate that, likewise for the LAE enhancer, *bab1* promoter sequences have been under strong selective pressure among the Brachycera, both in the Muscomorpha and Asilomorpha infraorders, while paralogous *bab2* promoter sequences diverged rapidly among muscomorphans.

### Unlike LAE, other *bab* CRE sequences have not been conserved beyond the Muscomorpha

The broad LAE sequence conservation led us to also trace back the evolutionary origins of the pleiotropic BER enhancer region as well as the cardiac CE, abdominal anterior AE and sexually-dimorphic DE *cis*-regulatory elements (see Fig 9). Sequences homologous to half of the BER OCS subregions could be detected among the 12 reference Drosophilidae genomes (52), in Calyptratae schizophorans and even in the Muscomorpha Aschiza subsection (e.g., OCS3) (S10-18 Fig). Unlike the LAE enhancer, homologous BER sequence elements (except the *bab1* promoter) could not be detected in non-muscomorphan families. Cardiac CE and abdominal DE are even less conserved given that related sequences could be only detected within schizophoran (excepted in Calyptratae) (Fig 9B and S9 Fig, panels B and C, respectively), whereas abdominal AE sequences could be only identified among drosophilids (Supplementary data) but not in aschizan, asilomorphan and nematoceran *bab* loci.

In conclusion, contrary to the LAE enhancer which among the Diptera emerged early on in the Brachycera suborder, other so-far identified *bab cis*-regulatory sequences have not been conserved beyond the Muscomorpha infraorder. Thus, as summarized in Fig 9B, and unlike the long-lasting brachyceran LAE (CR1-2) sequences, these data suggest that other enhancer sequences have been fixed within the Muscomorpha concomitantly (BER) or even after (CE, DE and AE) the *bab2* paralog emergence. Moreover, as expected for a pleiotropic enhancer region, BER sequence conservation allowed us to predict binding sites for transcription factors known, or so far unsuspected, prone to regulate directly the two *bab* genes in many distinct developmental contexts, and which are presented hereafter.

### Predictive TF combinatorial code governing *bab* gene expression in diverse tissues

We gathered our data from TF binding site evolutive conservation (described in Supplementary data and S10-19 Fig) with ChIP-Seq experiments from the literature (GEO datasets; see Materials and Methods) (Fig 5B and S3B Fig). Associated with our precise knowledge of *bab* locus enhancer sequences and with previous genetic data also gained from the literature, our compilation presented in Fig. 10 allows to propose new models for the TF code involved in *Dmel bab* locus regulation: (i) It provides new insights into limb-specific *bab1*/*2* regulation proposing additional direct regulators such as Sp1 in the legs, Hth in the antenna, Scalloped (Sd) in the wing and Ubx in the haltere (ChIP-Seq data shown in S3 Fig); (ii) It suggests that BER also acts as an enhancer region for *bab* gene regulation in other developmental contexts and that common TF sets (notably Abd-B together with Dsx) are acting through distinct *cis*-regulatory elements within BER to drive *bab* gene expression in distinct tissues (e.g., in abdominal histoblast nests versus genitalia); (iii) It proposes a direct Bab2 binding on LAE for *bab* gene auto- and cross regulation (tested above); (iv) Finally, analysis of sequences conserved among brachyceran *bab* loci identified predicted binding sites for more broadly-expressed transcriptional regulators, i.e., GAF, Pho and CTCF (directly interacting with BER OCS; see Fig 5B, lanes 3, 14 and 19, respectively, as well as S3 Fig for GAF), as well as Eip93F, Eip74EF, Chinmo, all related to chromatin organization whose putative roles in *bab* locus regulation are discussed hereafter.

**Fig 10.**
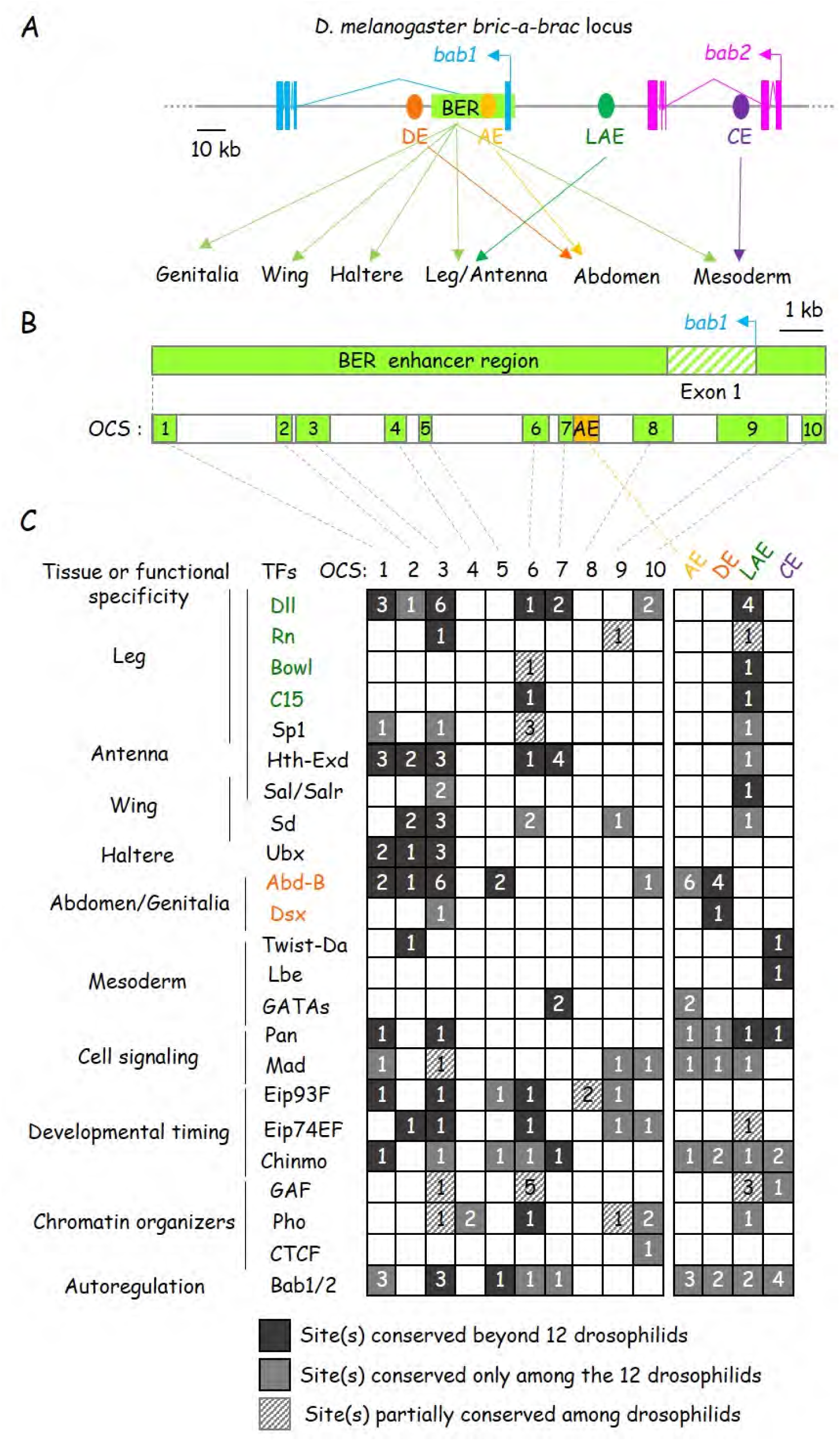
A comprehensive predictive tissue-specific TF code governing *bab* paralog expression. **(A)** Organization of the *Dmel bab* gene paralogs and enhancers, as depicted in Fig 9A. Expression tissue specificities conferred by each enhancer are shown in beneath. **(B)** BER structural and chromatin state organizations. The *bab1* first exon is hatched. OCS regions (see Fig 6C), as defined in leg tissues; are represented by light green boxes. The abdominal AE CRE (not detected in FAIRE-Seq data from leg and eye-antennal discs) is depicted as a light orange box. **(C)** Evolutive conservation of predicted TF binding sites within *Dmel bab cis*-regulatory sequences. Site conservation among and beyond drosophilids of transcriptional regulators involved in tissue-specific morphogenetic processes, cell signaling, developmental timing and chromatin organization. Predicted/validated TF site numbers are indicated within colored or hatched boxes reflecting their relative conservation are indicated in beneath. Experimentally-validated direct *bab* regulators are colored according to their well characterized bound-enhancer sequences (see A).

## Discussion

In this work, we have addressed the issue of the emergence and functional diversification of enhancers and promoters from two tandem gene duplicates. Using the *Drosophila bab* locus as a model, we showed that the paralogous genes *bab1*-*2* originate from an ancient *bab1* duplication in the Muscomorpha/Cyclorrhapha. The early-fixed brachyceran *bab1* LAE has been co-opted lately to regulate also *bab2* expression. Furthermore, this unique enhancer is also responsible for paralog-specific *bab2* expression along the P-D leg axis presumably through privileged interactions with the *bab2* promoter. Finally, LAE regulates only some aspects of *bab1-2* expression in the developing limbs because redundant information has emerged within a large pleiotropic enhancer driving *bab1* and/or *bab2* expression in highly-diverse tissues, by binding common sets of developmental transcription factors. This work brings some cues about (i) how a single enhancer can drive specificity among tandem gene duplicates, (ii) how enhancers evolutionary adapt with distinct cognate gene promoters, and (iii) which functional links can be predicted between dedicated transcription factors and chromatin dynamics during development.

### A shared enhancer differentially regulating two tandem gene paralogs through distinct promoter targeting specificities

Here, we have shown that a single enhancer, LAE, regulates two tandem gene paralogs at the same stage and in the same expression pattern. How can this work? It has been proposed that enhancers and their cognate promoters are physically associated within phase-separated nuclear foci composed of high concentrations of TFs and proteins from the basal RNA polymerase II (PolII) initiation machinery inducing strong transcriptional responses (59, 60). Our Hi-C data from eye-antennal discs show a strong interaction between *bab1*-*2* promoter regions (Fig 5), suggesting that both *bab* promoters could be in close proximity within such phase separated droplets, thus taking advantage of shared transcriptional regulators and allowing concerted gene regulation. In contrast, no strong chromosome contacts could be detected between LAE and any of the two *bab* promoter regions, indicating that this enhancer is not stably associated to the *bab2* or *bab1* promoter in the eye-antennal disc (where only the antennal distal part expresses both genes). It would be interesting to gain Hi-C data from leg discs, in which the *bab1*-*2* genes are much more broadly expressed.

In addition to be required for *bab1*-*2* co-expression in proximal tarsal segments, we showed here that the LAE enhancer is also responsible for paralog*-*specific *bab2* expression along the proximo-distal leg axis. While it has been proposed that expression pattern modifications occur through enhancer emergence, our present work indicates that differential expression of two tandem gene paralogs can depend on a shared pre-existing enhancer (i.e., LAE). How this may work? Relative to its *bab1* paralog, *bab2* expression extends more proximally within the Dac-expressing ts1 cells (44) and more distally in the ts5 segment expressing nuclear Bowl protein, whereas both Dac and Bowl proteins have been proposed to act as *bab2* (and presumably *bab1*) repressors (25, 33, 61). CRISPR/Cas9-mediated LAE excision allowed us to establish that this enhancer is critically required for paralog-specific *bab2* leg expression proximally and distally, in ts1 and ts5 cells, respectively. In this context, we and others have previously proposed that transiently-expressed Rotund activating TF may antagonize Bowl (and eventually Dac) repressive activity to precisely delimit *bab2* expression among ts1 cells (17, 61). Given that *bab1-2* are distinctly expressed despite being both regulated by Bowl and Rotund, we propose that paralog-specific LAE activity depends on privileged interactions with *bab2* promoter sequences (discussed below). Thus, we speculate that the *bab2* promoter responds to Rotund transcriptional activity differently from its *bab1* counterpart. Consistent with this view, ectopic Rotund expression reveals differential regulatory impacts on the two *bab* gene promoters (S1B Fig). Genetic together with Hi-C experiments indicate that this could occur through specific interactions between LAE-bound TFs (e.g., Rotund) and dedicated proteins within the PolII pre-initiation complex stably-associated to the *bab2* core promoter. We envision that the LAE-bound ZF protein Rotund, the chromatin-remodeling ZF protein GAF (for <u>GA</u>GA-associated <u>F</u>actor) and the PolII-associated TFIID subunit TAF3, the latter known to interact physically with GAF and Bab2 BTB proteins (62, 63), are parts of the underlying promoter targeting molecular mechanism.

In this context, despite that sequence homologies between both promoters (consistent with an ancient duplication event mobilizing the ancestral *bab1* promoter) are still detectable, it is significant that the *bab2* promoter evolves much faster than its *bab1* counterpart. While the *bab1* promoter sequence has been strongly conserved among brachycerans, predating *bab2* gene emergence in the Muscomorpha, the *bab2* promoter sequence has only been fixed recently among Drosophilidae, notably through the Initiator (Inr) sequence duplication, indicating very fast evolutionary drift after the gene duplication process which yielded the *bab2* paralog. We envision that this evolutionary ability has largely contributed to allow novel expression patterns for *bab2*, presumably through differential enhancer-promoter pairwise interplay.

### Differential evolutionary conservation of tissue-specific versus pleiotropic enhancers

Our comprehensive phylogenomics analyses from highly diverse Diptera families indicate that the *bab* gene complex has been generated through tandem duplication from an ancestral *bab1*-related gene singleton within the Muscomorpha (Cyclorrhapha), about 100-140 years ago. This result contrasts with published data reporting that the duplication process having yielded the tandem *bab* genes occurred much earlier in the Diptera lineage leading to both the Brachycera (true flies; i.e., with short antenna) and Nematocera (long horned “flies”, including mosquitos) suborders (13). In fact, tandem duplication events implicating the *bab* locus did occur in the Bibionomorpha, as reported (13)), and even in the Psychodomorpha with three *bab1*-related gene copies (Figure 8 and S6 Fig), but our phylogenetic analysis supports independent events. Thus, within the Diptera, the ancestral *bab1* singleton had a high propensity to duplicate locally.

In this study, we have shown a strong evolutionary conservation of LAE subsequences among brachycerans, notably its CR2 element containing Dll and Bowl binding sites (S7C Fig). This conservation suggests a long-lasting enhancer function in distal limb-specific regulation of ancestral singleton *bab1* genes. In striking contrast, BER sequence conservation could only be detected among extant muscomorphan *bab* loci. We assume that during evolution large pleiotropic enhancers may better assimilate binding sites for gene-specific transcription factors, thus generating regulatory novelties in distinct imaginal discs.

Gene duplication is a major source to generate phenotypic innovations during evolution, through diverging expression and molecular functions, and eventually from single gene copy translocation to another chromosomal site. Emergence of tissue-specific enhancers not shared between the two gene duplicates, as well as of “shadow” enhancers, have been proposed to be evolutionary novelty sources (64) (6). Our work indicates that the presumably long-lasting brachyceran LAE enhancer has recently been co-opted in drosophilids to allow differential *bab* gene expression. Conversely, the large BER region has apparently accumulated regulatory sequence elements bound by diverse tissue-specific transcription factors (e.g., Dll, Hth, Abd-B and Dsx) acting in different cellular contexts.

### A pleiotropic enhancer region overlapping with a PcG-response element

ChIP-Seq analysis for histone H3 modifications (H3K4me3, H3K27Ac and H3K4me1 enhancer/promoter marks; H3K27me3 chromatin repressive mark) from eye-antennal discs has revealed the pleiotropic BER enhancer region but also an overlapping repressive PcG (<u>P</u>oly<u>c</u>omb <u>G</u>roup family) domain, indicating that BER encompasses a bivalent chromatin domain, while another one is detected over the *bab2* promoter region. A dual enhancer/silencing function for <u>P</u>cG-<u>R</u>esponse <u>E</u>lement (PRE) during embryogenesis has recently been established genome-wide (65), and the authors have proposed that reuse of enhancer regulatory elements by PcG proteins may help fine-tune gene expression and ensure the timely maintenance of cell identities throughout development. More recently, we have shown widespread enhancer-PcG domain interplay during developmental gene activation through chromatin looping in eye-antennal discs (38). Altogether these data suggest that the *bab1-2* genes might be poised for activation throughout the eye-antennal disc, and possibly other imaginal discs as well.

The Pleiohomeotic (Pho) protein is a DNA-binding PcG complex recruiter, critical for gene silencing maintenance during development (66). Pho interaction with several BER subregions, as detected in ChIP-Seq experiments from L3 tissues, as well as the presence of many evolutionarily-conserved predicted Pho binding sites, support a role for Pho in PcG repression throughout BER. We thus propose that Pho-containing PcG repressive complex bound at PREs within the *bab* bivalent locus is counteracted by one or several tissue-specific transcriptional activators identified in this work, which remain(s) to be characterized.

In this context, it is significant that in the eye-antennal disc, the ZF protein CTCF, acting redundantly with other chromatin insulator proteins, strongly interacts directly with the two flanking regions of the TAD covering the *bab* locus and also with several BER OCSs (Fig 5B). Significantly, two of these predicted CTCF interacting sites overlap with putative optimal binding sites for the PcG-recruiter Pho (S18-19 Fig). These data suggest that the CTCF architectural protein and the PcG-recruiter Pho may functionally interact to regulate the *bab* locus chromosome topology. Interestingly, the human Pho homolog (YY1) is a structural enhancer-promoter looping regulator (67) and orchestrates, together with the CTCF protein, a 3D chromatin looping switch during early neural lineage commitment (68). To our knowledge, functional relationships between CTCF and Pho proteins have not been investigated genome-wide in *Drosophila*.

### Dynamic *bric-a-brac* locus chromatin accessibility during development

Recent data indicate that chromatin accessibility is dynamic during *Drosophila* larval development, being triggered by the ecdysone hormone (69). Dynamic enhancer activity and chromatin accessibility have been proposed to be regulated by the ecdysone-induced Eip93F (<u>E</u>cdysone-induced <u>p</u>rotein 93F, also called E93) transcriptional regulator (70). ChIP data from early pupal wings indicate that Eip93F binds many BER OCS as well as the *bab1* promoter region (70). Consistently, many putative Eip93F binding sites are present in these BER subregions and several have been conserved beyond Drosophilidae (Fig 10C). Interestingly, the human Eip93F homolog interacts with CtBP through a conserved motif (71), and *Drosophila* CtBP is known to recruit diverse chromatin-modifying complexes, notably to participate in Pho-mediated PcG recruitment to PREs (72). Thus, Eip93F binding to BER *cis*-regulatory elements may impact the proposed dual PcG activity at the *bab* locus. In addition to Eip93F, BER regulatory sequences include many evolutionarily-conserved putative binding sites for the Eip74EF protein (Fig 10C), another ecdysone-induced TF, including one which overlaps with a conserved putative Pho binding site, suggesting again functional correlation between Ecdysone regulation and PcG activity.

Lastly, the *cis*-regulatory landscape within the *bab* locus (i.e., AE, DE, CE, LAE and BER) includes one or several evolutionarily-conserved predicted binding sites for the Chinmo BTB-ZF protein participating to developmental timing, notably through interplay with ecdysone signaling (73, 74). Consistently, ChIP-Seq experiments from embryos (ModENCODE data; http://www.modencode.org/) indicate Chinmo binding to BER sequences (75). Intriguingly, the Chinmo ZF protein is an additional BTB-containing TF prone to regulate directly the *bab* genes, possibly through molecular partnerships with the chromatin organizer GAF (another BTB-ZF protein interacting directly with both *bab* promoter regions; Fig 5B and S3 Fig) and the twin Bab BTB-BabCD proteins themselves. Thus, Chinmo implication in chromatin organization and enhancer activity within the *bab* locus undoubtedly deserves to be investigated.

In summary, the *bab* locus offers a good paradigm to investigate molecularly in great details how chromatin structure, particularly higher-order chromosome organization, impacts on transcriptional memory during development and selective enhancer-promoter interplay in diverse tissular contexts. Indeed, our comprehensive predictive combinatorial code for tissue-specific, as well as broadly-expressed architectural transcription factors (e.g., CTCF, Pho and GAF) regulating two tandem gene paralogs, offers the opportunity to dissect underlying molecular mechanisms, which are prone to be conserved during animal evolution and thus to be of broad biological significance.

## Material sand Methods

### Fly stocks, culture and genetic manipulations

*D. melanogaster* stocks were grown on standard yeast extract-sucrose medium. The *vasa*-PhiC31 ZH2A *attP* stock (kindly provided by F. Karch) was used to generate the *LAEpHsp70-GFP* reporter lines and the *BAC69B22* construct as previously described (17). *LAE-GFP* and *LAE-RFP* constructs (including both *Hsp70* and *bab2* core promoters) inserted on the ZH2A (X chromosome) or ZH86Fb (third chromosome) *attP* landing platforms, and displaying identical expression patterns, have been previously described (17, 25). *C15^2^*/TM6B, *Tb^1^* stock was kindly obtained from G. Campbell. Mutant mitotic clones for null alleles of *bowl* and *rotund* were generated with the following genotypes: *y w LAE-GFP*; *DllGal4^EM2012^*, *UAS*-*Flp/+; FRT82B, Ub-RFP/FRT82B rn^12^* (i.e., *rn* mutant clones are RFP negative; Fig 1E) and *y w LAE-RFP*; *DllGal4^EM2012^*, *UAS*-*Flp/+*; *Ub-GFP, FRT40A*/*bowl^1^ FRT40A* (i.e., *bowl* mutant clones are GFP negative; Fig 1F), respectively. Rotund protein gain-of-function within the *Dll*-expressing domain was obtained with the following genotype: *y w LAE-GFP*; *DllGal4^EM2012^*; *UAS-Rn^1^*/+. The *Dll^EM212^-Gal4* line was provided by M. Suzanne, while the *UAS-Rn^1^* line was obtained from the Bloomington stock center. “Flip-out” (FO) mitotic clones over-expressing dsRNA against *lines* were generated by 40 mn heat shocks at 38°C, in mid-late L2 to early-mid L3 larvae of genotypes: *y w LAE-RFP hsFlp*; *UAS-dsRNAlines/pAct>y+>Gal4, UAS-GFP (*i.e., FO clones express GFP in S1C Fig). Mutant mitotic clones for the null *bab^AR07^* allele were generated by 30 mn heat shocks at 38°C, in early first to late second-instar larvae of genotypes: y w *LAE-GFP*, hsFlp; FRT80B/*bab^AR07^*, FRT80B. FO mitotic clones over-expressing dsRNA against *bab1* or *bab2* were generated by 40 mn heat shocks at 38°C, in early to early-mid L3 larvae of genotypes: *y w LAE-RFP hsFlp*; *UAS-dsbab2 /Pact>y+>Gal4, UAS-GFP* and *y w LAE-RFP, hsFlp*; *Pact>y+>Gal4*, *UAS-GFP*/+; *UAS-dsbab1*/+ (i.e., FO clones express GFP in Fig 7). *UAS-dsRNA* stocks used to obtain interfering RNA against *lines* (#40939), *bab1* (#35707) and *bab2* (#37260) were obtained from the Bloomington stock Center.

### Immuno-histochemistry and microscopy

Leg discs were dissected from wandering (late third instar stage) larvae (L3). Indirect immuno-fluorescence was carried out as previously described (17) using a LEICA TCS SP5 or SPE confocal microscope. Rat anti-Bab2 (11), rabbit anti-Bab1 (14), rabbit ant-Dll (76), rabbit anti-Bowl (61), and rabbit anti-C15 (31) antibodies were used at 1/2000, 1/500, 1/200, 1/1000 and 1/200, respectively.

### CRISPR/Cas9-mediated chromosomal deletion

Guide RNAs (gRNAs) were designed with CHOPCHOP at the Harvard University website (https://chopchop.cbu.uib.no/). Four gRNA couples were selected that cover two distinct upstream and downstream LAE positions: TGCGTGGAGCCTTCTTCGCC<u>AGG</u> or TGGAGCCTTCTTCGCCAGGC<u>CGG</u>; and TATACTGTTGAGATCCCATG<u>CGG</u> or TTAGGCGCACATAAGGAGGC<u>AGG</u> (the PAM protospacer adjacent motif sequences are underlined), respectively. Targeting tandem chimeric RNAs were produced from annealed oligonucleotides inserted into the pCFD4 plasmid, as described in (http://www.crisprflydesign.org/). Each pCFD4-LAE-KO construct was injected into 50 *Vasa-Cas9* embryos (of note the *vasa* promoter sequence is weakly expressed in somatic cells). F0 fertile adults and their F1 progeny, with possible somatic LAE-deletion events and candidate mutant chromosomes (balanced with *TM6B, Tb*), respectively, were tested by polymerase chain reactions (PCR) with the following oligonucleotides: AGTTTTTCATCCCCCTTCCA and GTATTTCTTTGCCTTGCCATCG (predicted wild-type amplified DNA: 2167 base pairs).

### Quantitative RT-PCR analysis

T1-3 leg imaginal discs were dissected from homozygous *white^1118^* and *bab^ΔLAE-M1^* late L3 larvae in PBS 0.1% Tween. 50 discs of each genotype were collected and frozen in nitrogen. Total messenger RNAs were purified using RNeasy kit (Qiagen) and reverse transcribed by SuperScript II (ThermoFisher). *bab1*, *bab2* or *rp49* cDNA levels were monitored by quantitative PCR using the following oligonucleotides: Bab1Fw: CGCCCAAGAGTAACAGAAGC; Bab1Rev: TCTCCTTGTCCTCGTCCTTG; Bab2Fw: CTGCAGGATCCAAGTGAGGT; Bab2Rev: GACTTCACCAGCTCCGTTTC; RP49Fw: GACGCTTCAAGGGACAGTATCTG; RP49Rev: AAACGCGGTTCTGCATGAG. A Wilcoxon test was performed to evaluate the difference between samples.

### Homology searches, sequence alignments and phylogenetic analyses

Homology searches were done at the NCBI Blast site (https://blast.ncbi.nlm.nih.gov/Blast.cgi). Protein or nucleotide sequence alignments were done using MAFFT (Multiple Alignment using Fast Fourier Transform) (https://mafft.cbrc.jp/alignment/server/). Phylogenetic relationships were inferred through a maximum likelihood analysis with W-IQ-Tree (http://iqtree.cibiv.univie.ac.at/) and visualized with the ETE toolkit (http://etetoolkit.org/treeview/).

### Transcription factor binding prediction

DNA binding predictions were done using the motif-based sequence analysis tool TomTom from the MEME suite (http://meme-suite.org/tools/tomtom) and the Fly Factor Survey database (http://mccb.umassmed.edu/ffs/).

### Gene expression omnibus datasets

The following gene expression omnibus (GEO) datasets were extracted from the NCBI website (https://www.ncbi.nlm.nih.gov/gds/): GSE59078; GSM1261348; GSM1426265; GSE126985; GSM3139658; GSM948715; GSE113574; GSM948718; GSM948717; GSE38594; GSM948720; GSM948716; GSM659162; GSM948719; GSE102339; GSE50363.

### Hi-C and histone tail mark analyses from L3 eye-antennal imaginal discs

Hi-C and histone mark ChIP-Seq analyses from L3 eye-antennal discs have been recently published in (38).

## Acknowledgments

We thank F. Karch, M. Suzanne, T.M. Williams, G. Boekhoff-Falk, S. Bray, G. Campbell, T. Kojima, F. Laski, J.L. Couderc and the Bloomington Stock Center for fly stocks and reagents. We are grateful to Alain Vincent for his proofreading of the manuscript. We thank Julien Favier for technical assistance, particularly in managing the transgenic facility. Lastly, we acknowledge Brice Ronsin and the Toulouse RIO Imaging platform.

---

**S1 Figure.**
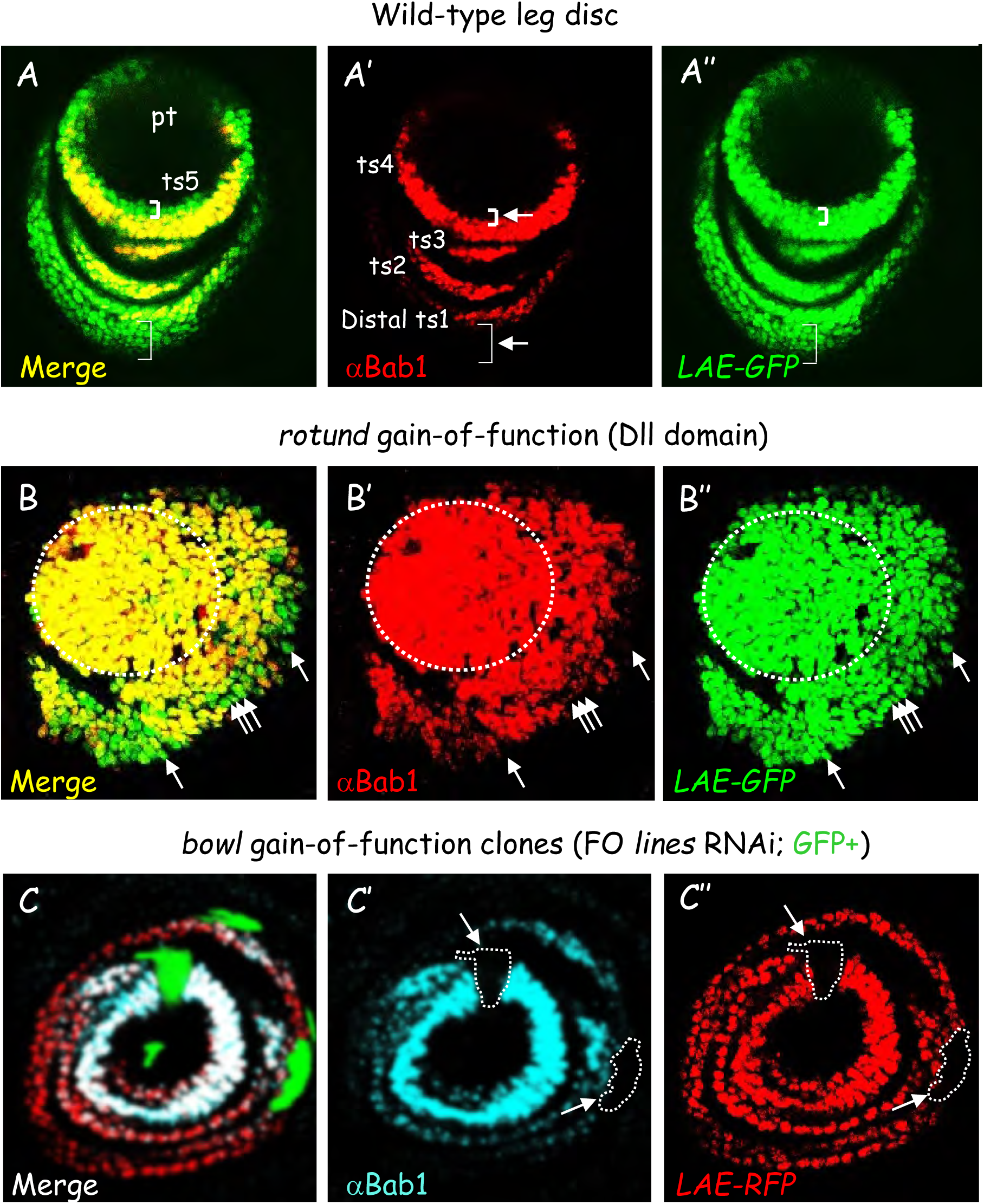

**S2 Figure.**
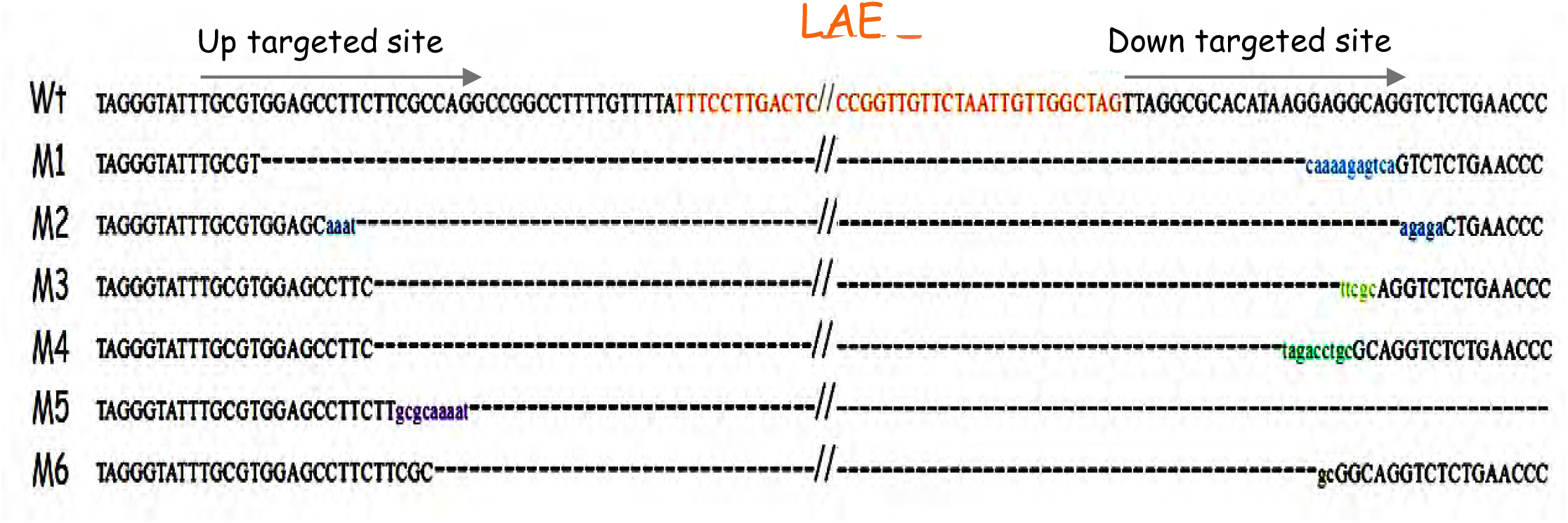

**S3 Figure.**
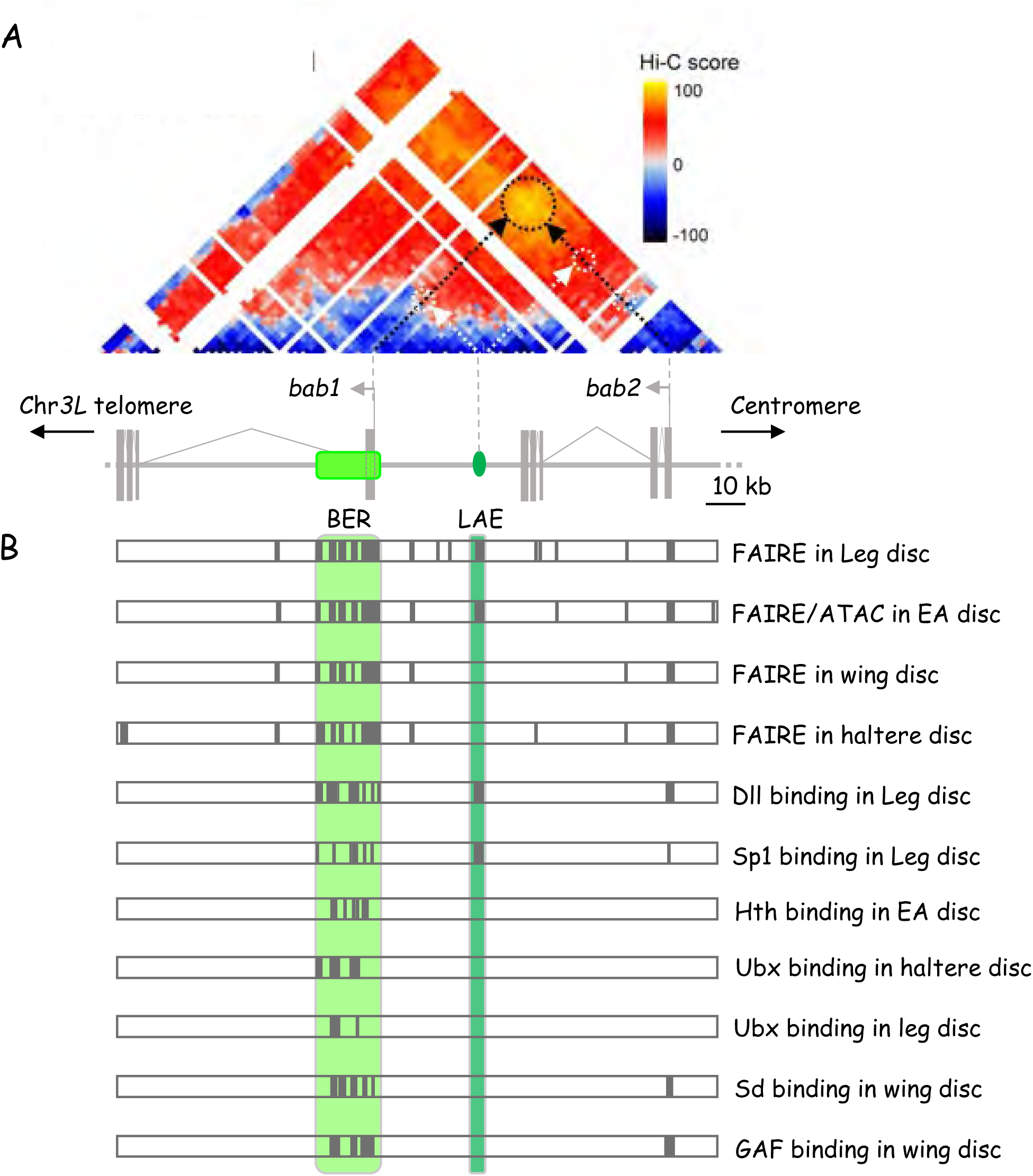

**S4 Figure.**
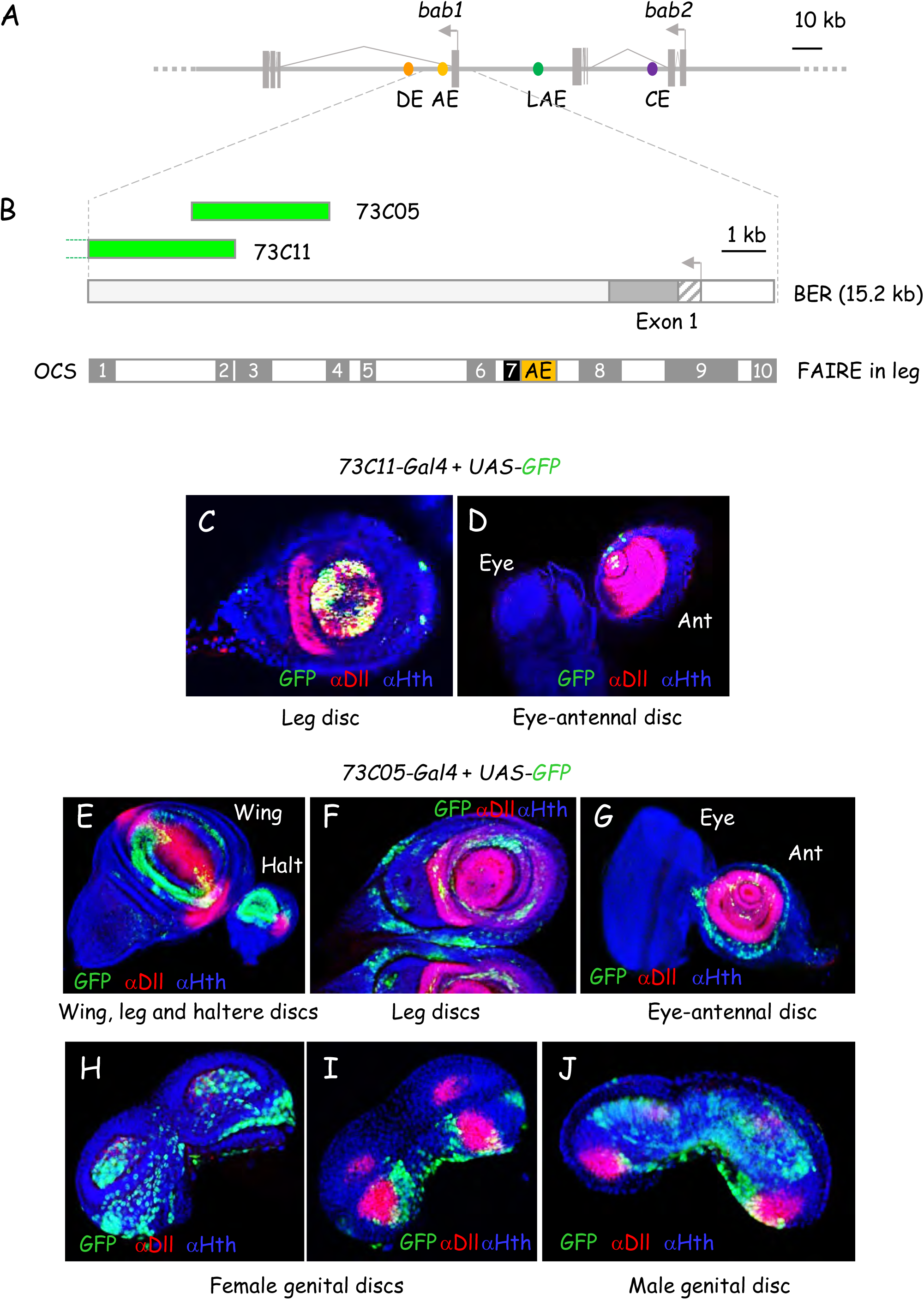

**S5 Figure.**
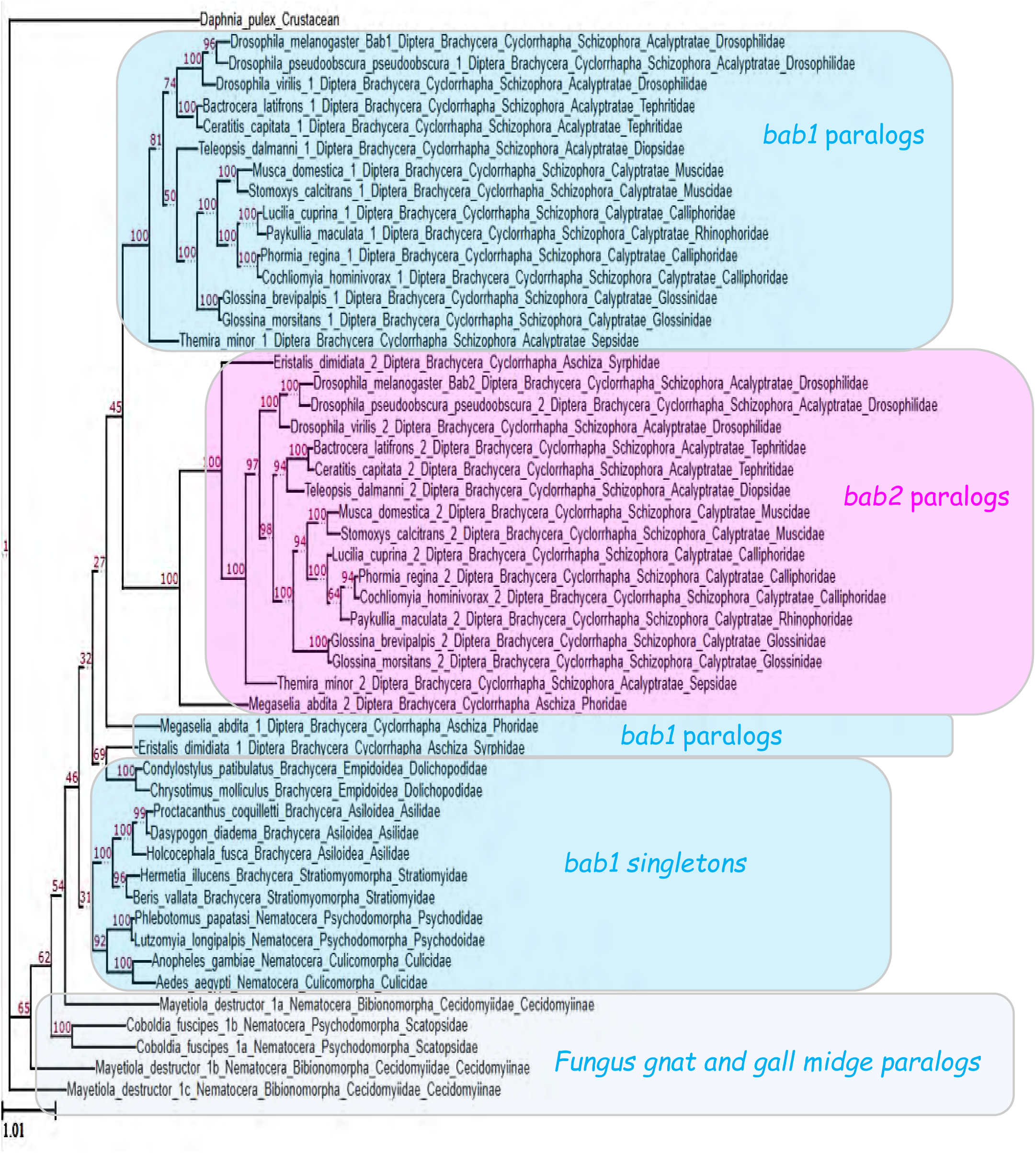

**S6 Figure.**
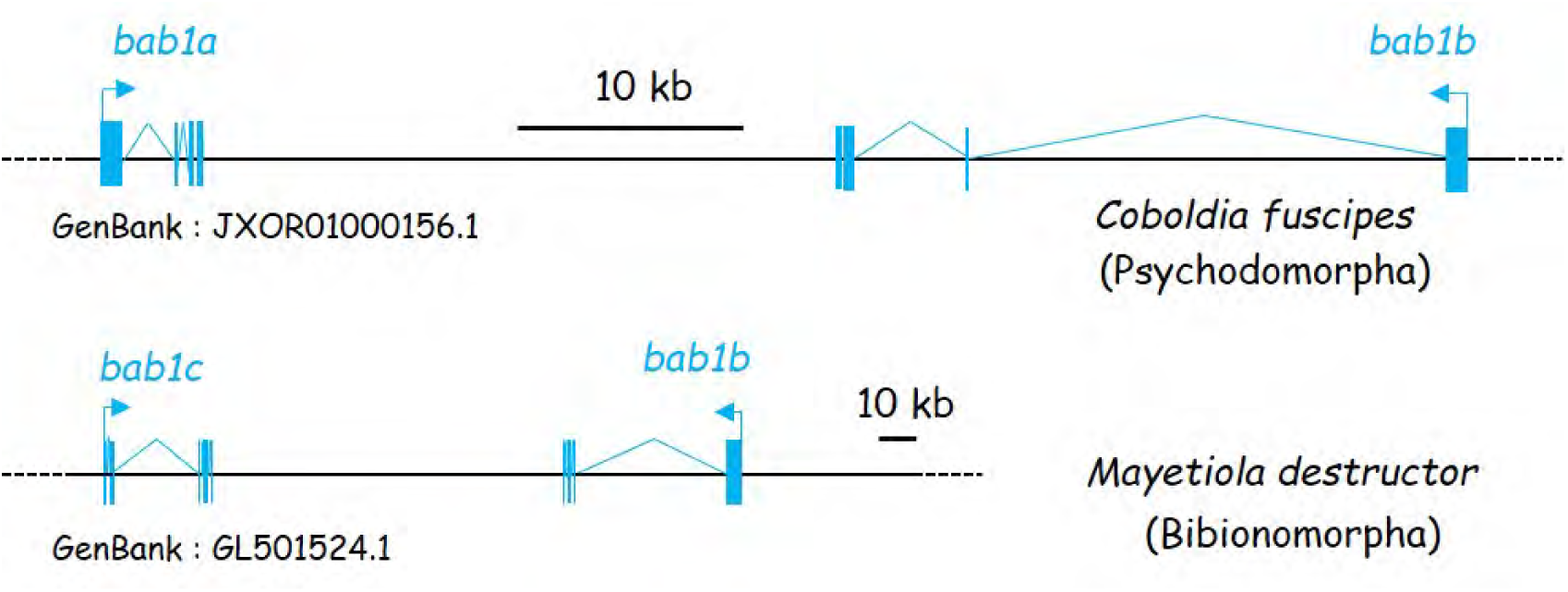

**S7 Figure.**
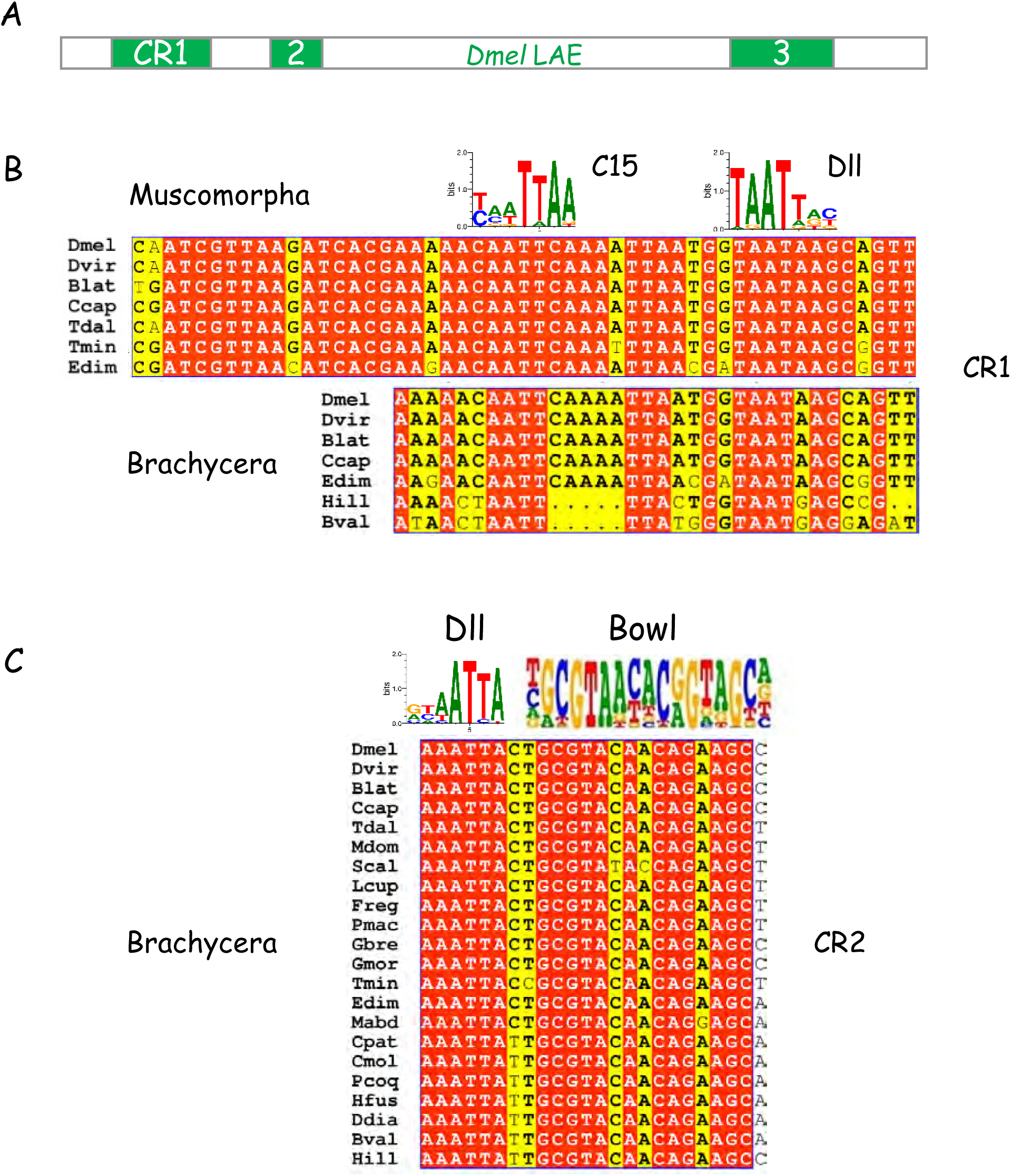

**S8 Figure.**
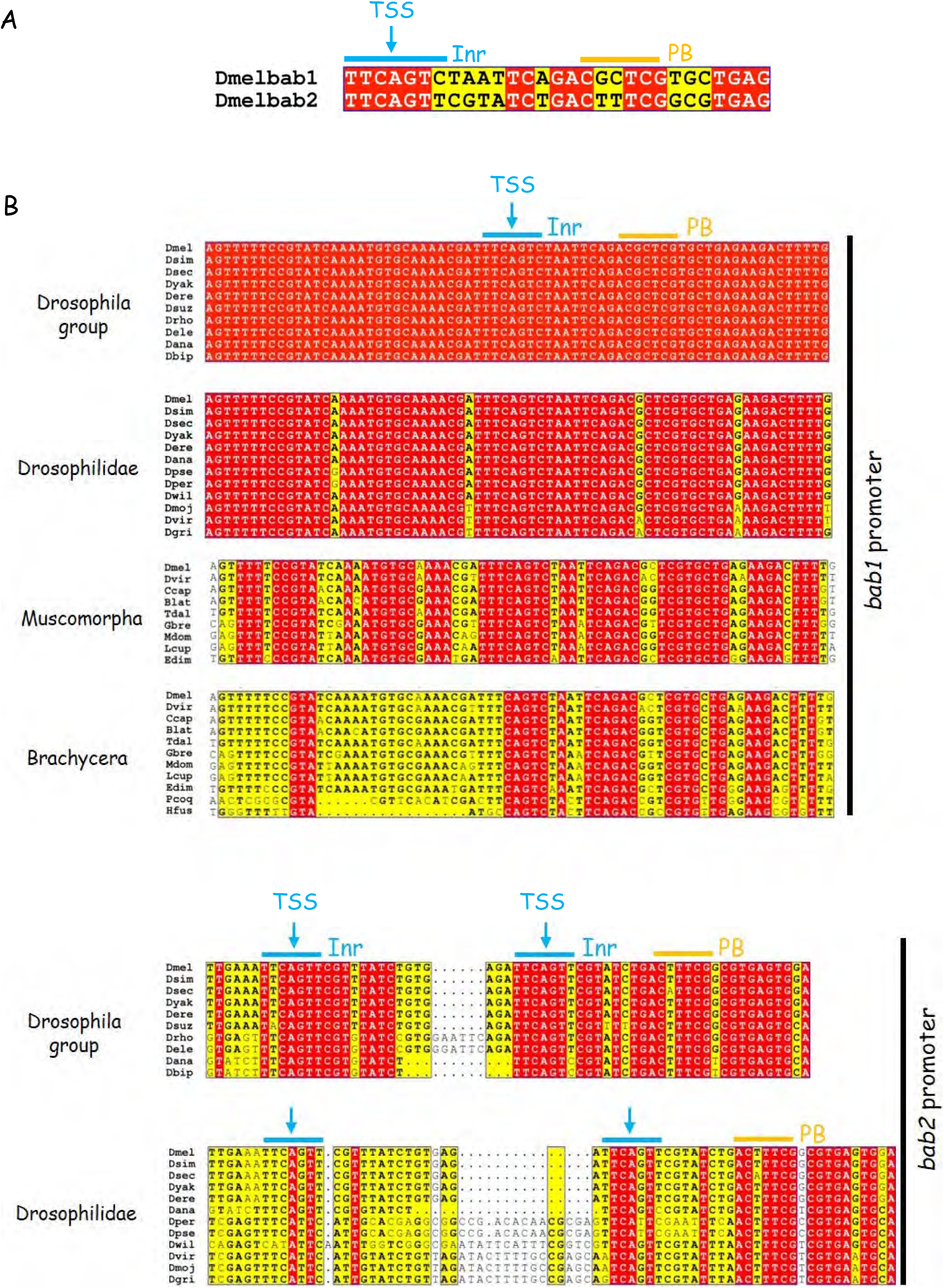

**S9 Figure.**
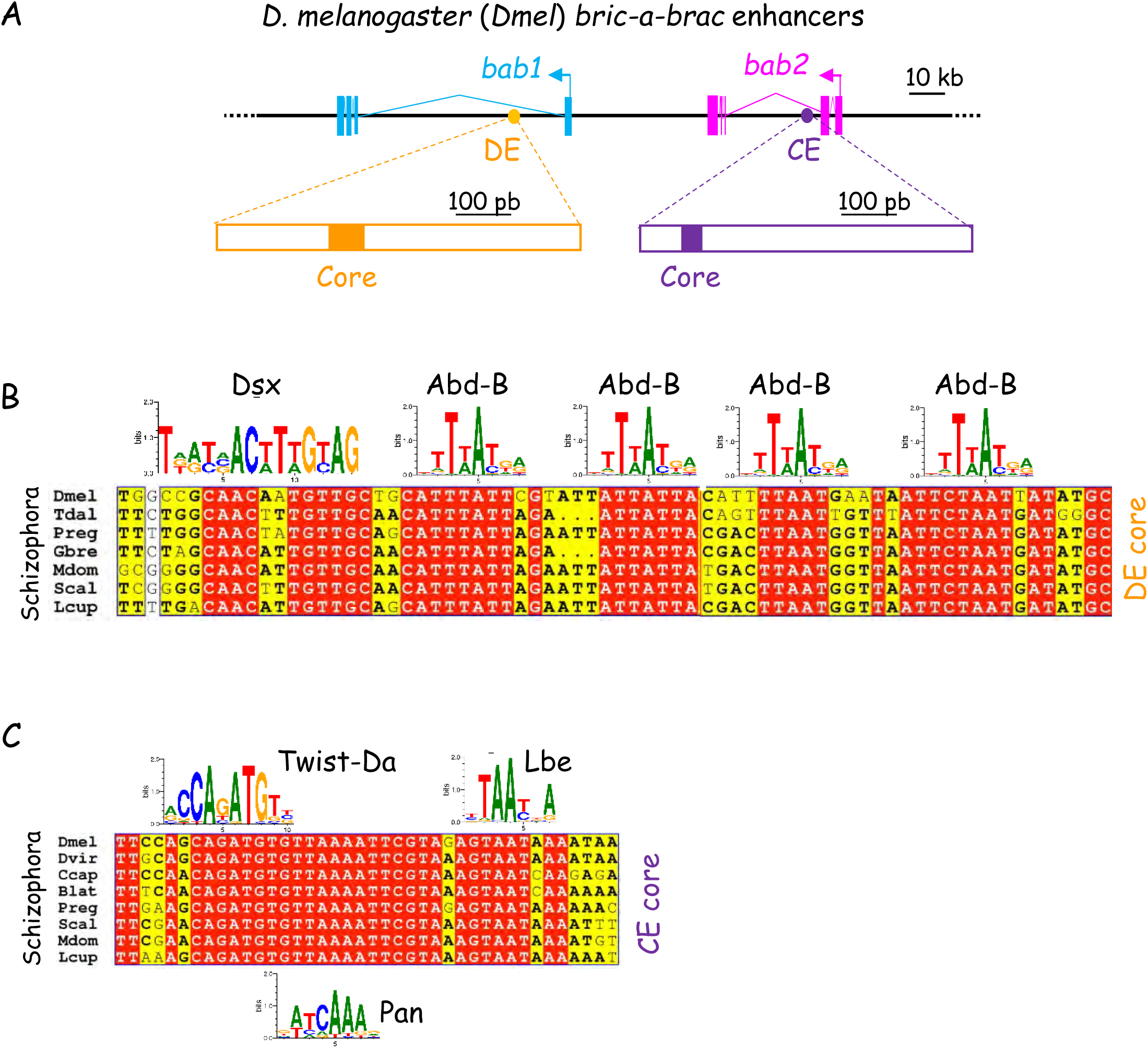

**S10 Figure.**
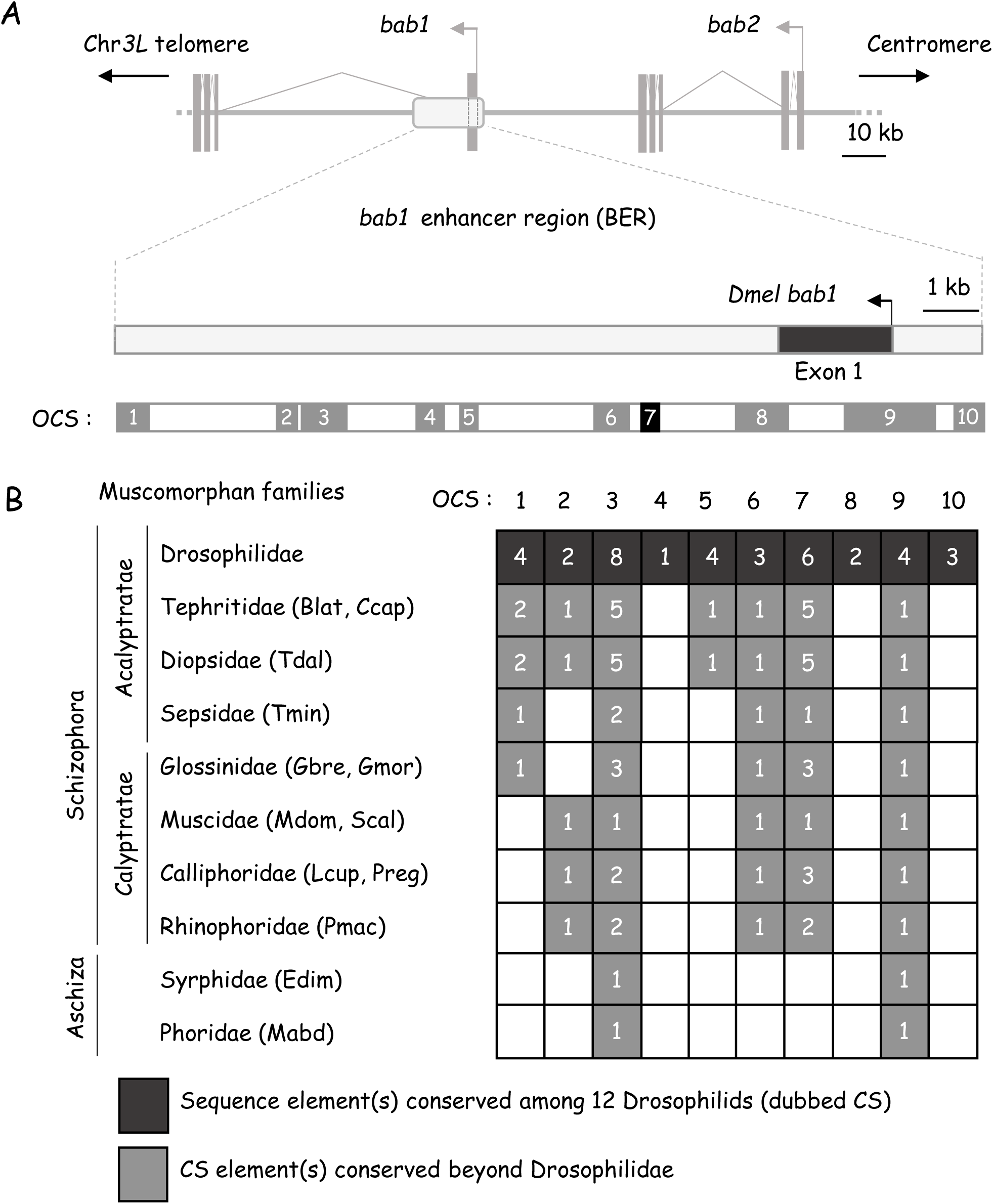

**S11 Figure.**
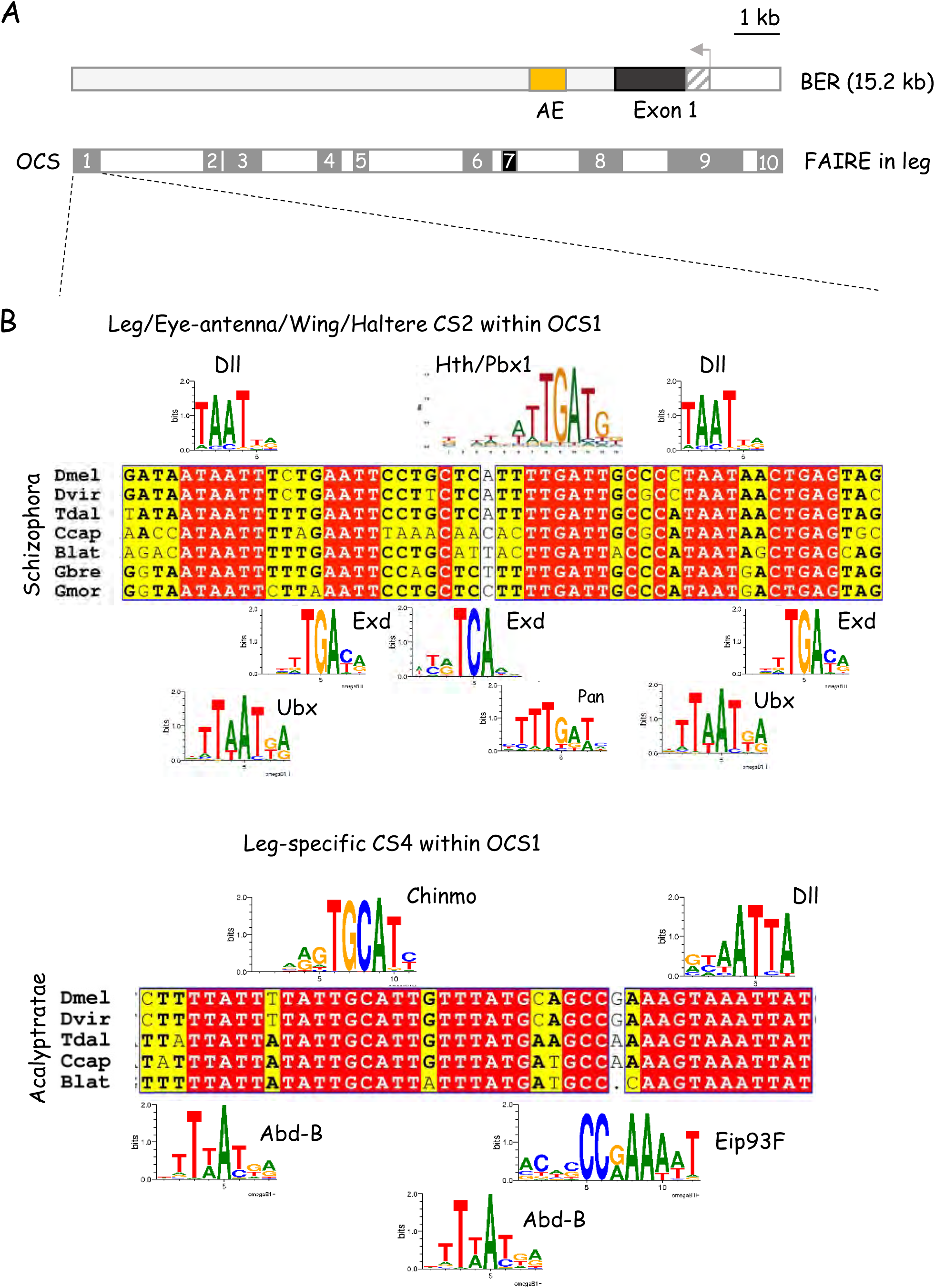

**S12 Figure.**
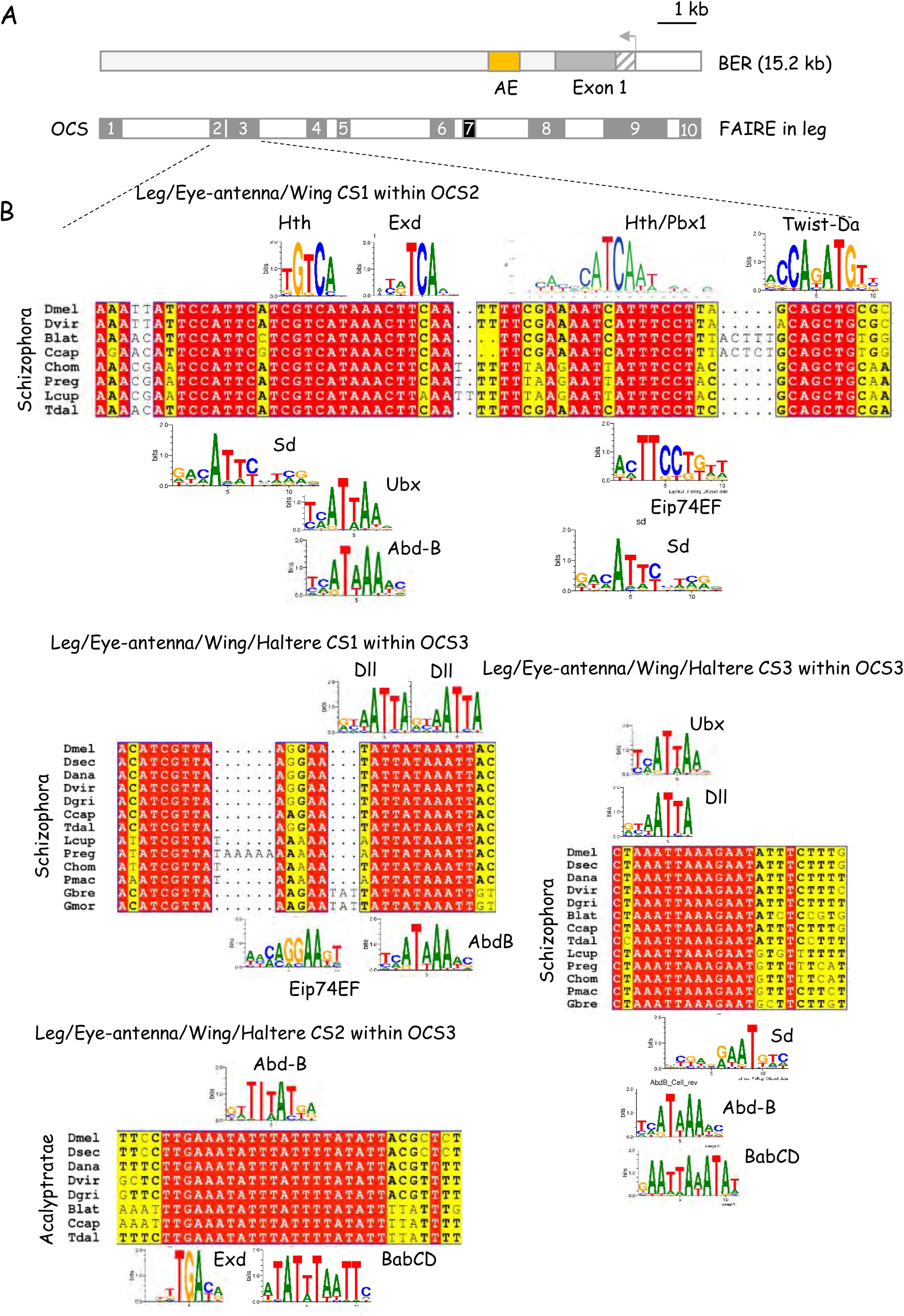

**S13 Figure.**
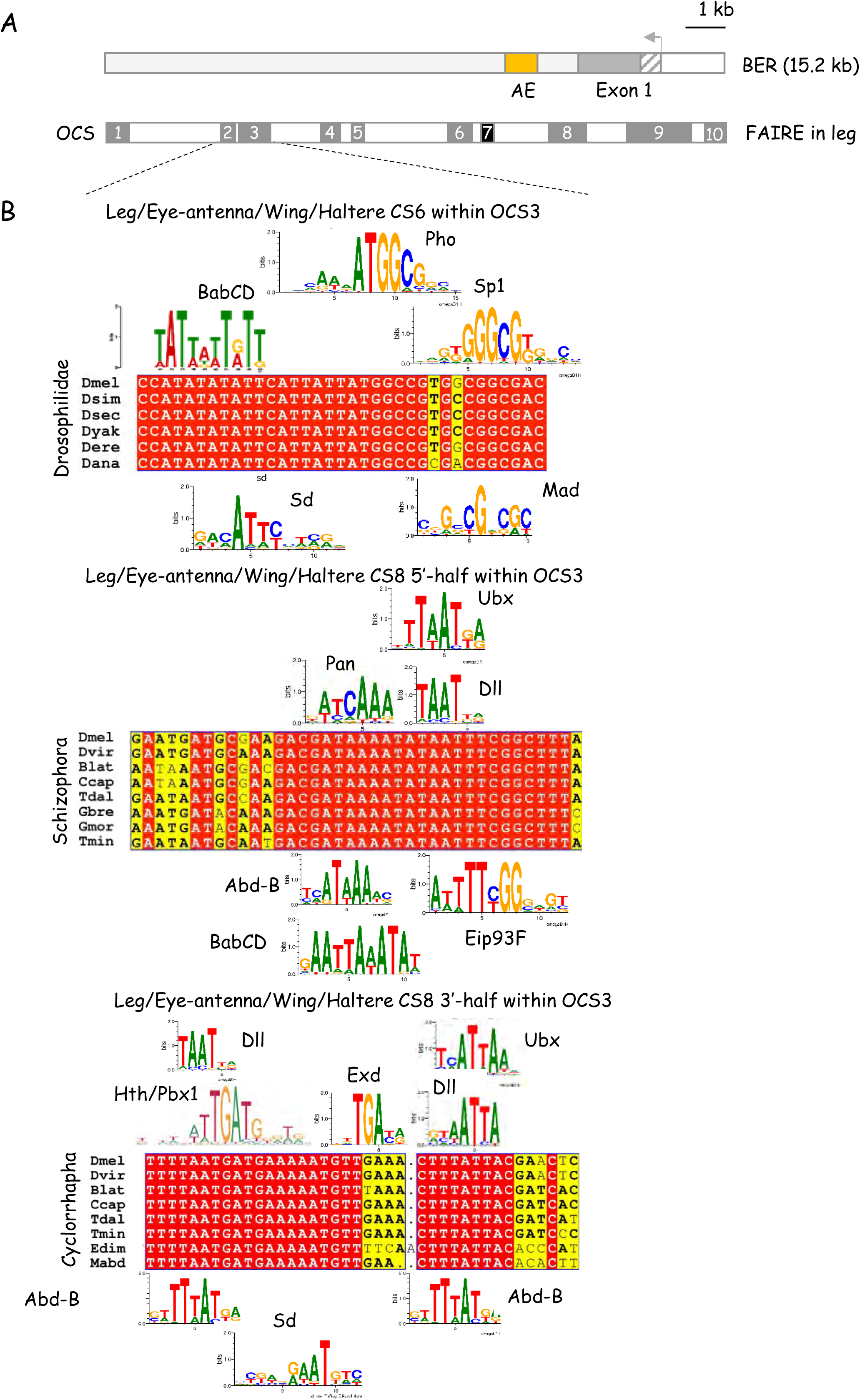

**S14 Figure.**
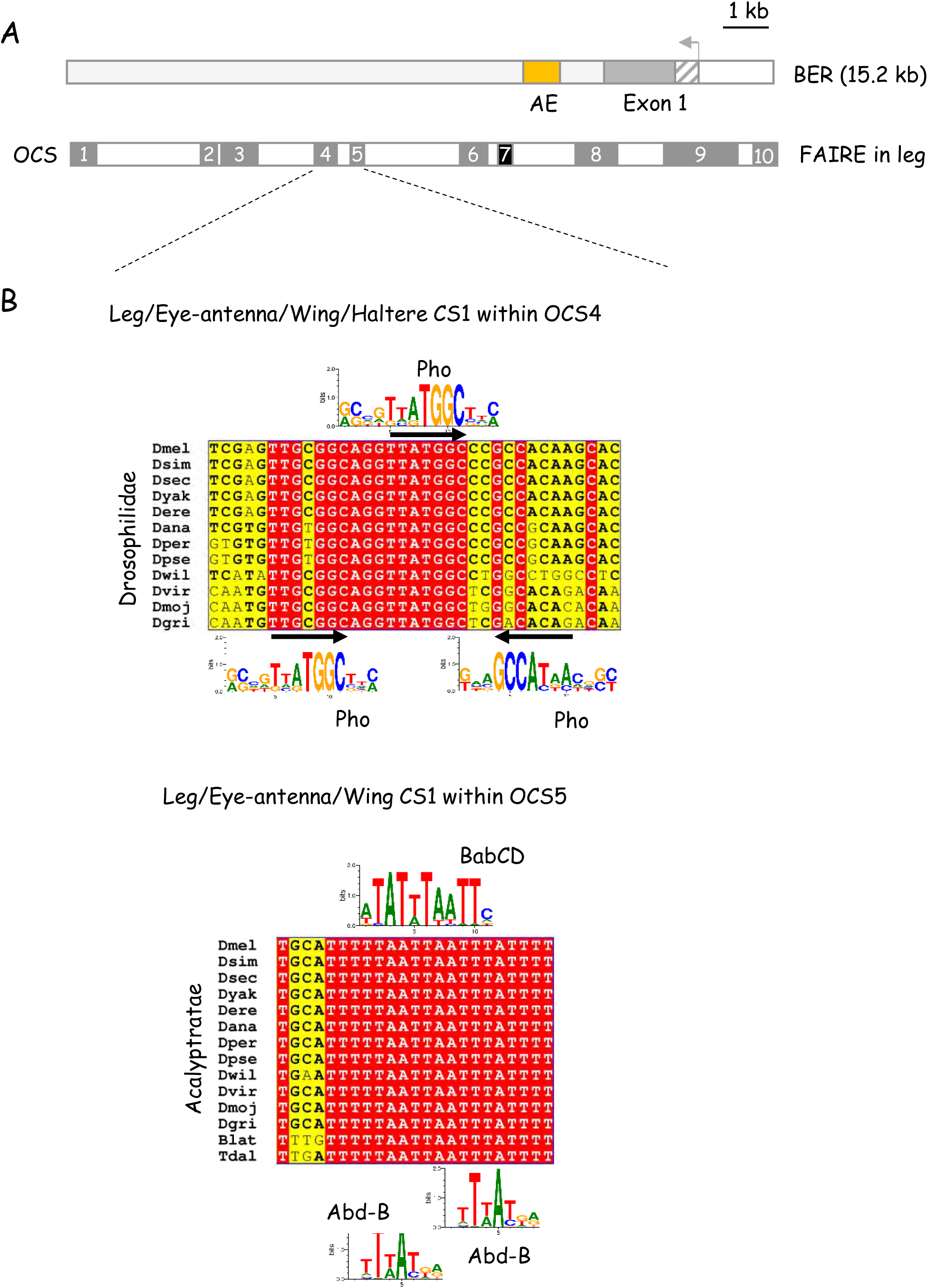

**S15 Figure.**
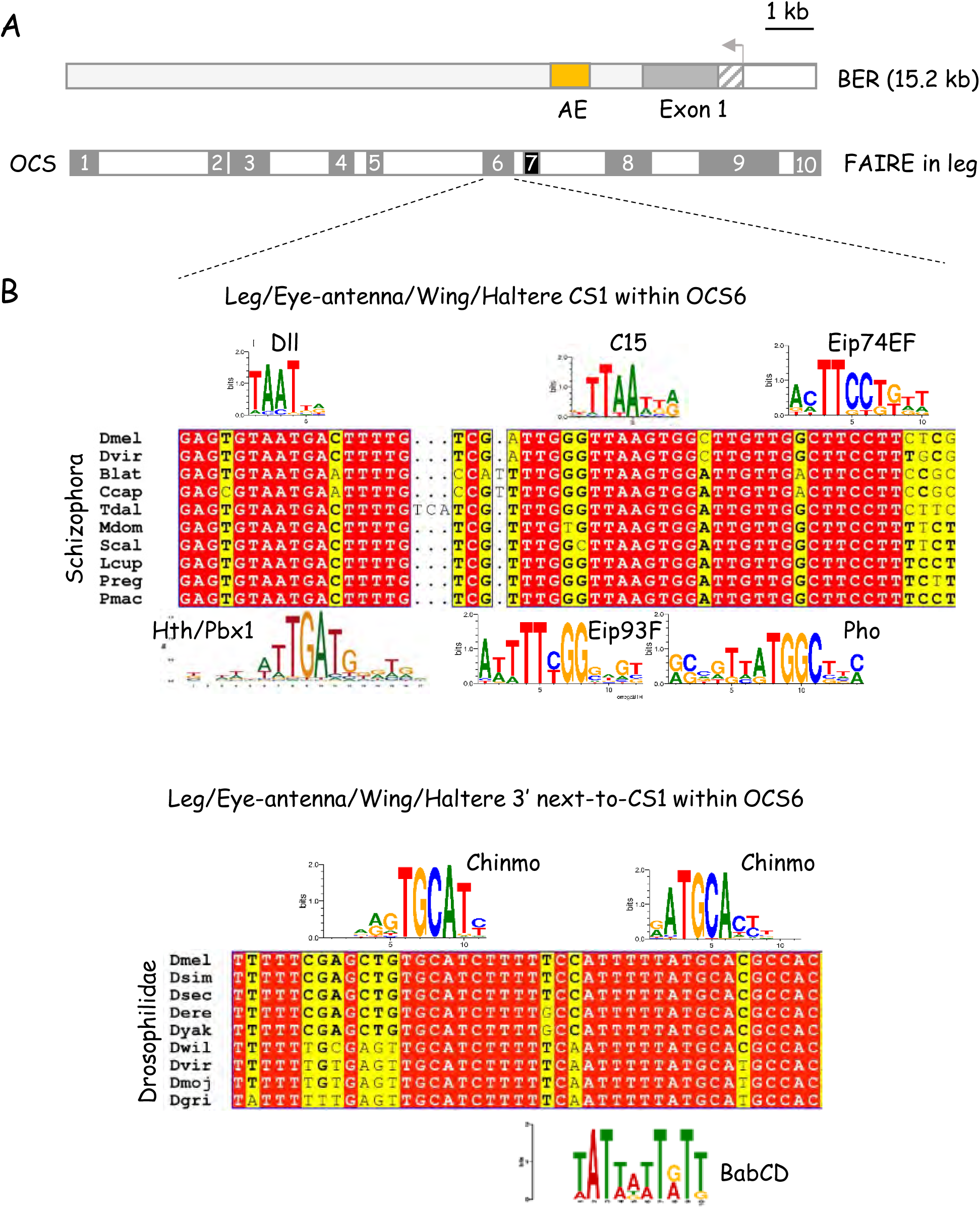

**S16 Figure.**
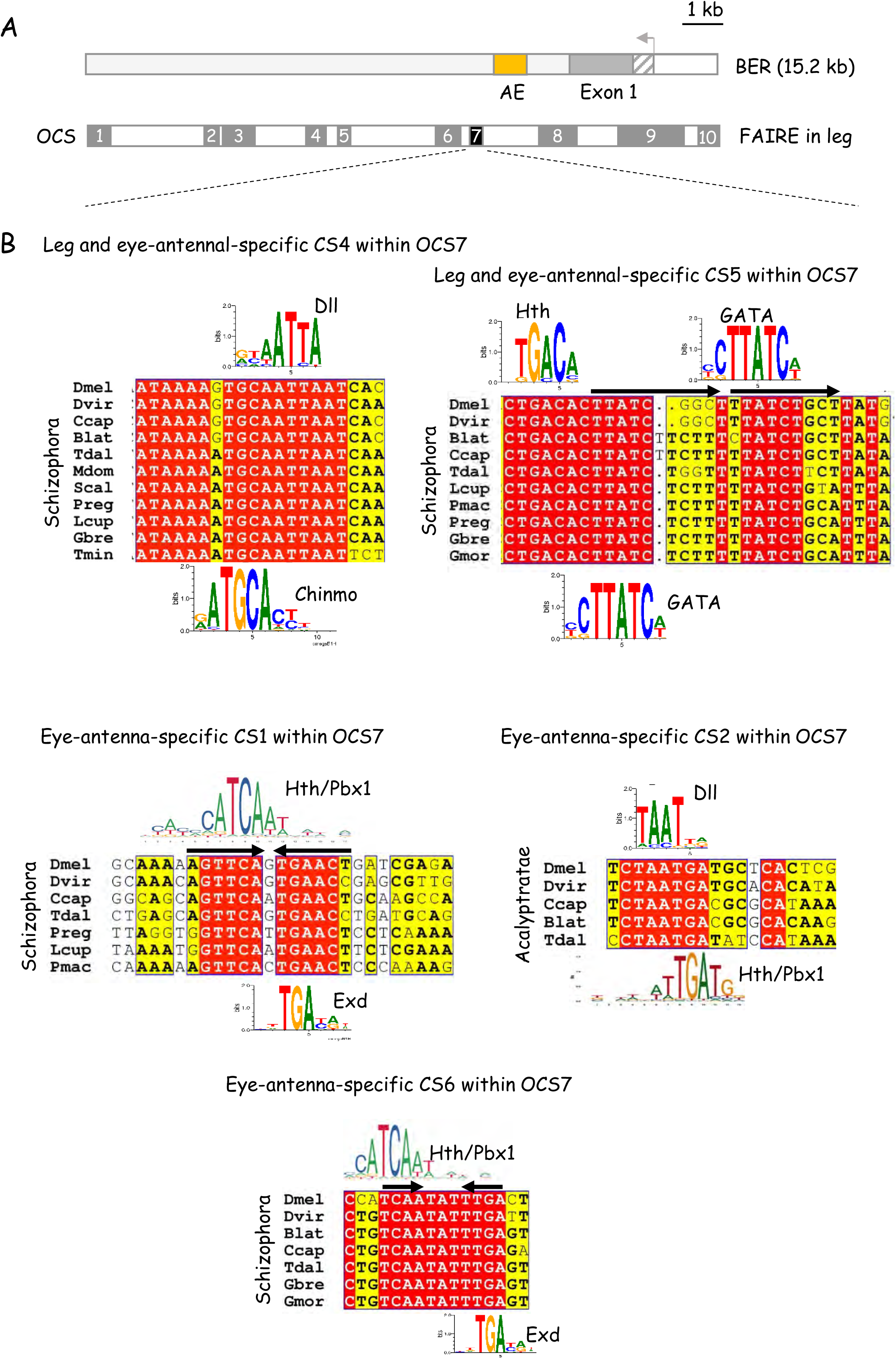

**S17 Figure.**
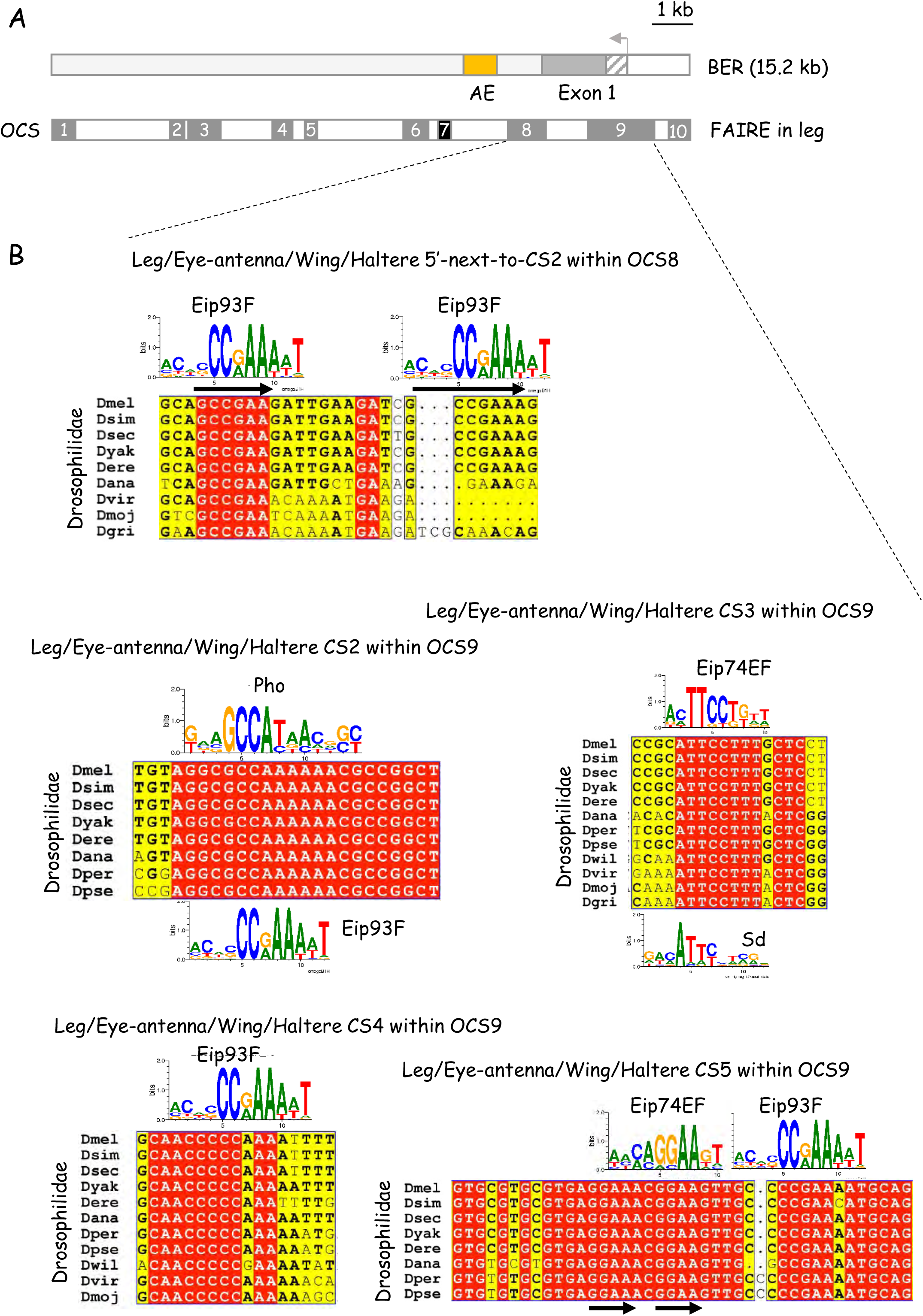

**S18 Figure.**
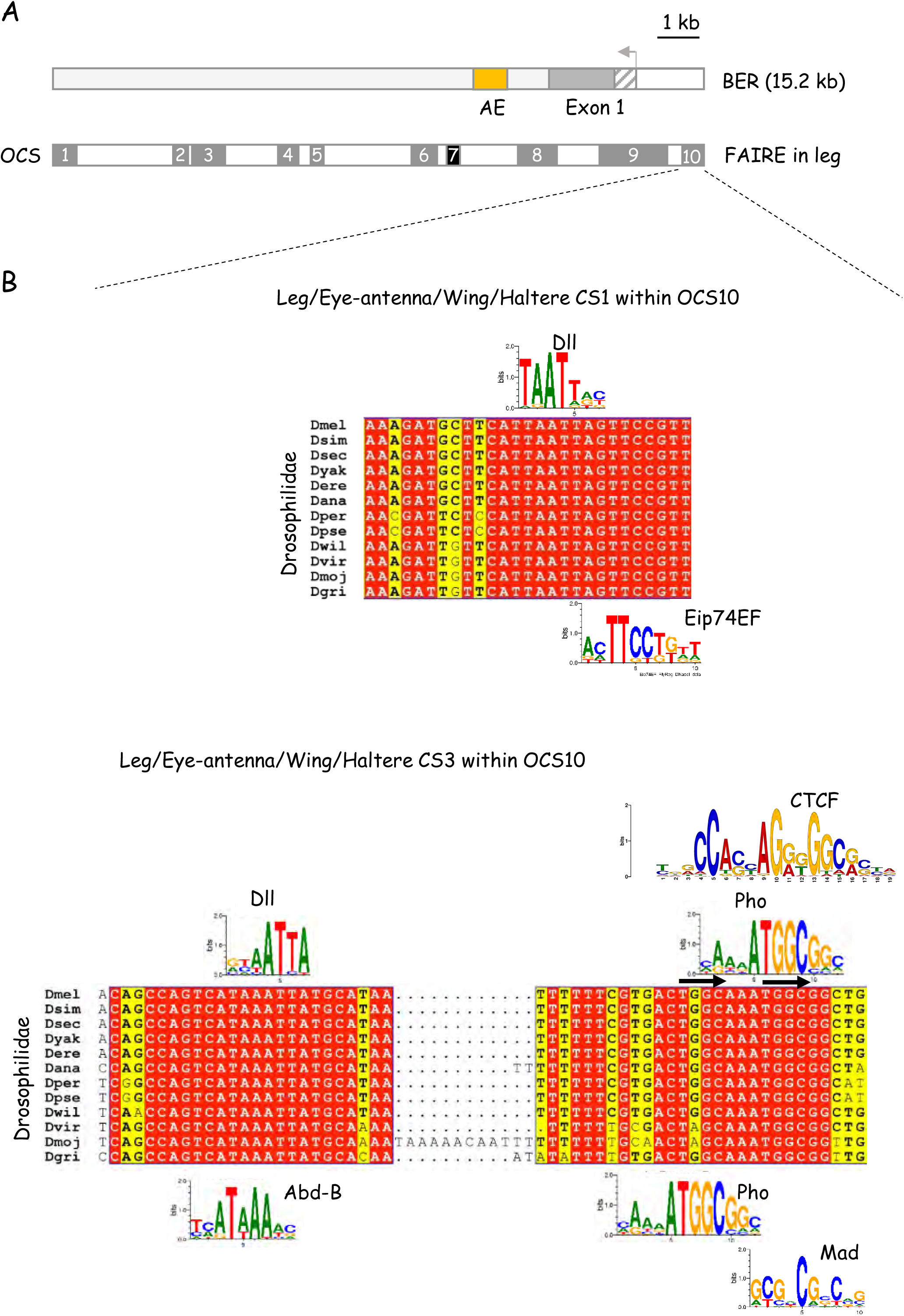

**S19 Figure.**
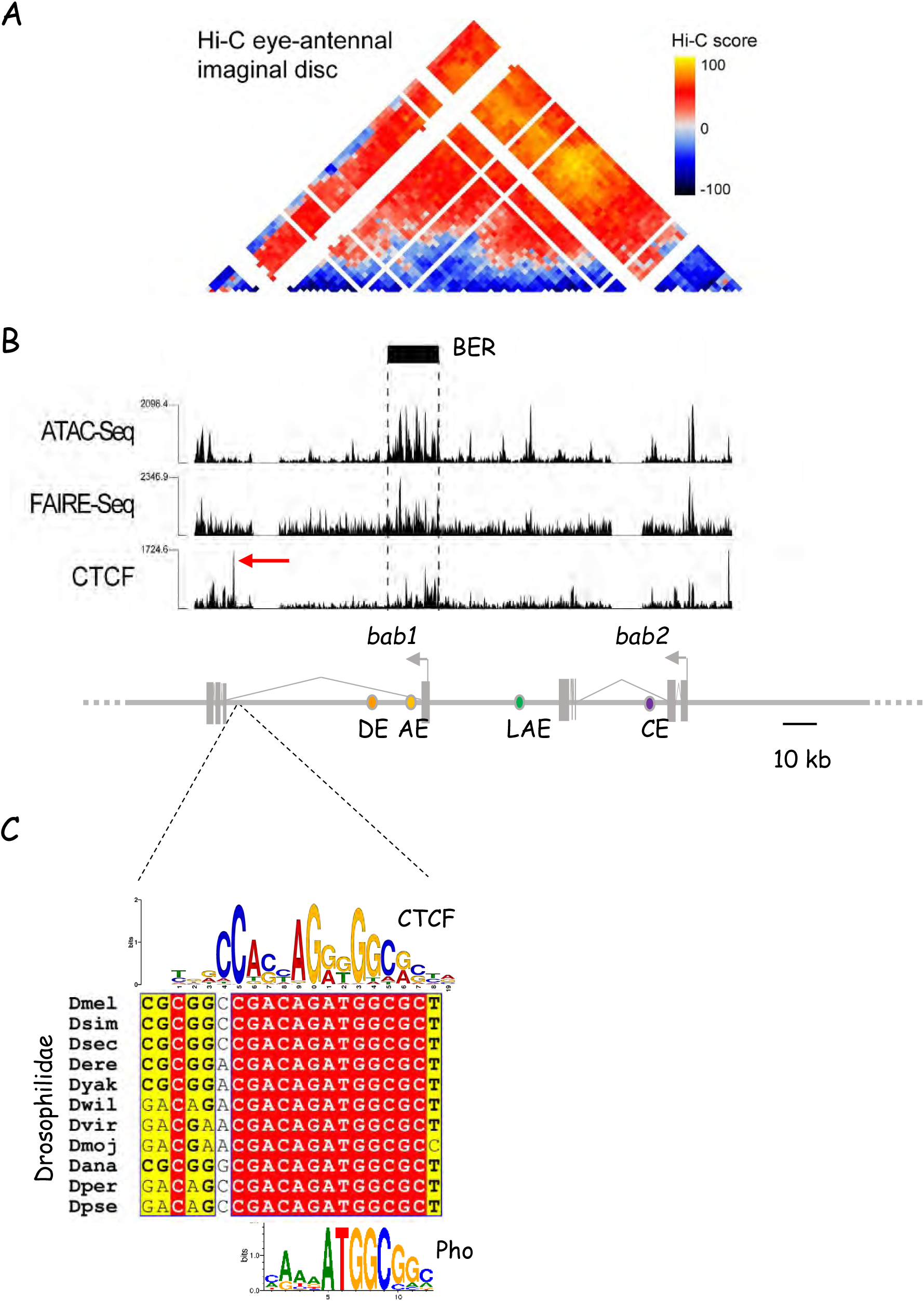

## Supplementary data

Sequence conservation between paralogous Bab1/2 proteins among muscomorphans. The four-letter species abbreviations are as listed below (page 2). Strictly conserved amino-acid residues are indicated by white characters on a red background while partially conserved ones are in black characters on a yellow background. Locations of the strongly-conserved BTB and BabCD domains are indicated along the right side (see black lines).

Conservation among twelve reference drosophilids of *D. melanogaster* BER OCS1-10 as well as LAE, CE, AE and DE sequences. The four-letter Drosophilidae species abbreviations are as listed below (page 2). Locations of conserved sequence elements (CS) are indicated by underneath black bars. Sequence LOGOs of predicted binding sites for the Dll, Bowl, C15, Sp1, Rn, Sal/Salr, BabCD, GAF, Pho, Eip74EF, Dsx, Abd-B, Sd, Chinmo, Pan, Mad, GATA factors, Twist-Da and Lbe transcription factors are depicted above or below the alignments.

Four letter abbreviations for investigated species

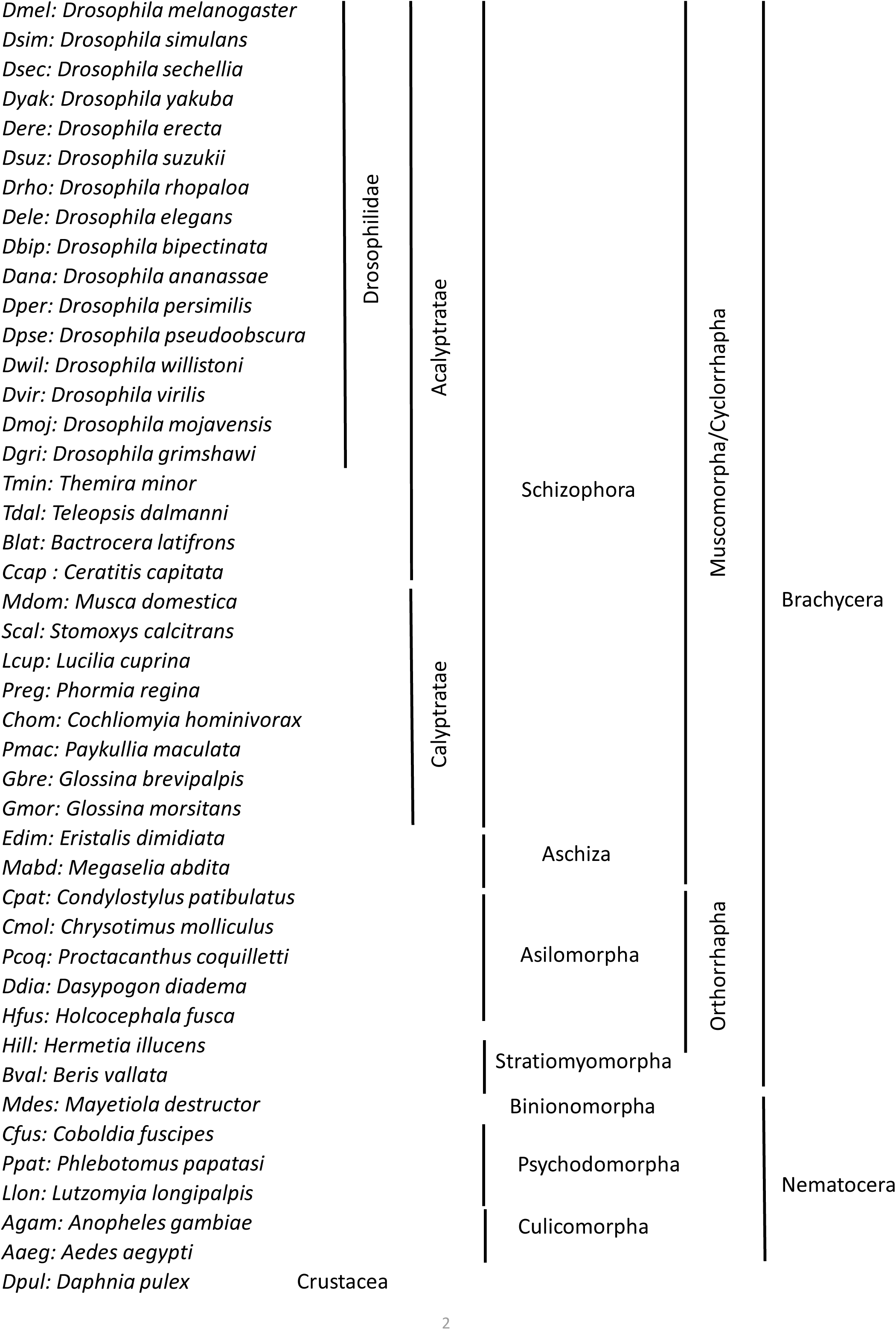

Bab1 sequence conservation among muscomorphans (Part1)

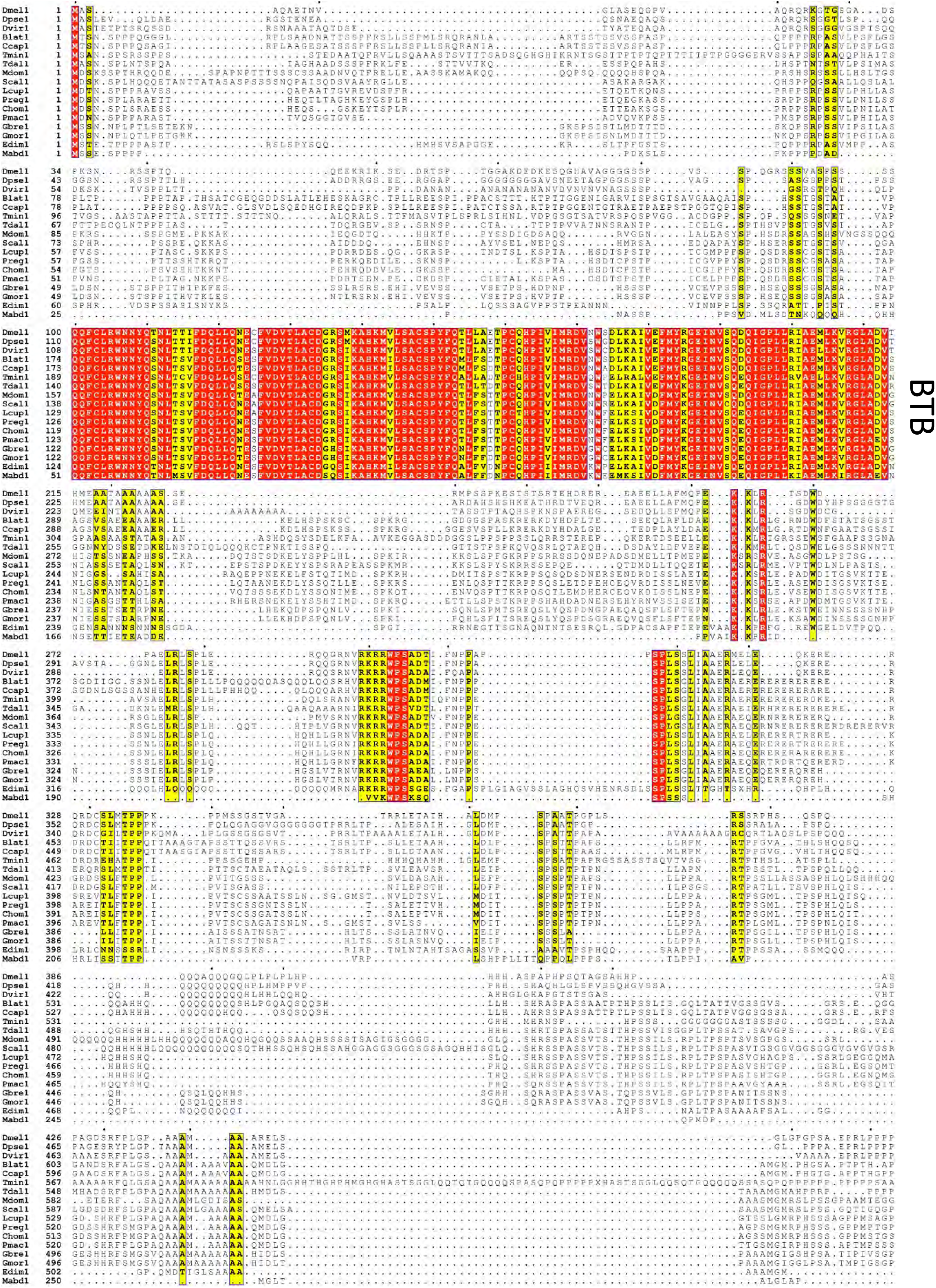

Bab1 sequence conservation among muscomorphans (Part2)

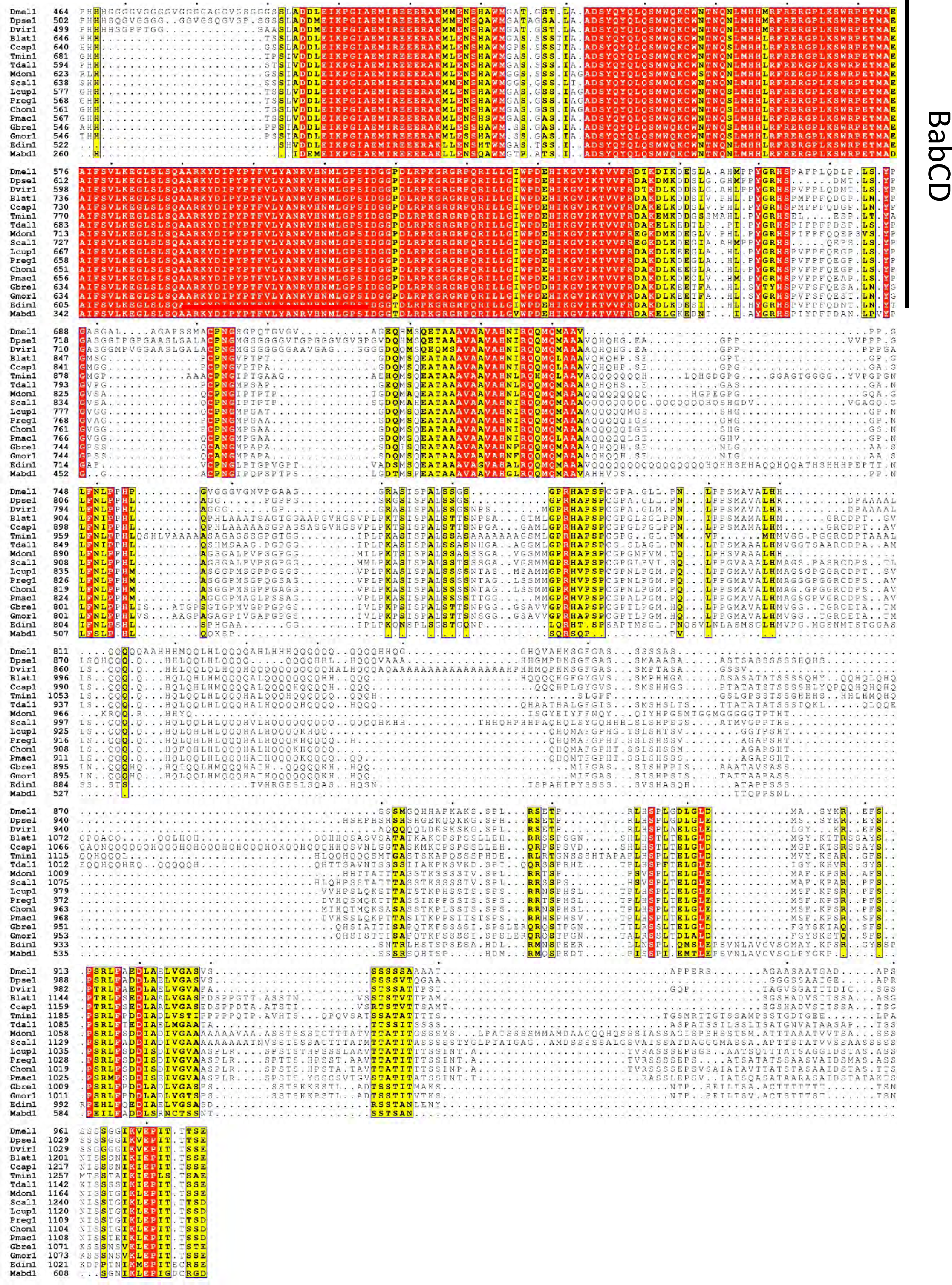

Bab2 sequence conservation among muscomorphans (Part1)

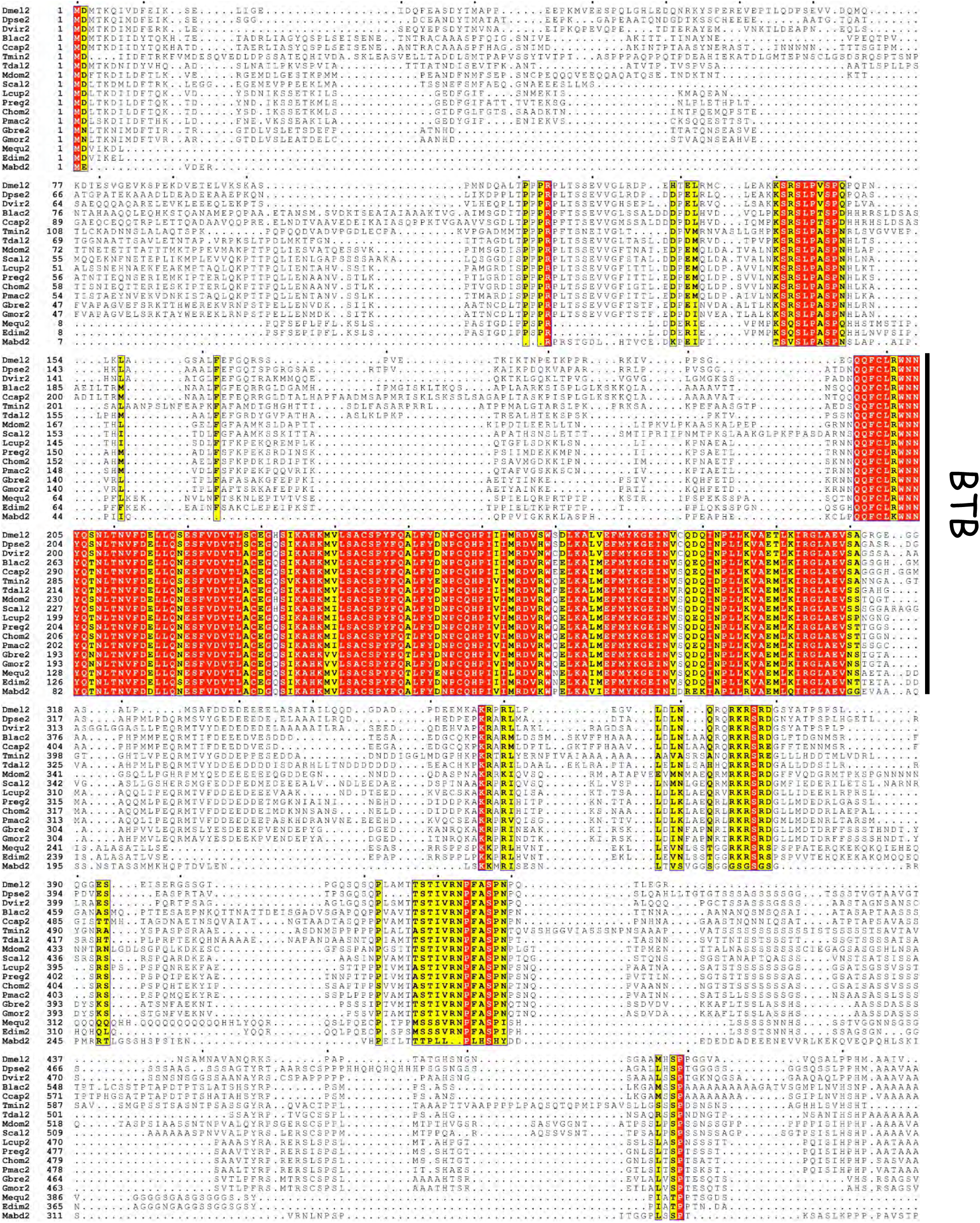

Bab2 sequence conservation among muscomorphans (Part2)

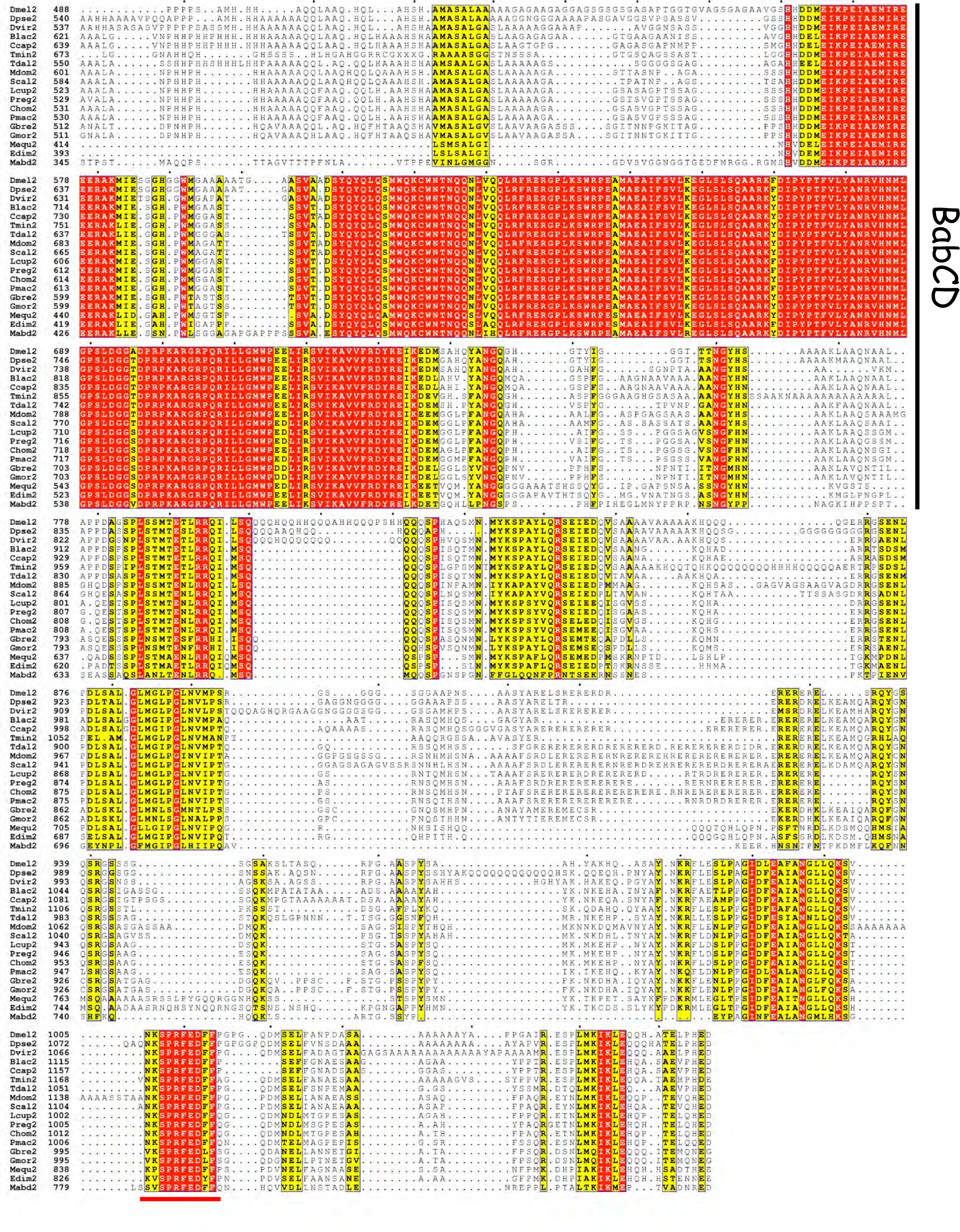

Sequence conservation between Bab1/2 paralogs (Part1)

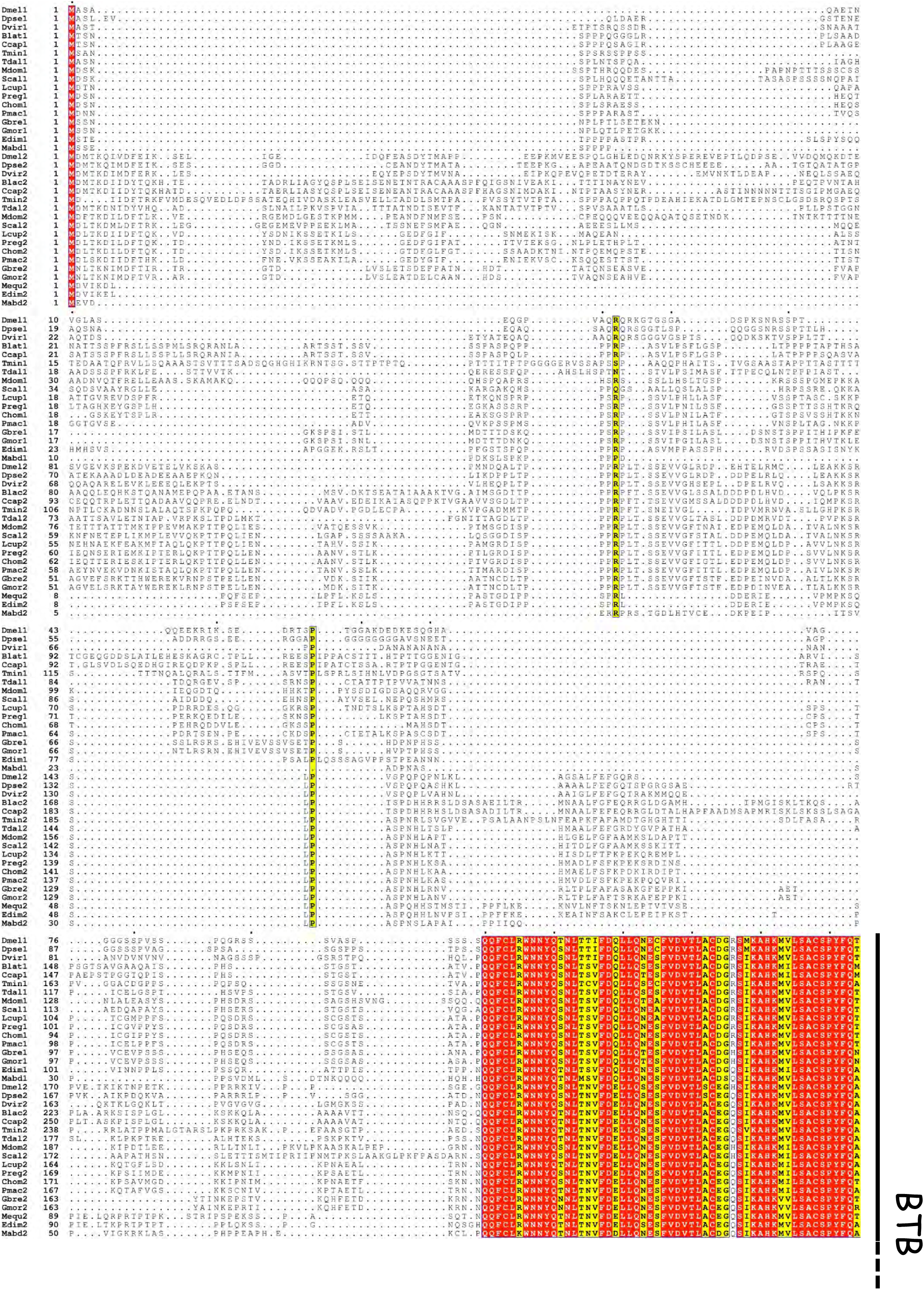

Sequence conservation between Bab1/2 paralogs (Part2)

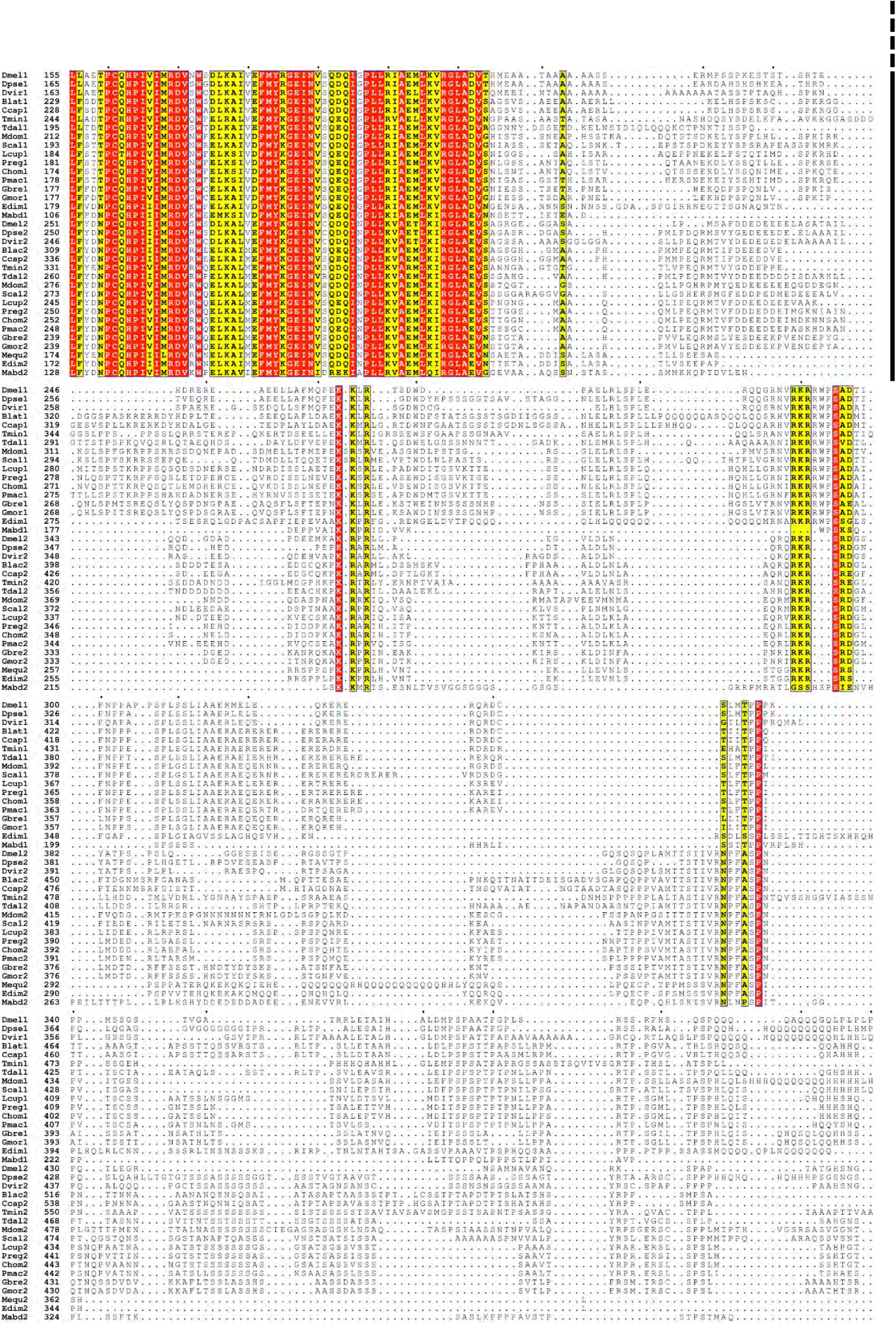

Sequence conservation between Bab1/2 paralogs (Part3)

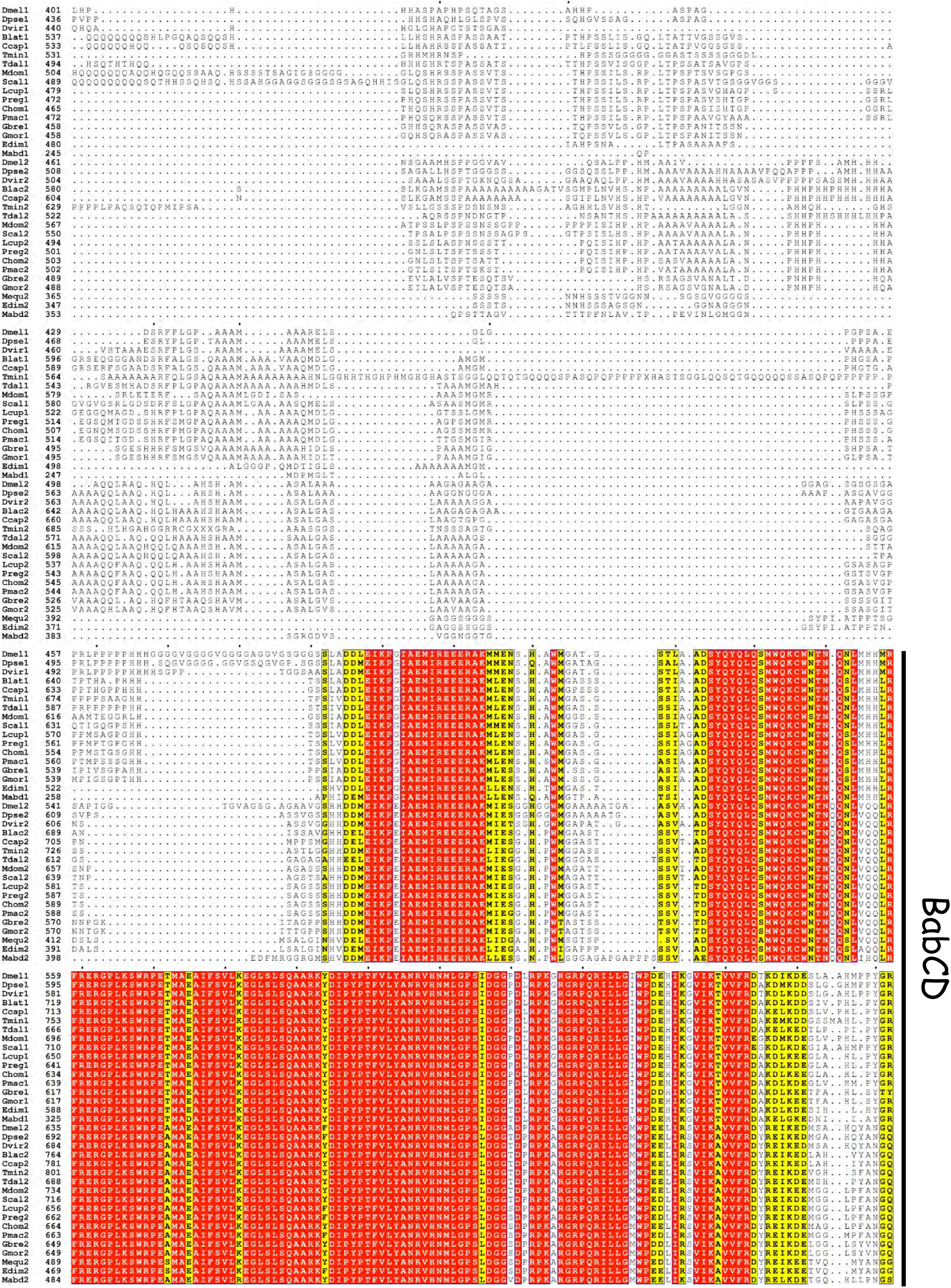

Sequence conservation between Bab1/2 paralogs (Part4)

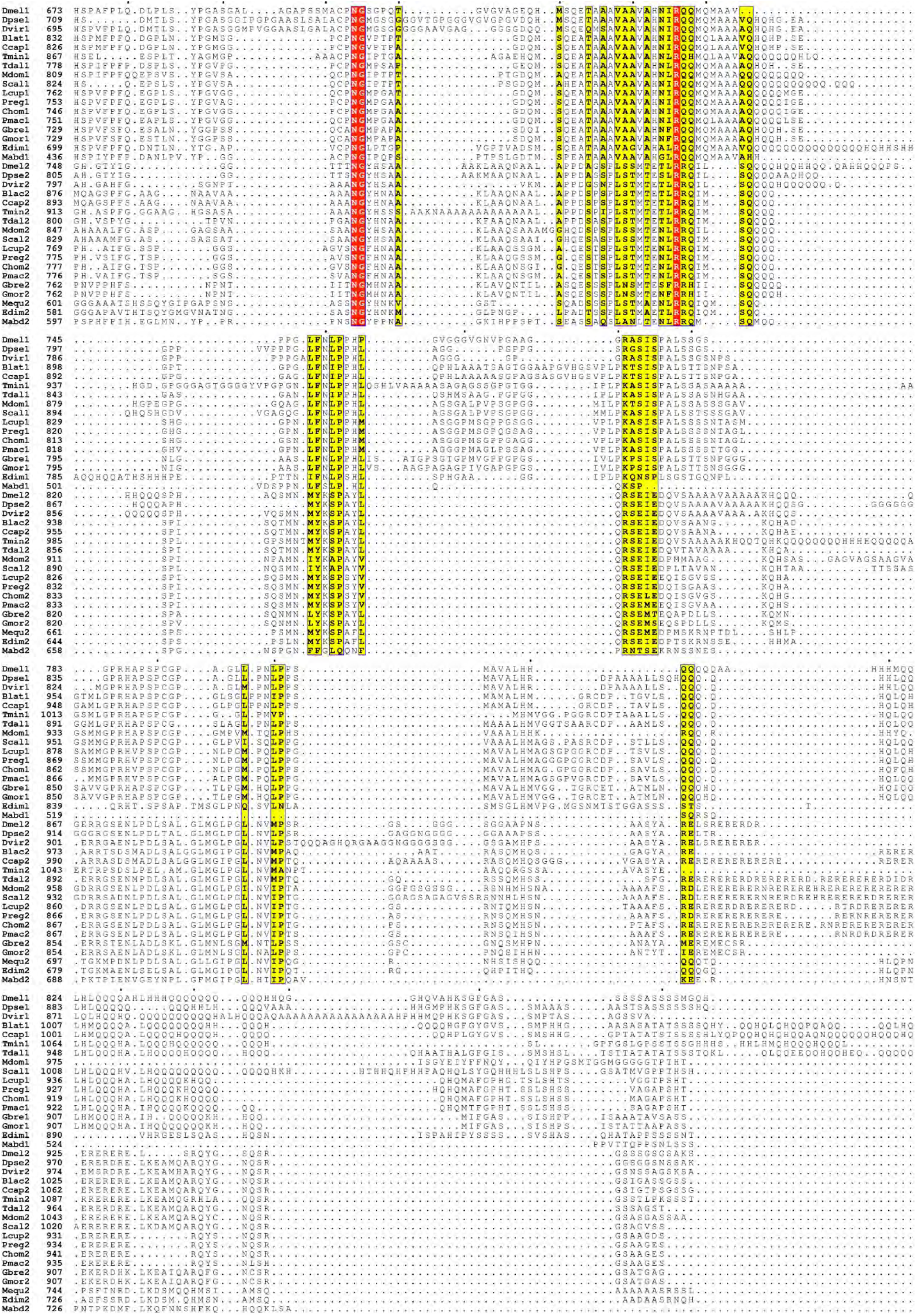

Sequence conservation between Bab1/2 paralogs (Part5)

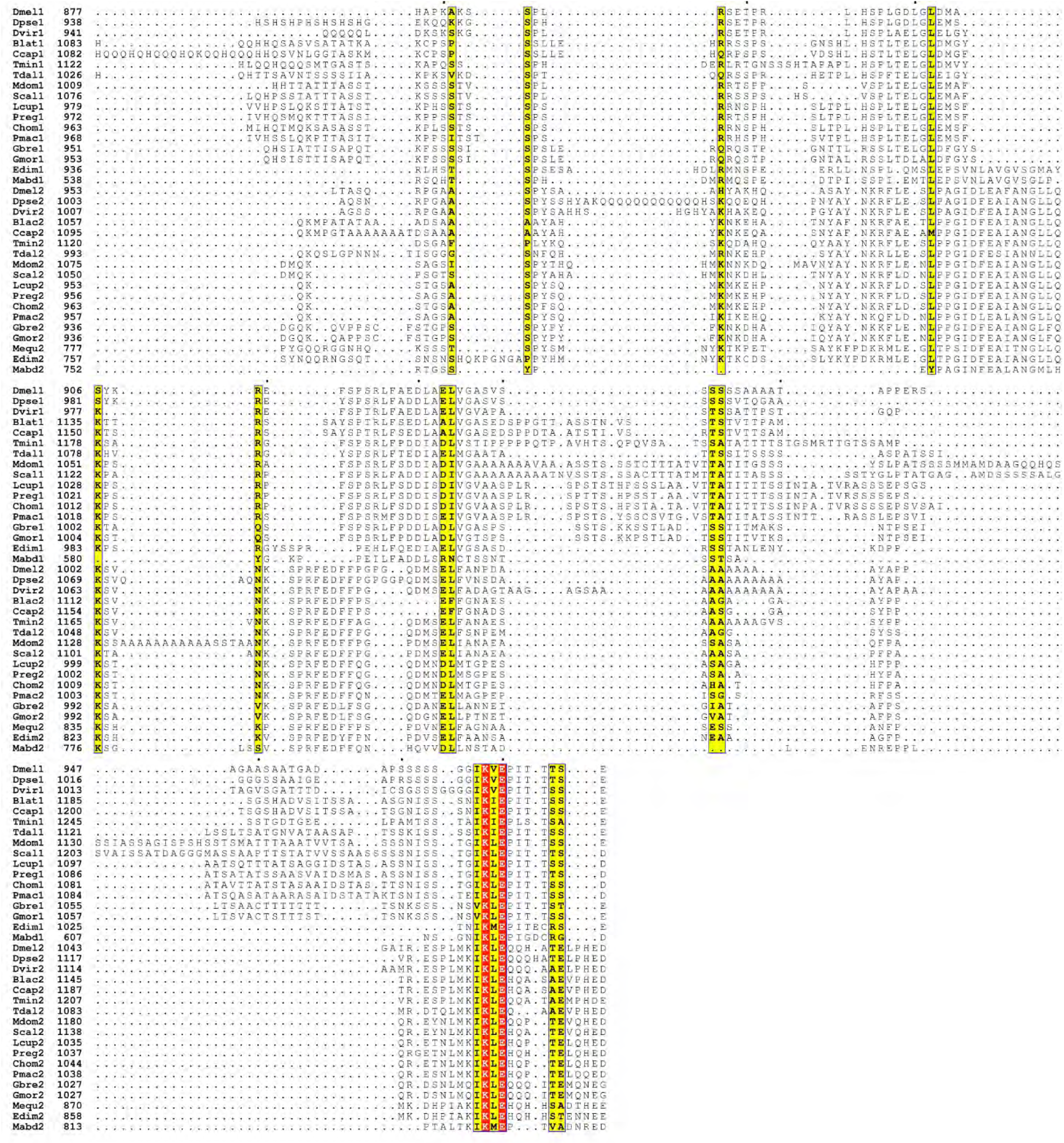

BER^OCS1^ sequence conservation among Drosophilidae (Part1)

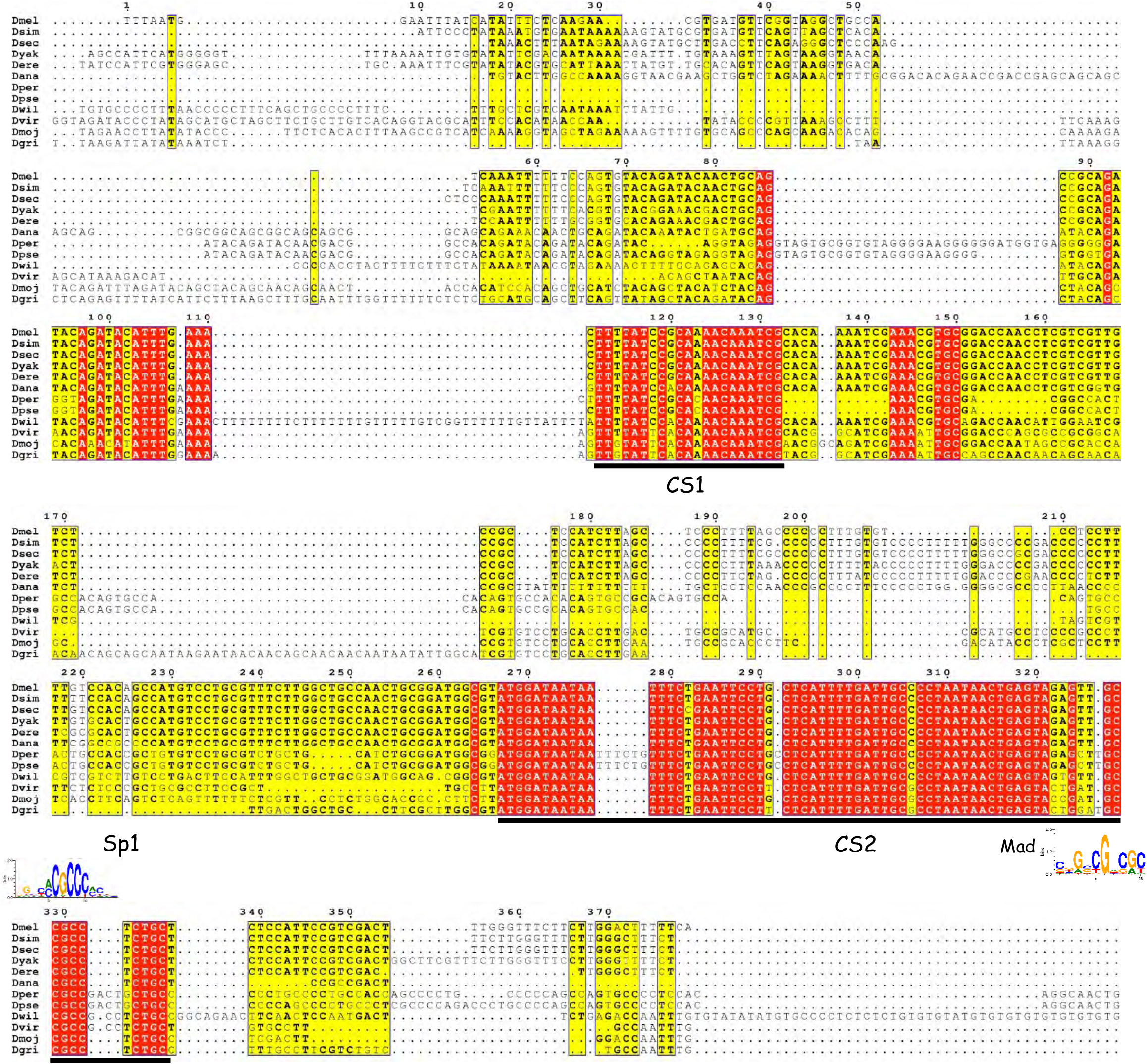

BER^OCS1^ sequence conservation among Drosophilidae (Part2)

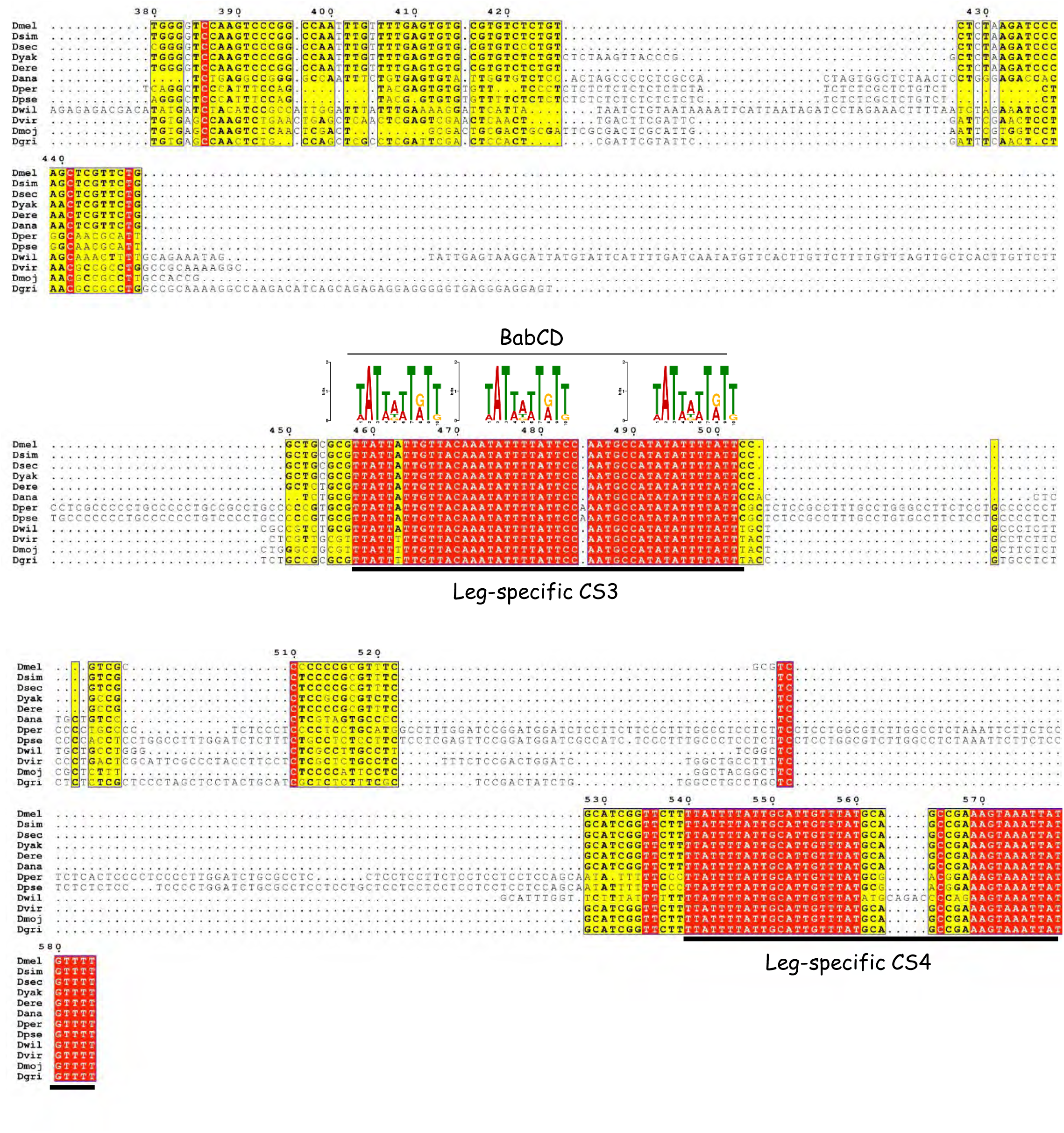

BER^OCS2^ sequence conservation among Drosophilidae

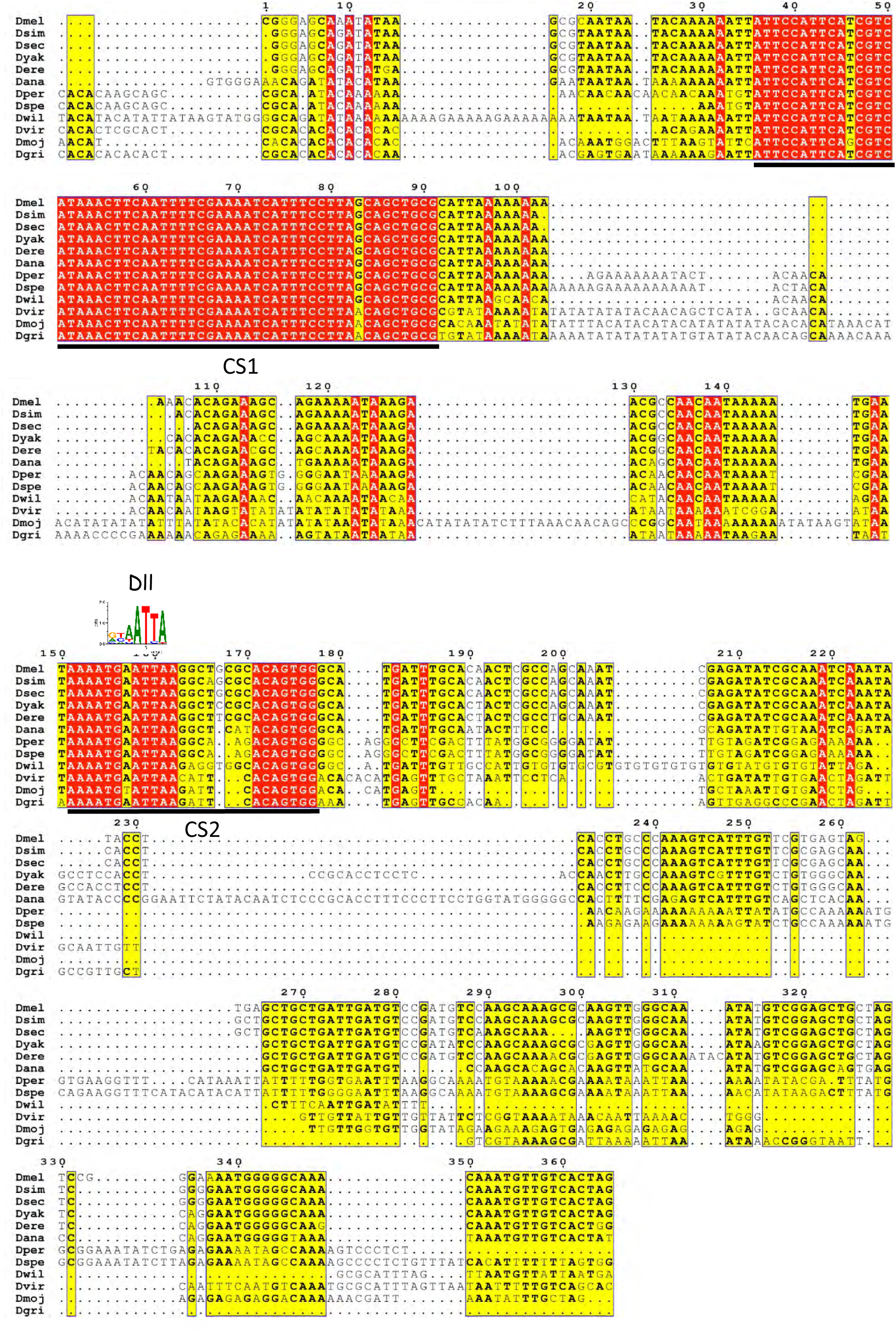

BER^OCS3^ sequence conservation among Drosophilidae (Part1)

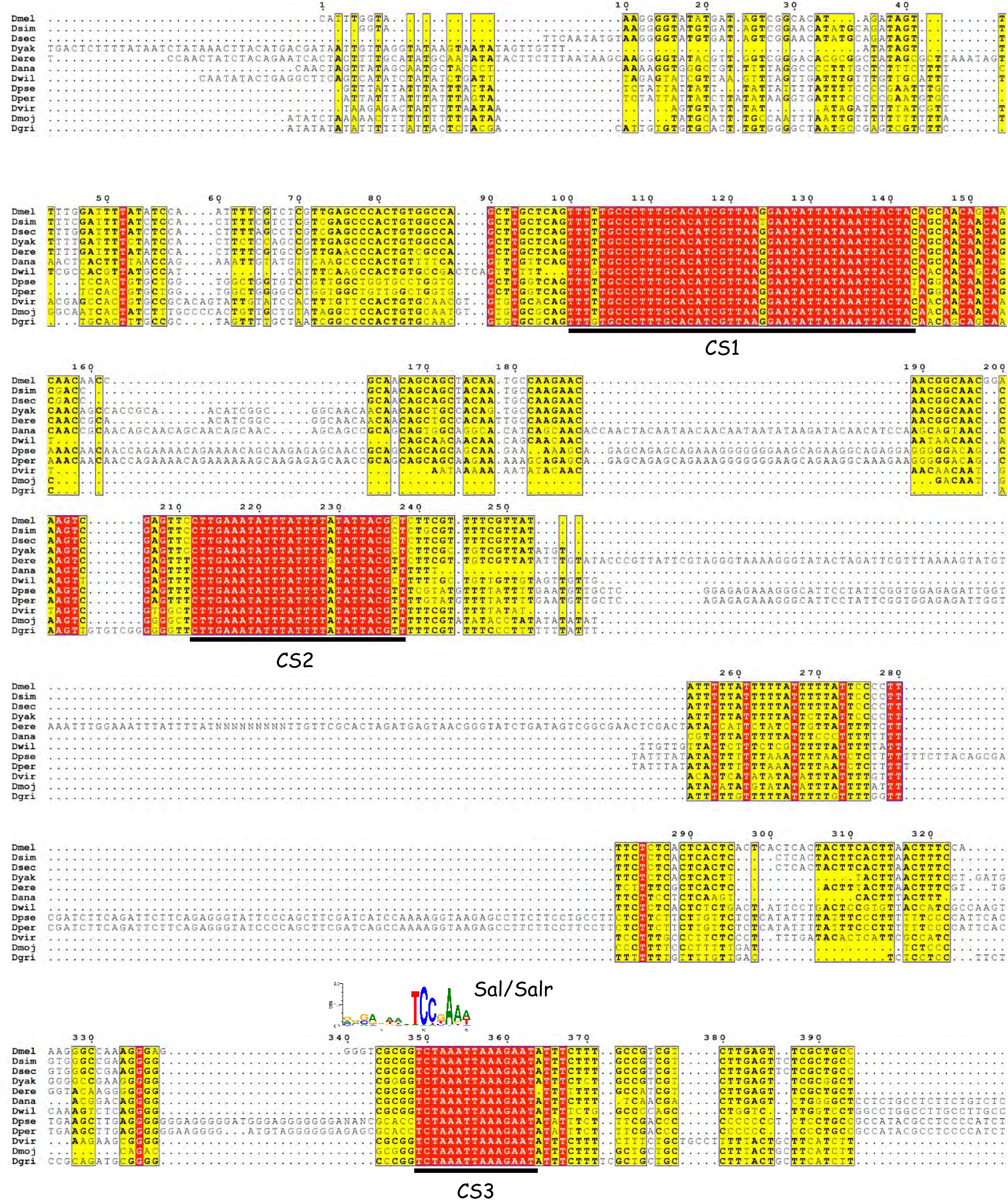

BER^OCS3^ sequence conservation among Drosophilidae (Part2)

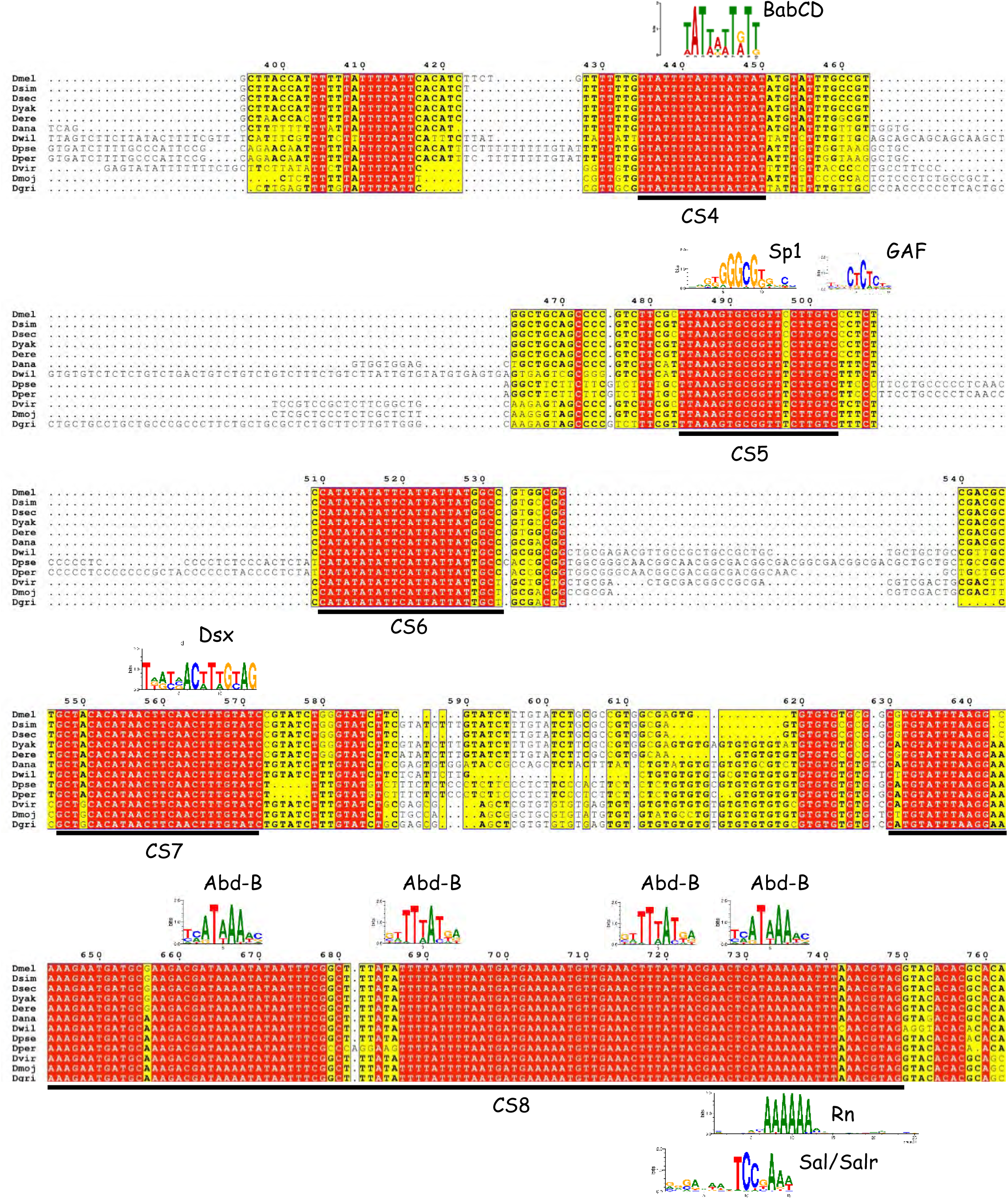

BER^OCS4^ sequence conservation among Drosophilidae

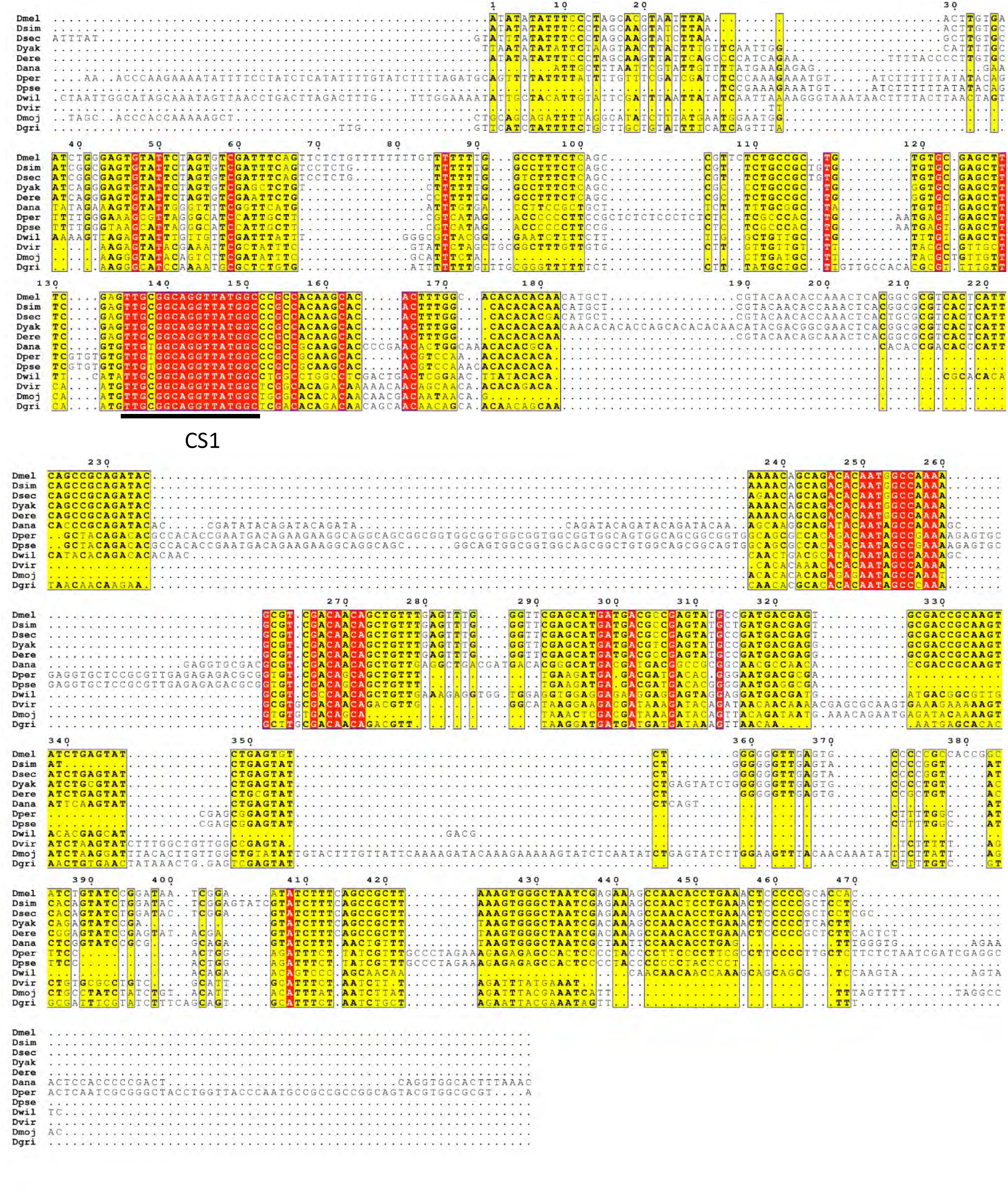

BER^OCS5^ sequence conservation among Drosophilidae

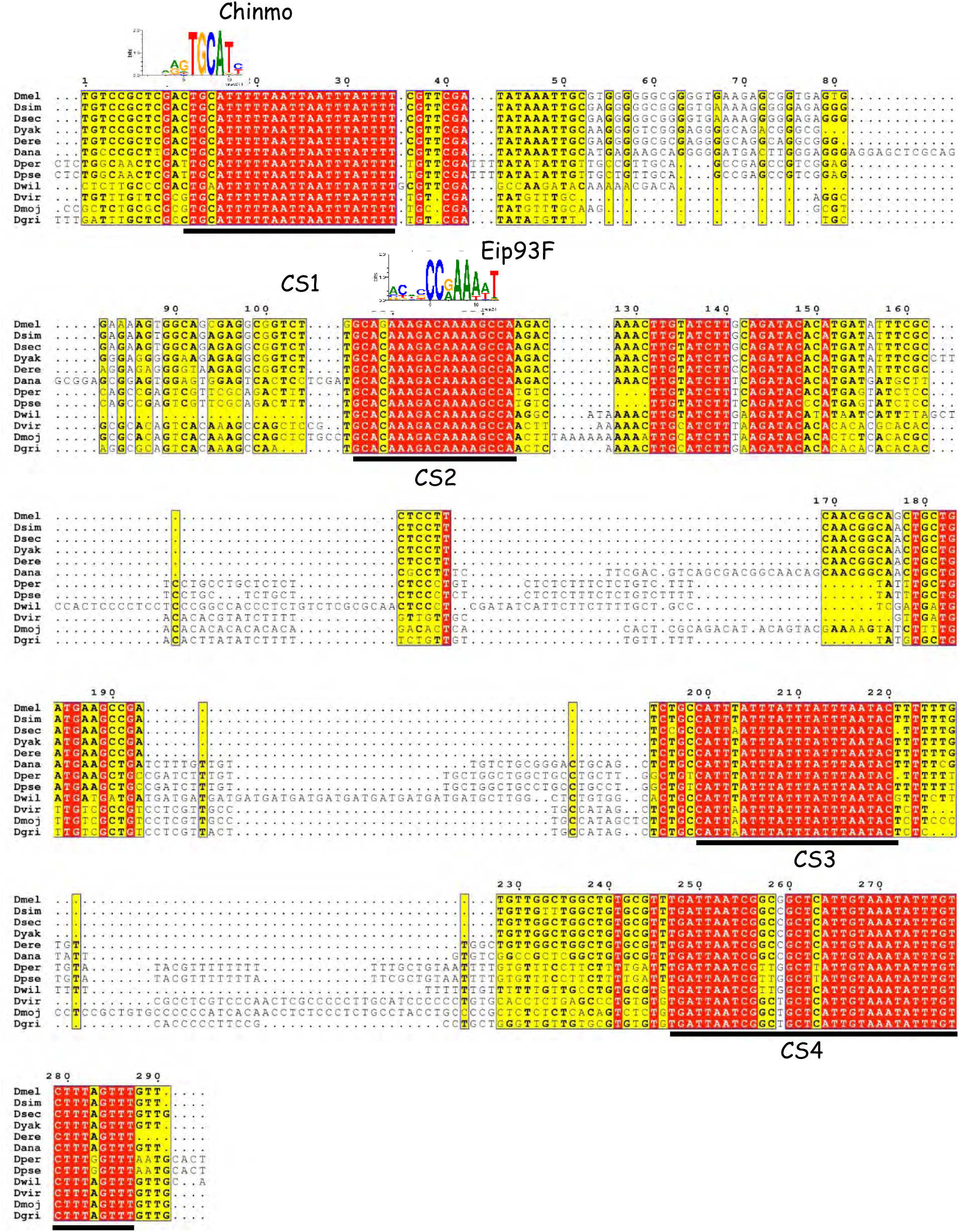

BER^OCS6^ sequence conservation among Drosophilidae (Part1)

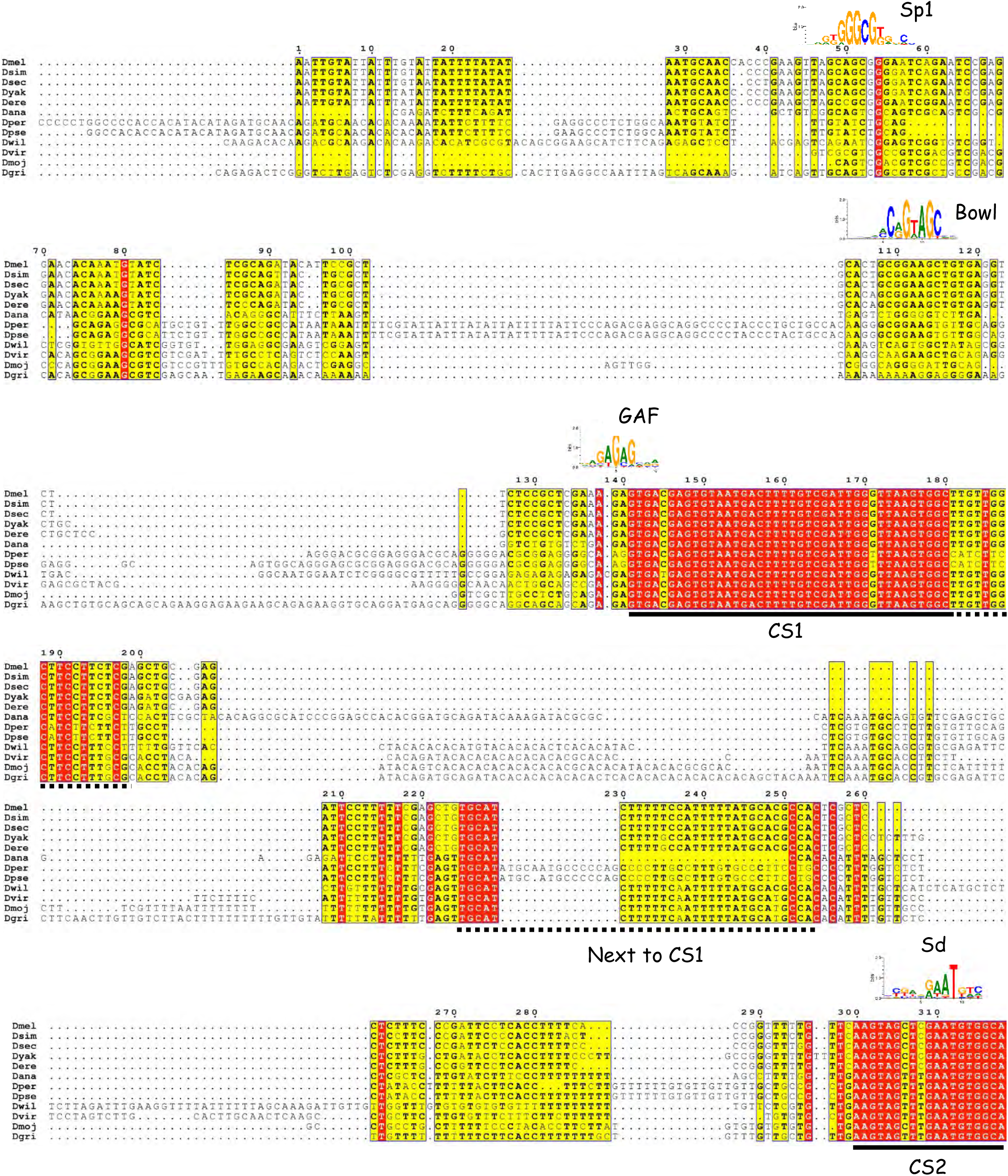

BER^OCS6^ sequence conservation among Drosophilidae (Part2)

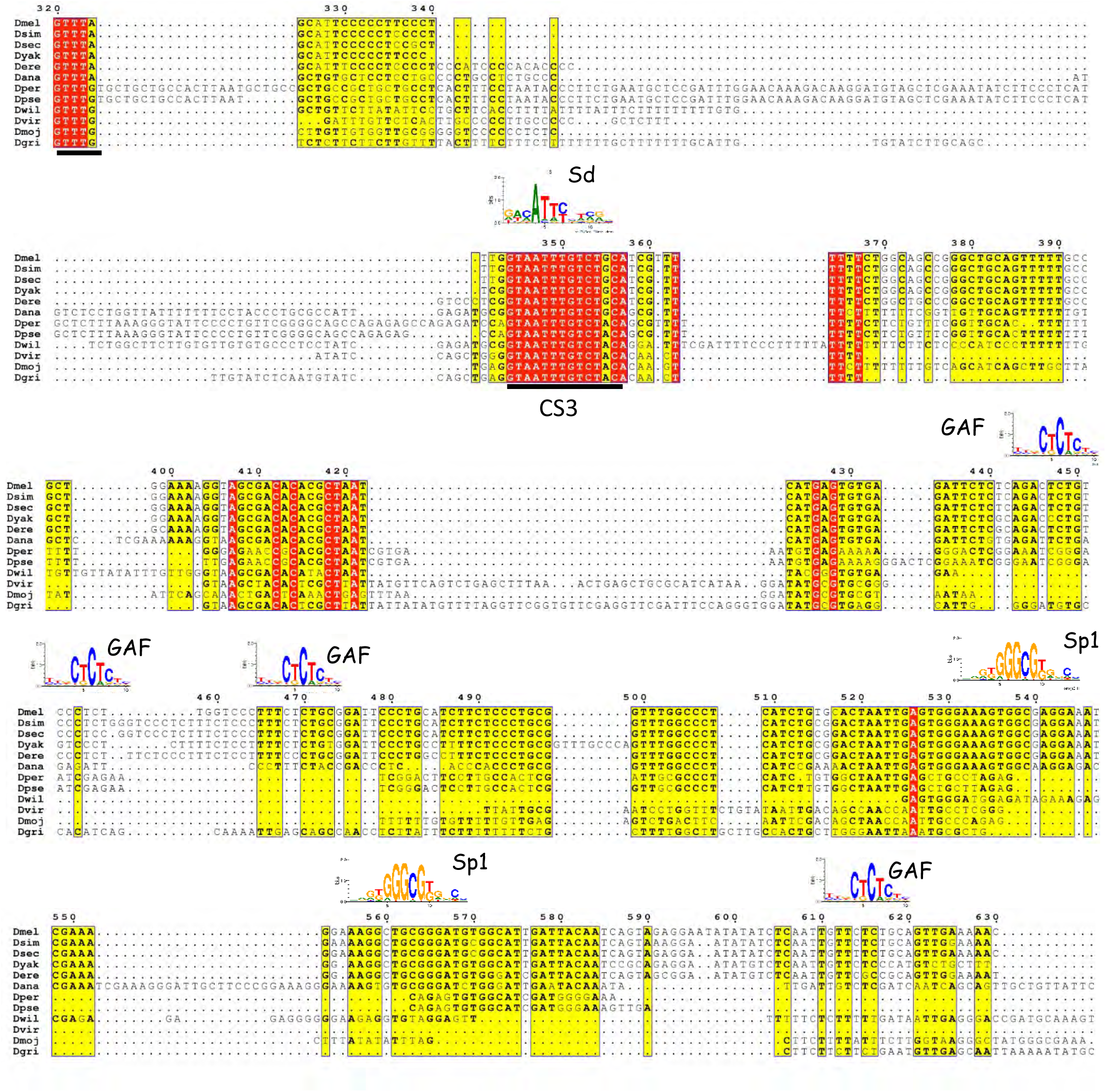

BER^OCS7^ sequence conservation among Drosophilidae (Part1)

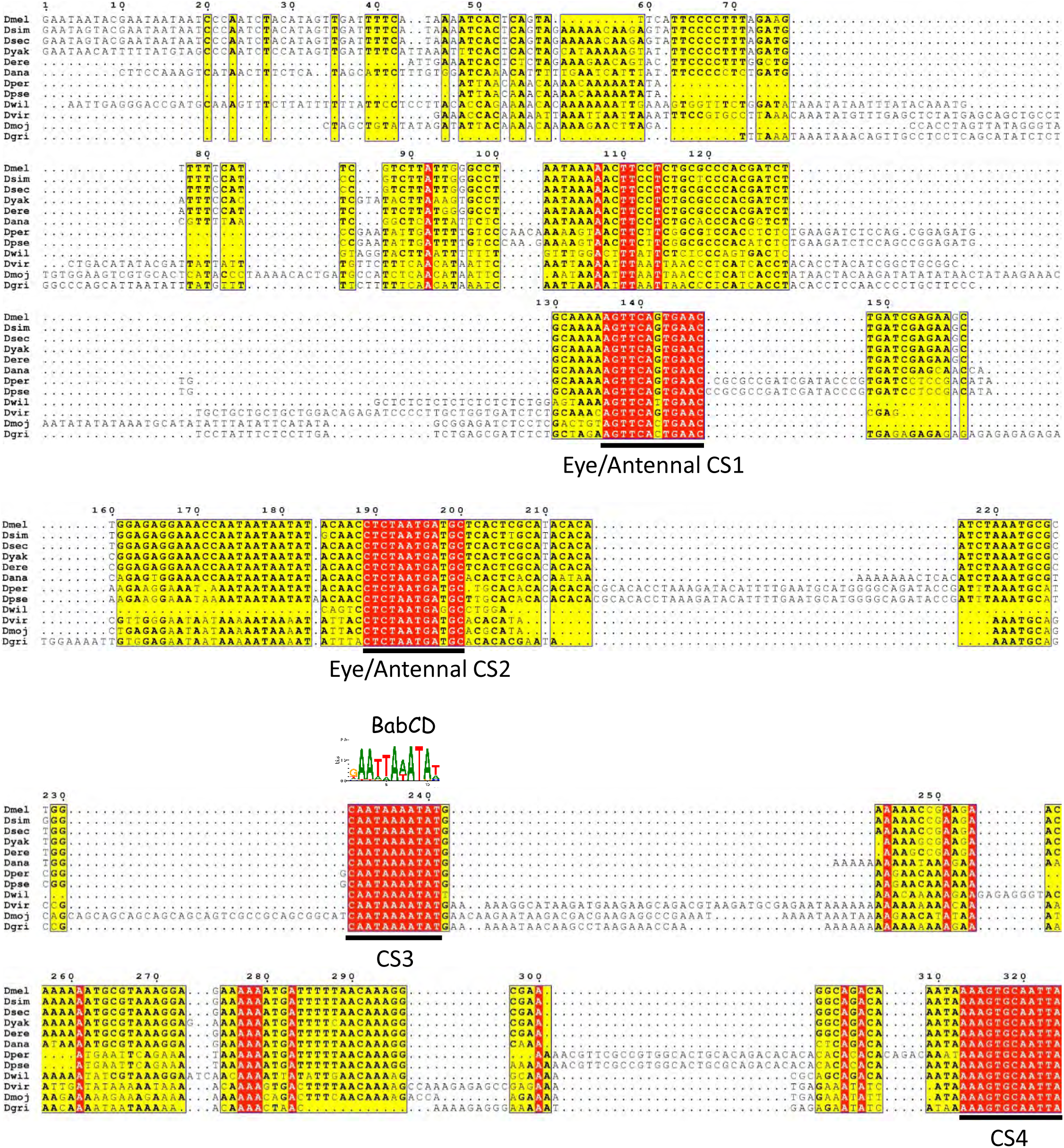

BER^OCS7^ sequence conservation among Drosophilidae (Part2)

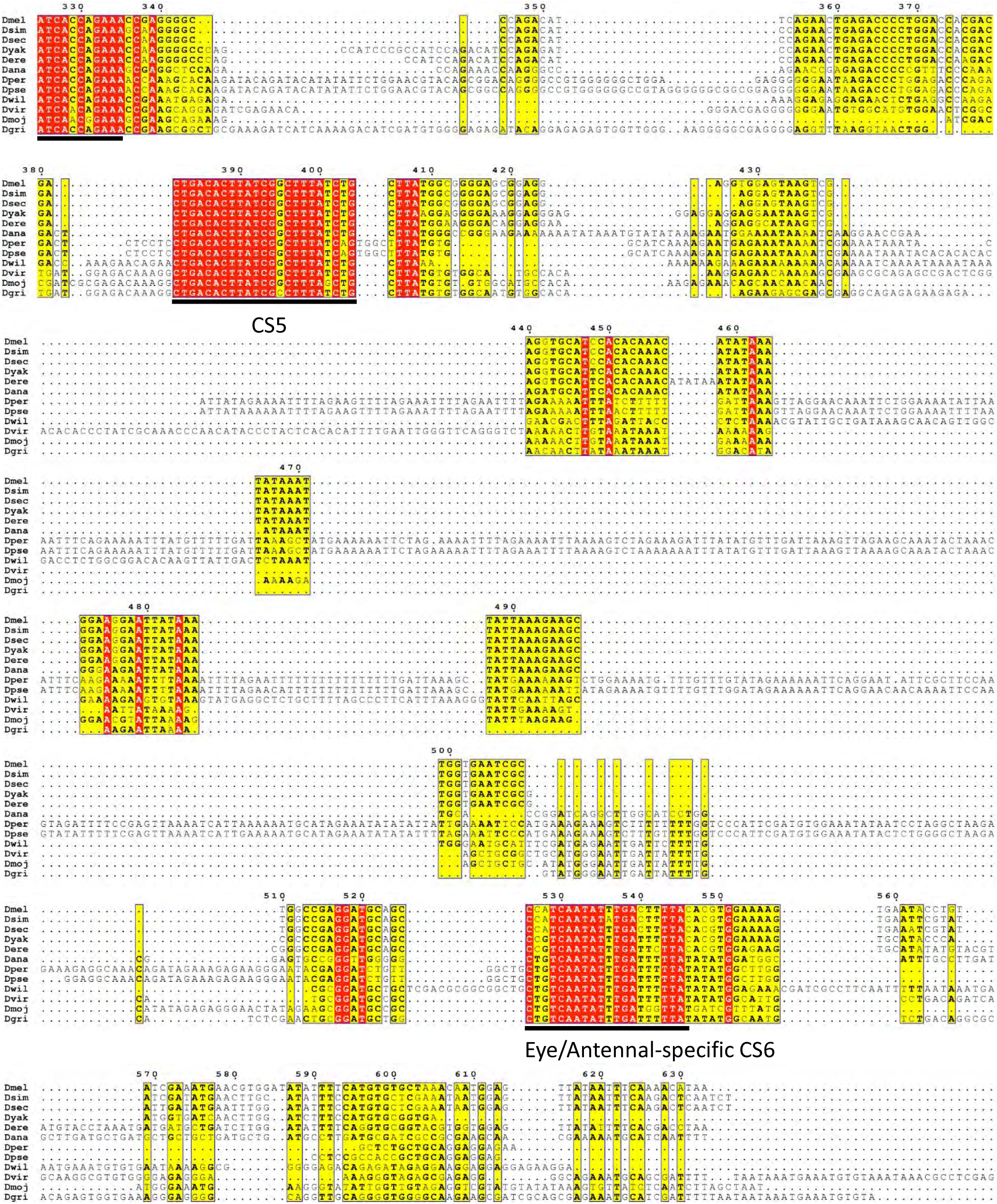

BER^OCS8^ sequence conservation among Drosophilidae (Part1)

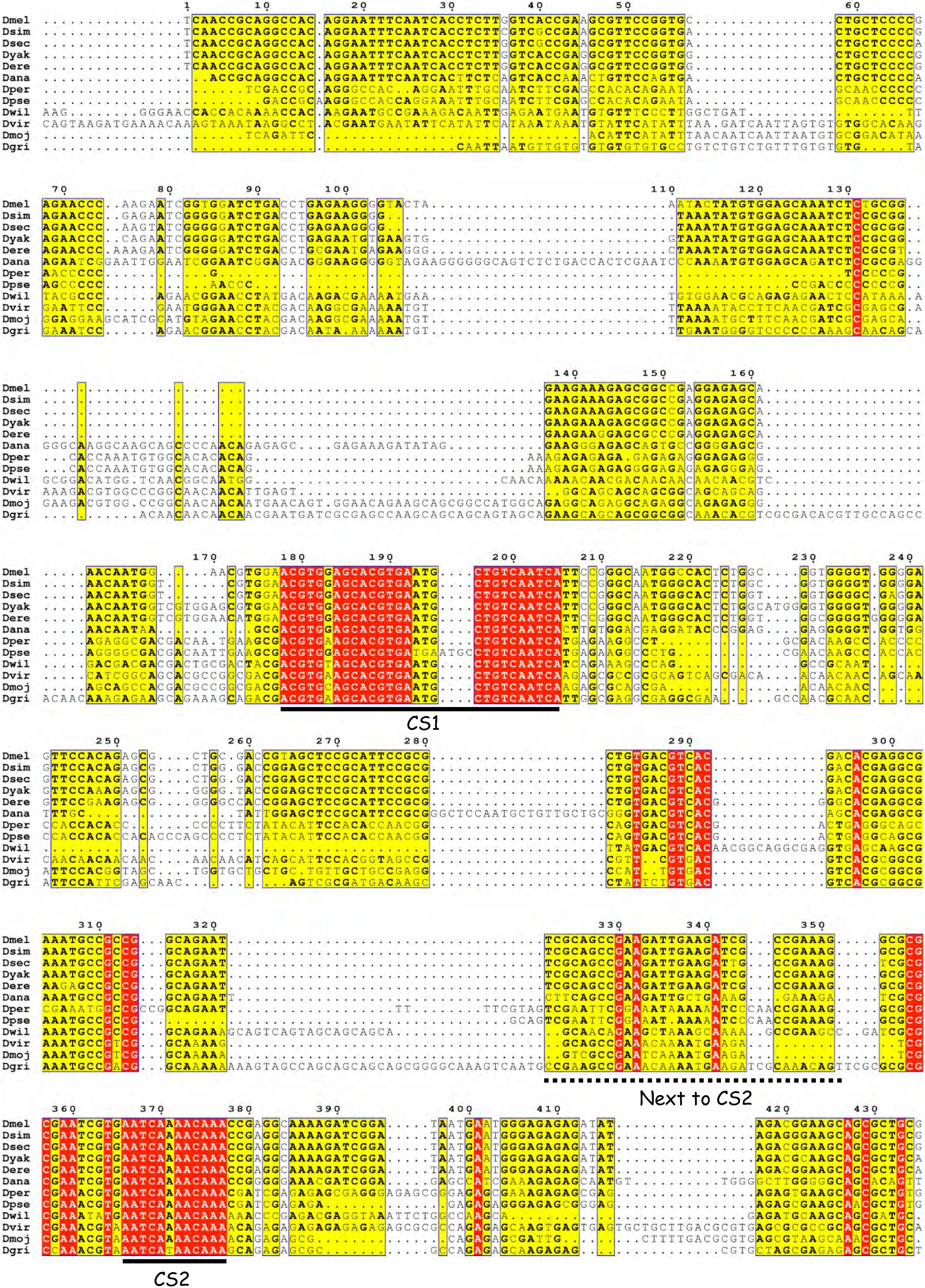

BER^OCS8^ sequence conservation among Drosophilidae (Part2)

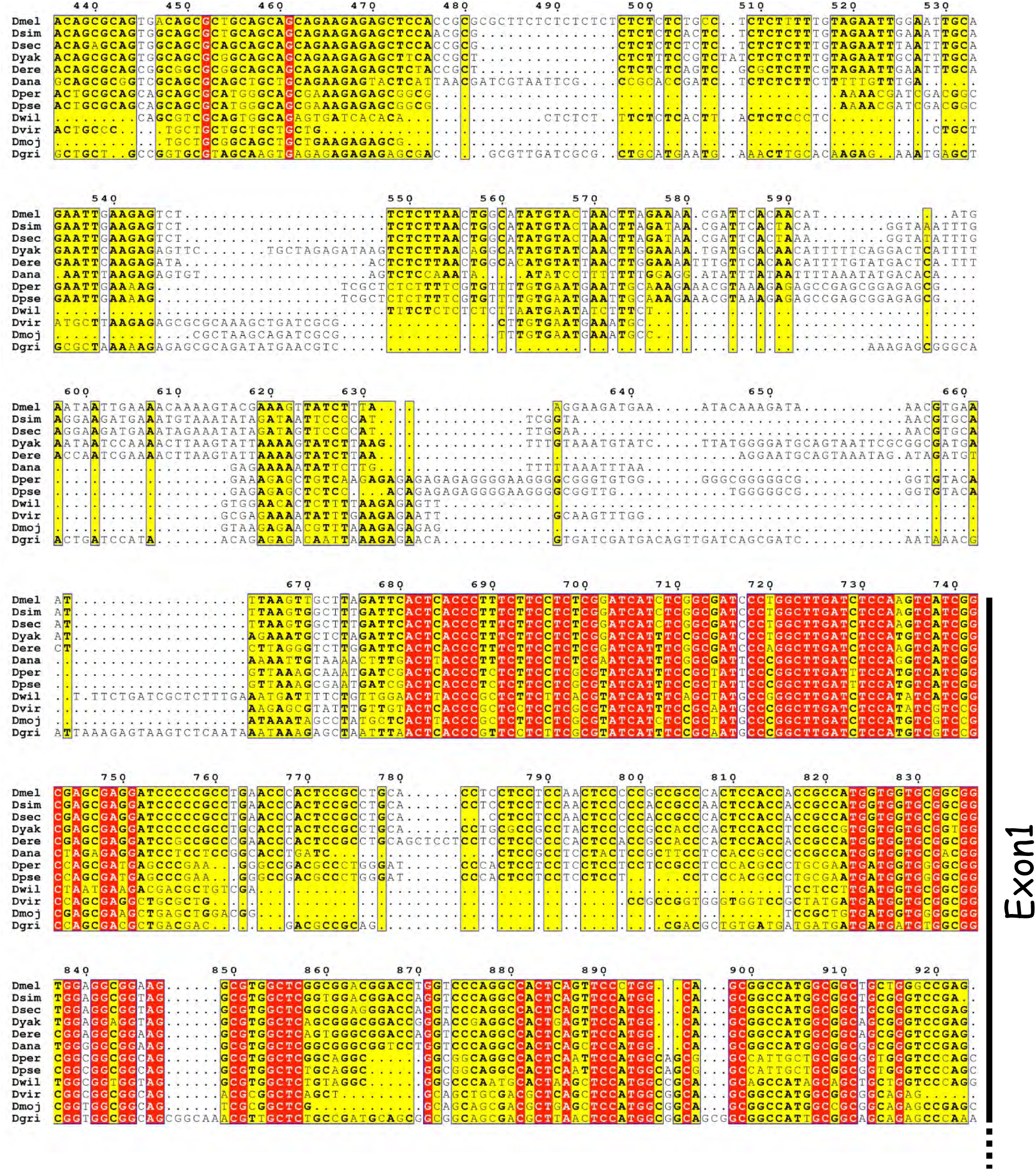

BER^OCS9^ sequence conservation among Drosophilidae (Part1)

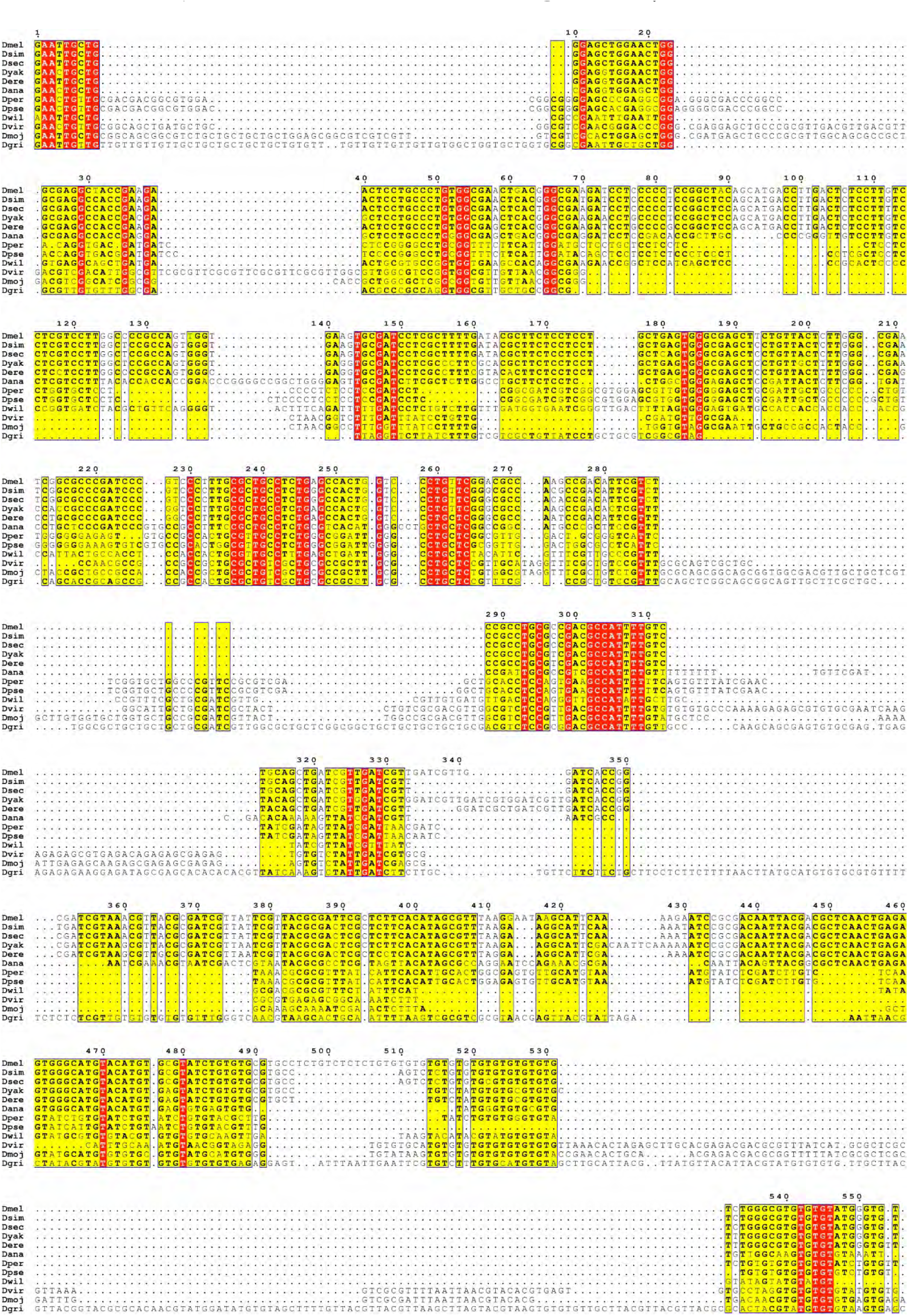

BER^OCS9^ sequence conservation among Drosophilidae (Part2)

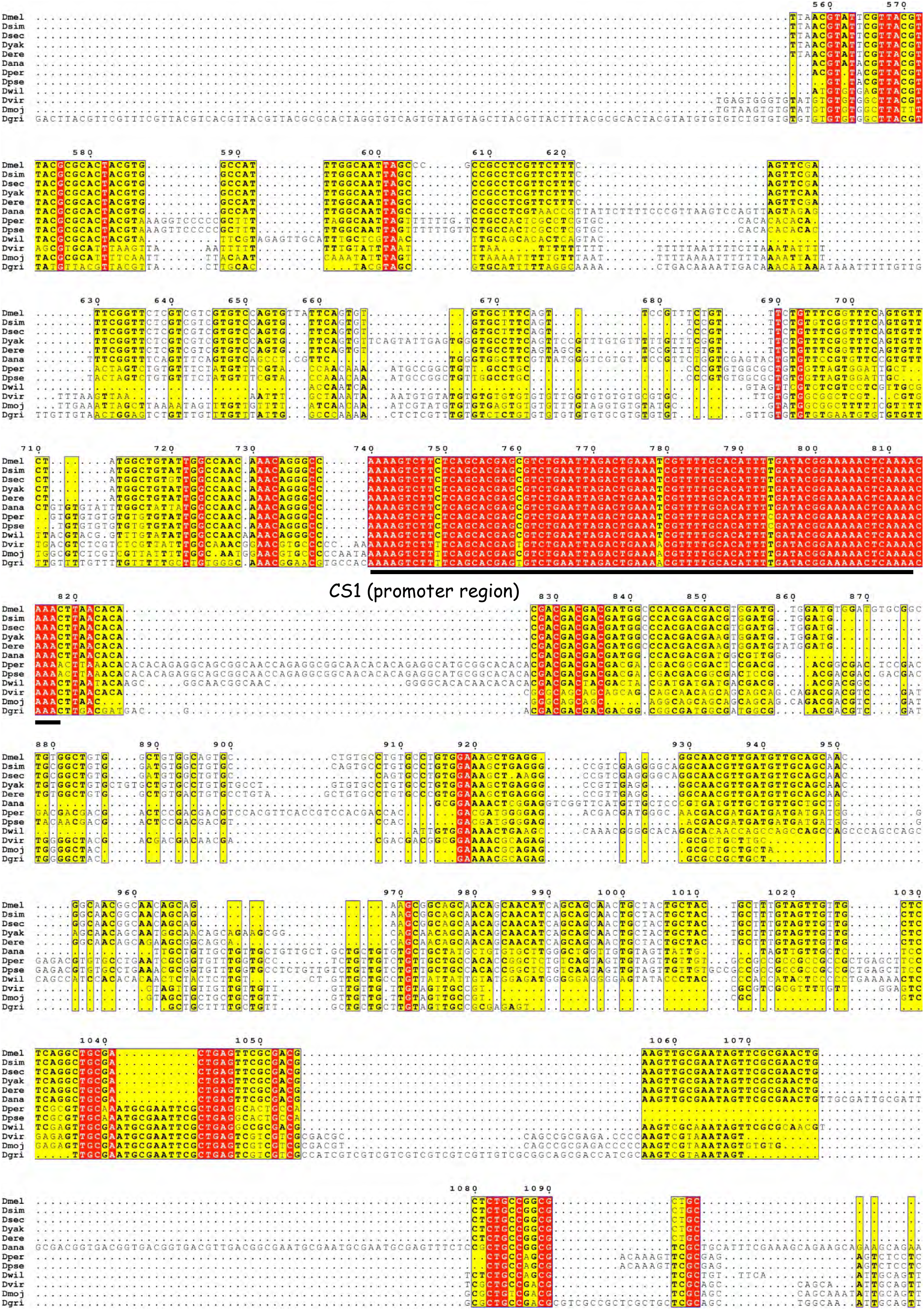

BER^OCS9^ sequence conservation among Drosophilidae (Part3)

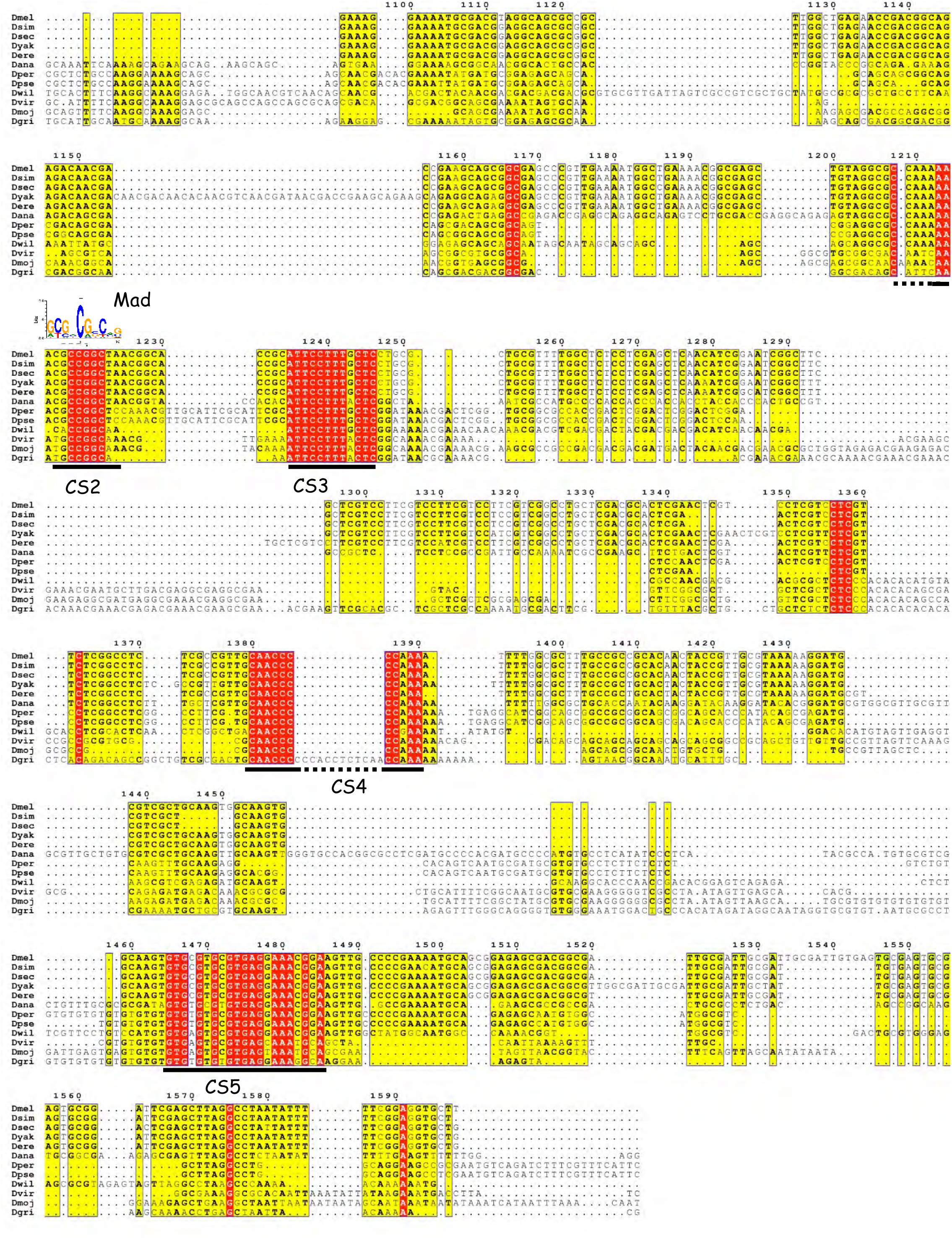

BER^OCS10^ sequence conservation among Drosophilidae (Part1)

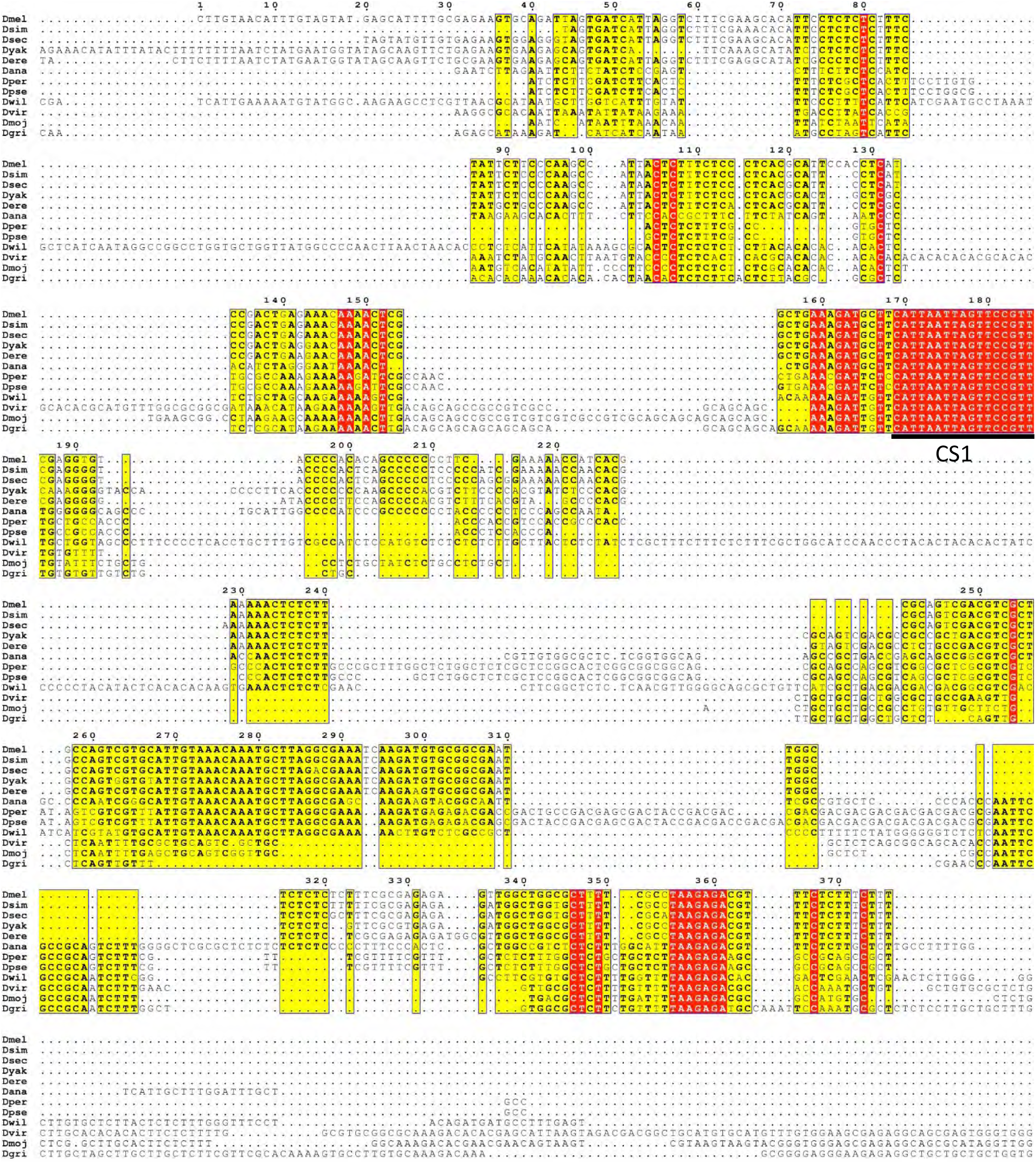

BER^OCS10^ sequence conservation among Drosophilidae (Part2)

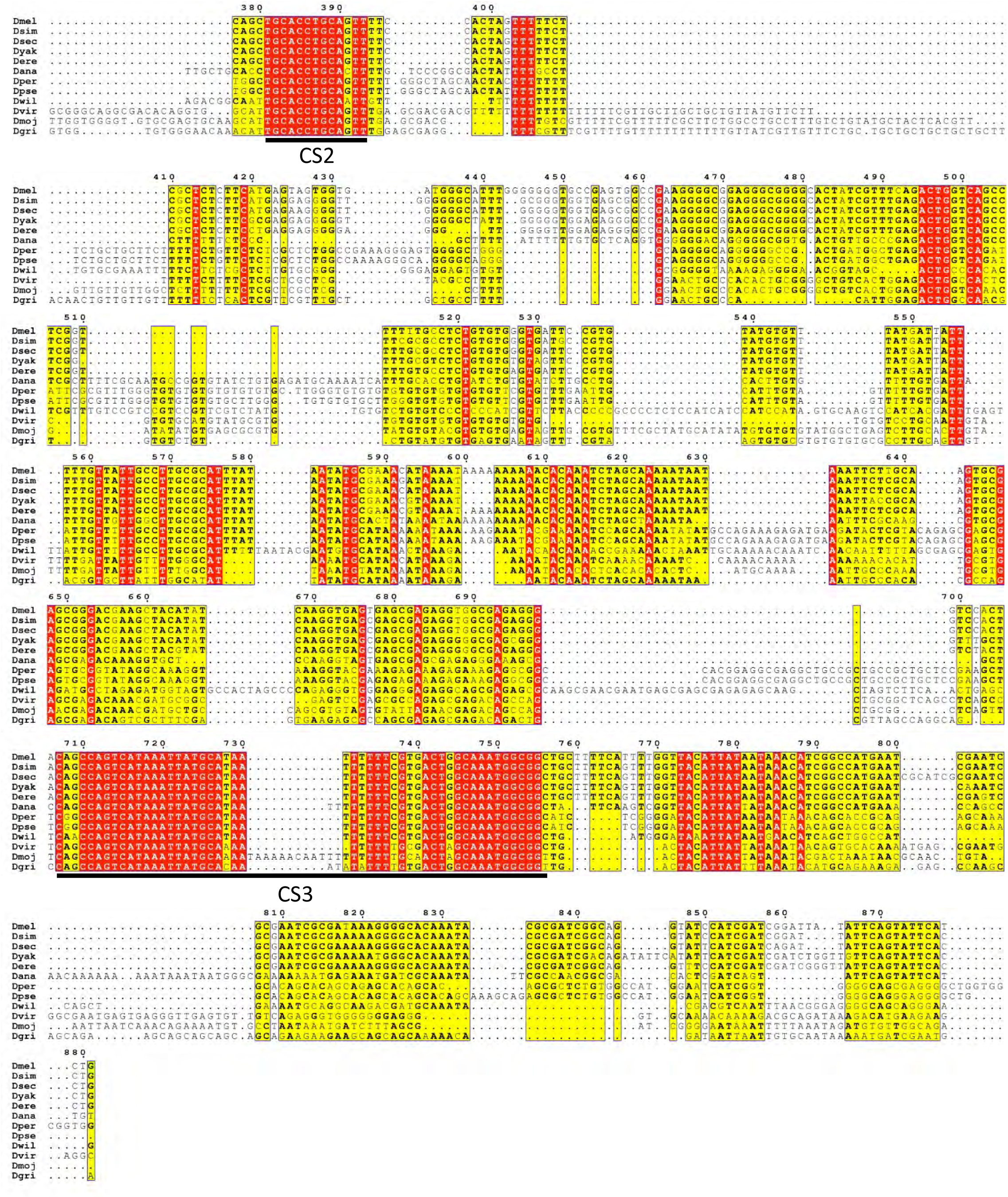

LAE sequence conservation among Drosophilidae (Part1)

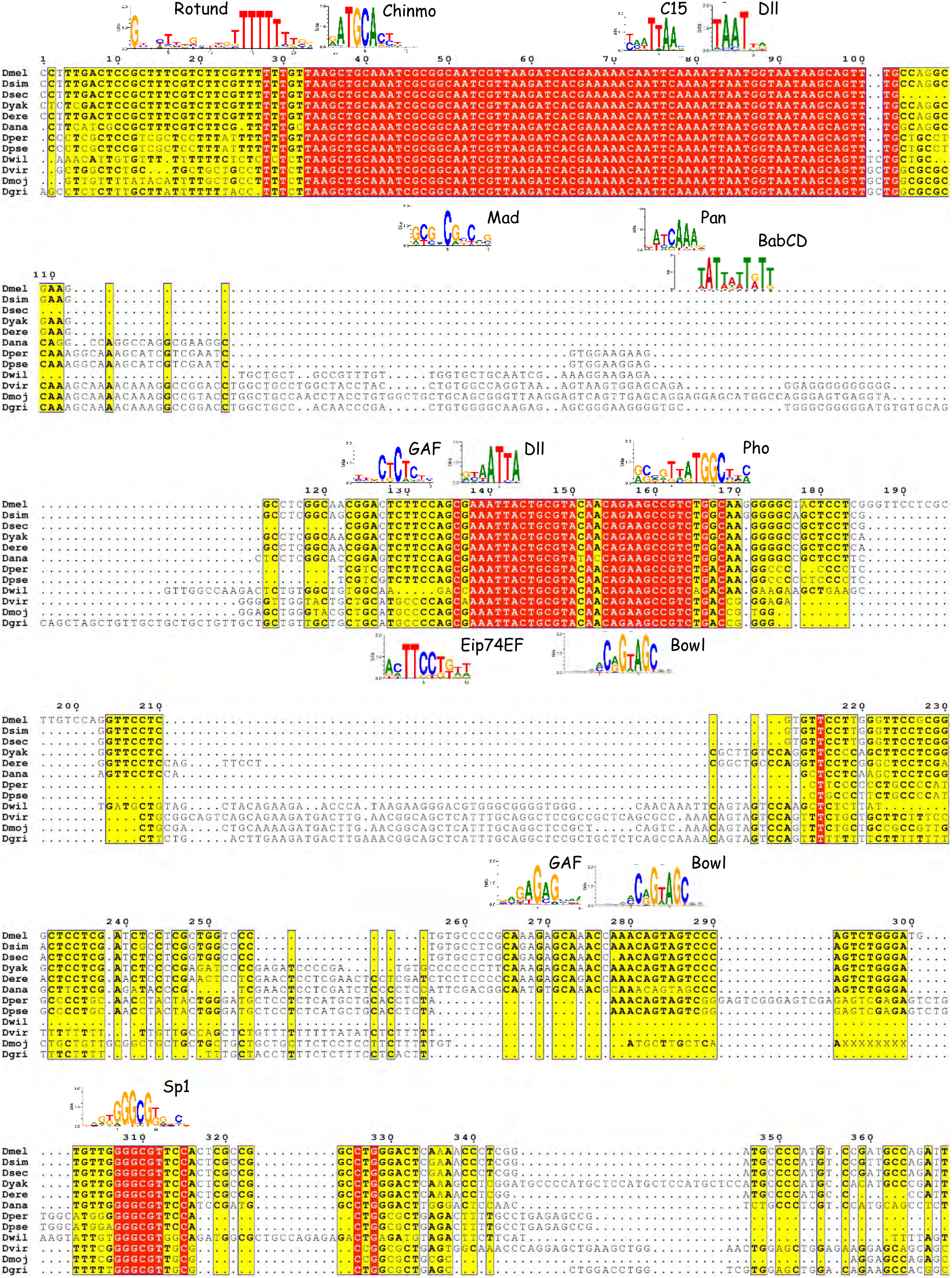

LAE sequence conservation among Drosophilidae (Part2)

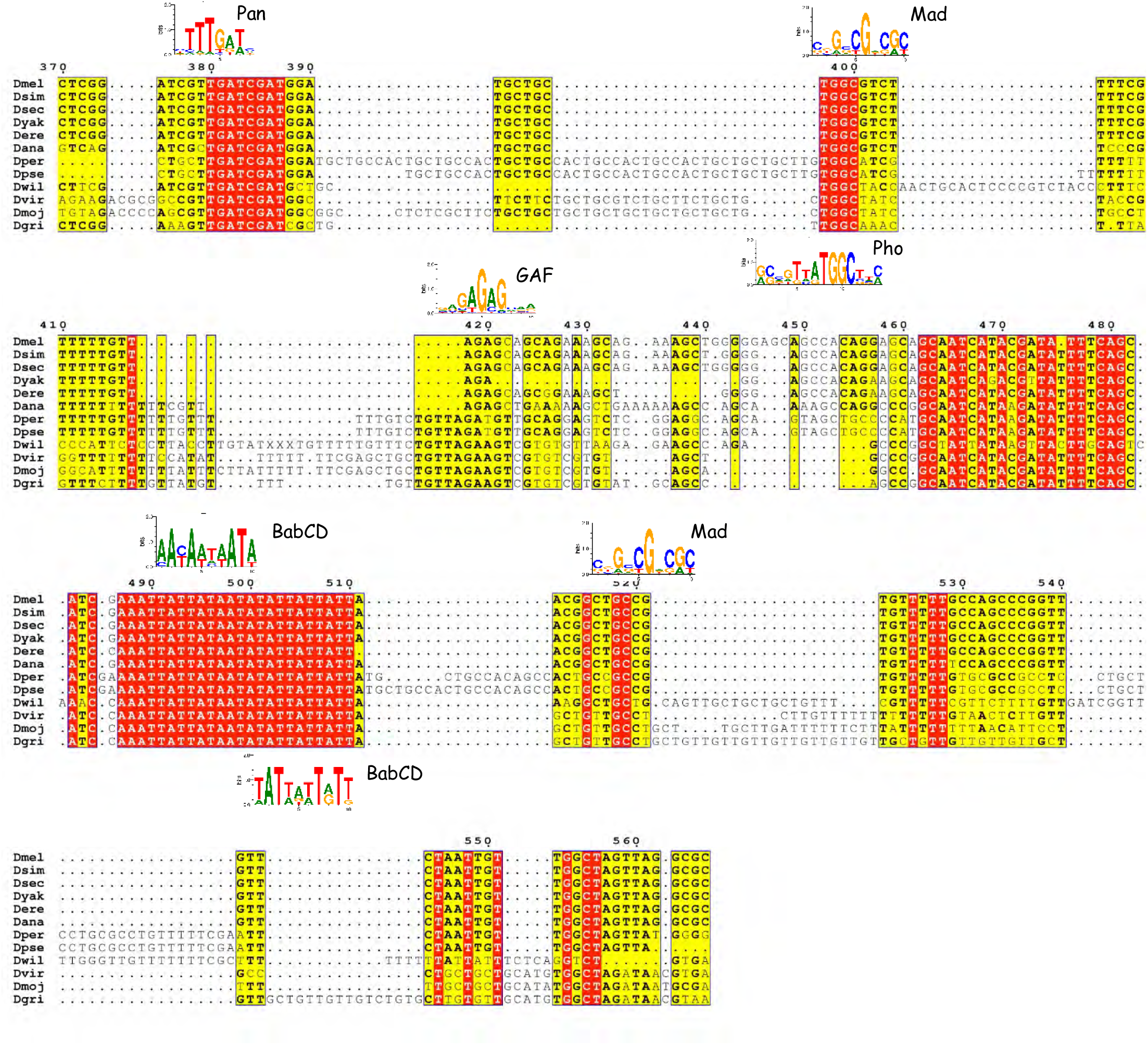

Cardiac CE sequence conservation among Drosophilidae (Part1)

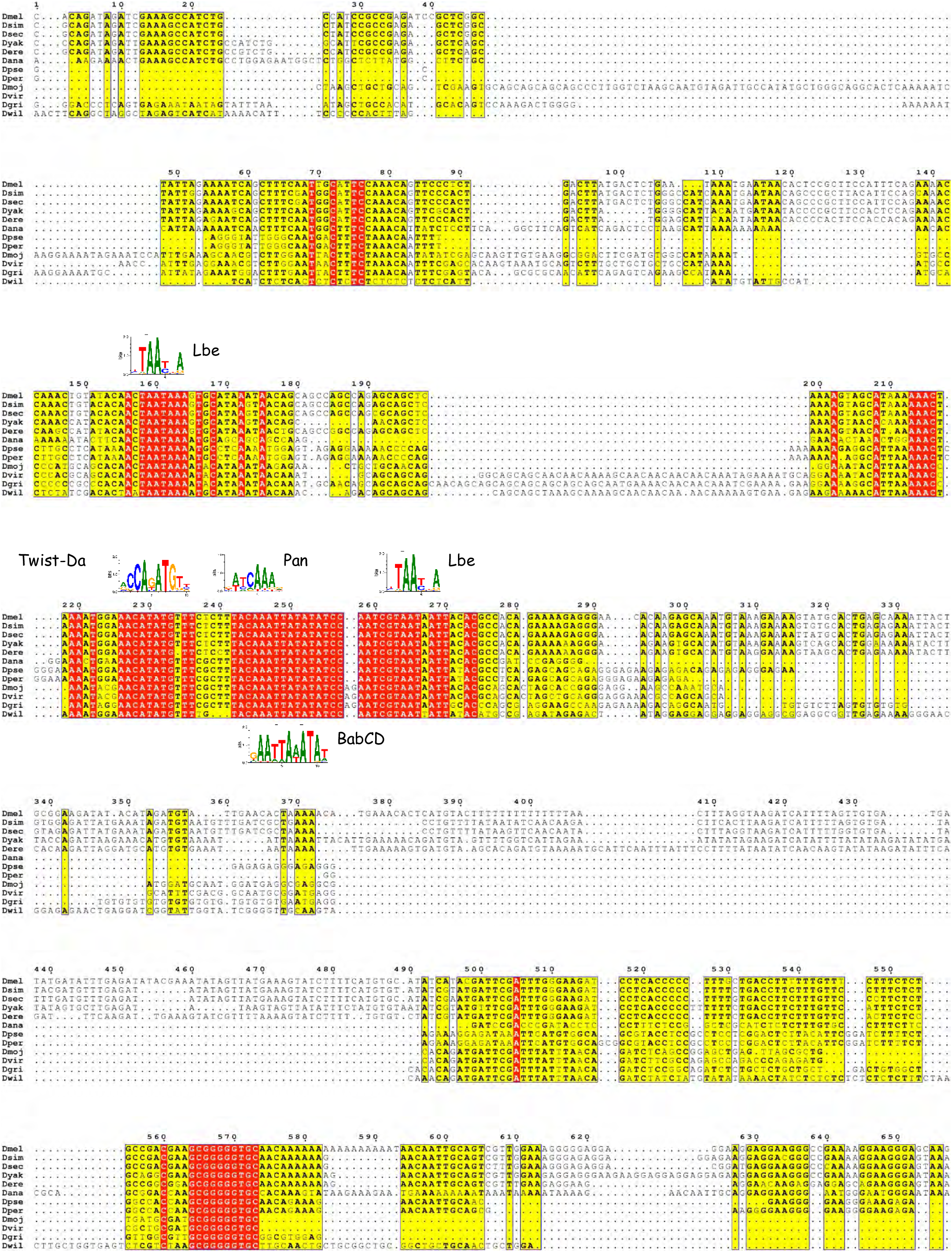

Cardiac CE sequence conservation among Drosophilidae (Part2)

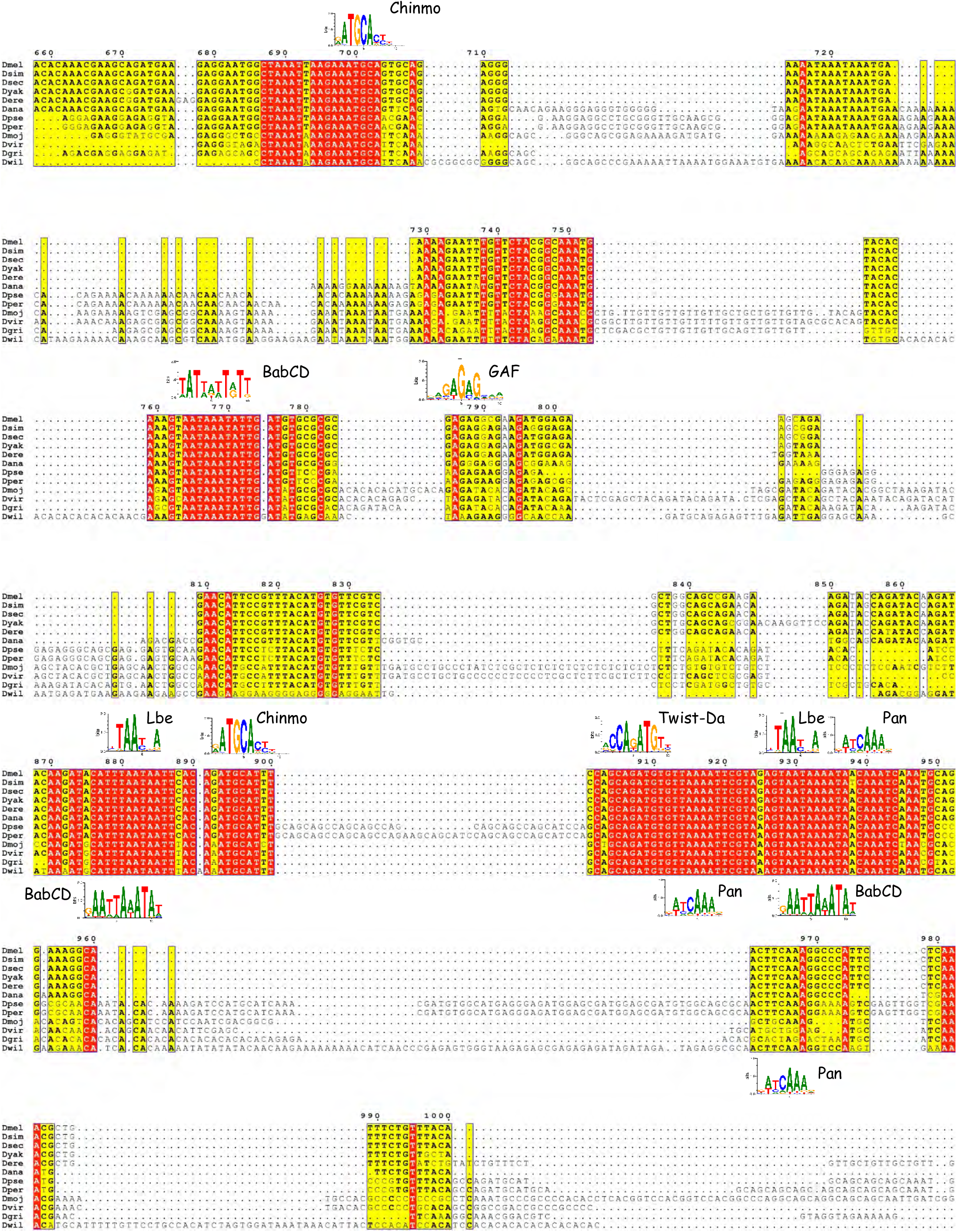

Abdominal AE sequence conservation among Drosophilidae (Part1)

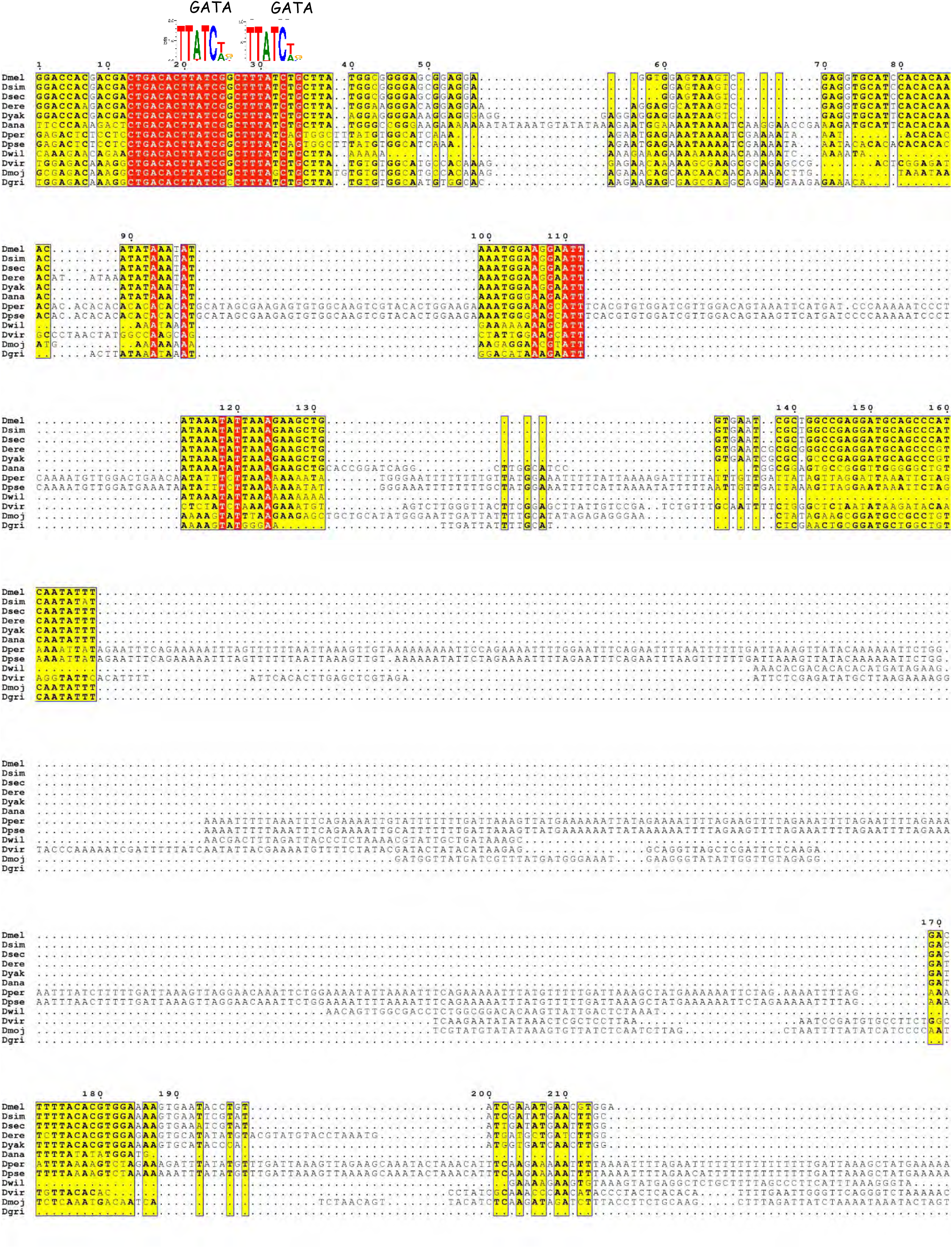

Abdominal AE sequence conservation among Drosophilidae (Part2)

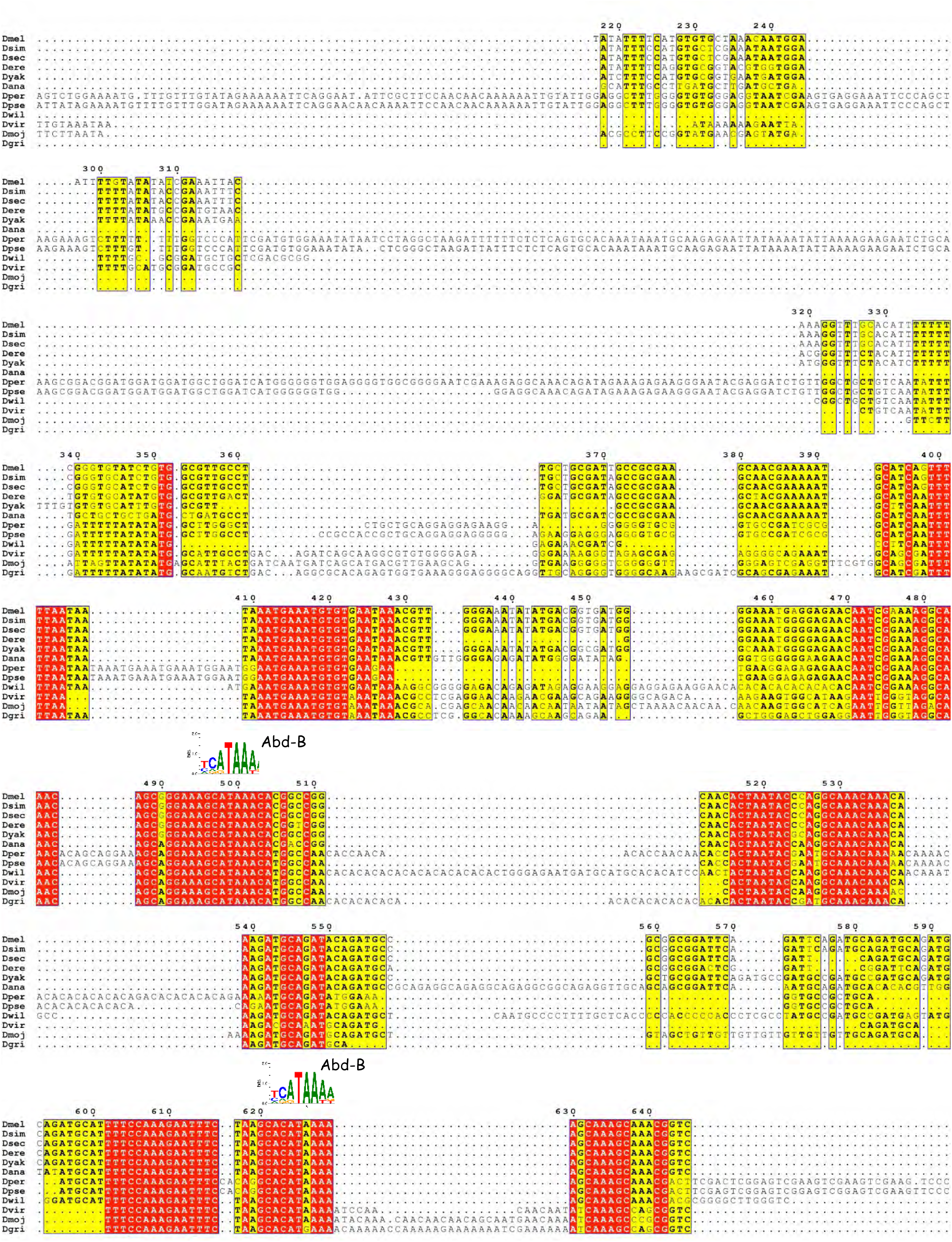

Abdominal AE sequence conservation among Drosophilidae (Part3)

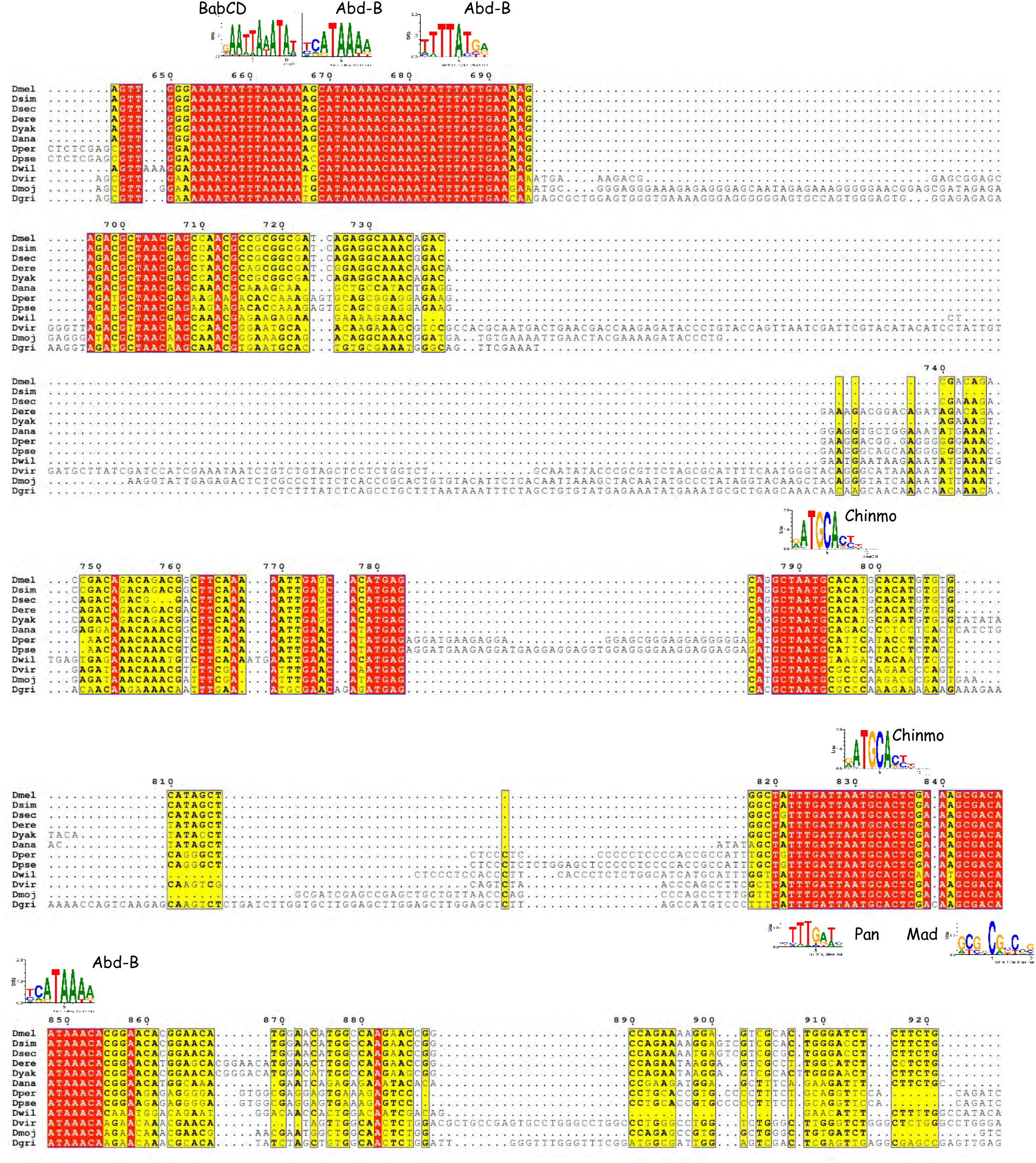

Abdominal AE sequence conservation among Drosophilidae (Part4)

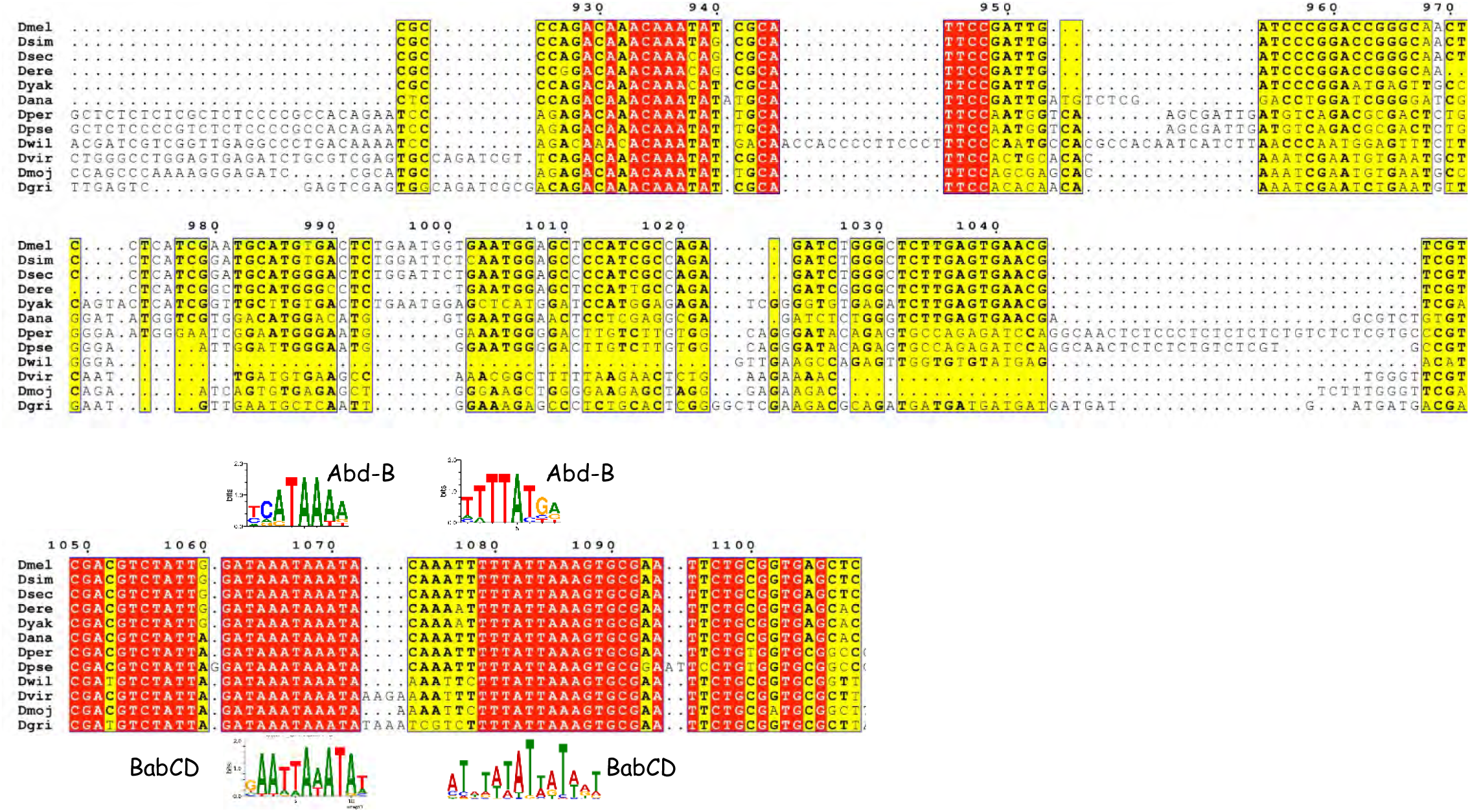

Abdominal DE sequence conservation among Drosophilidae (Part1)

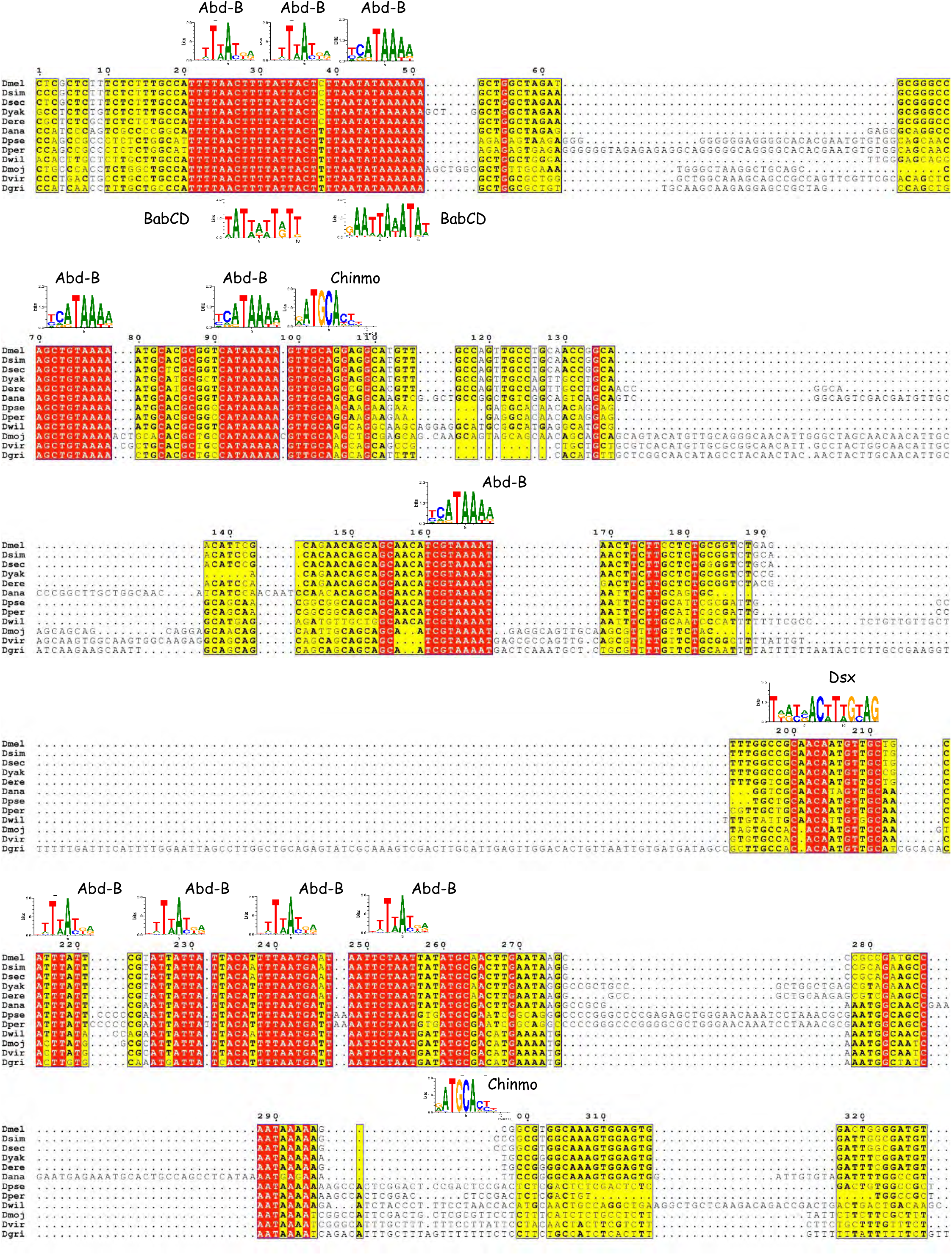

Abdominal DE sequence conservation among Drosophilidae (Part2)

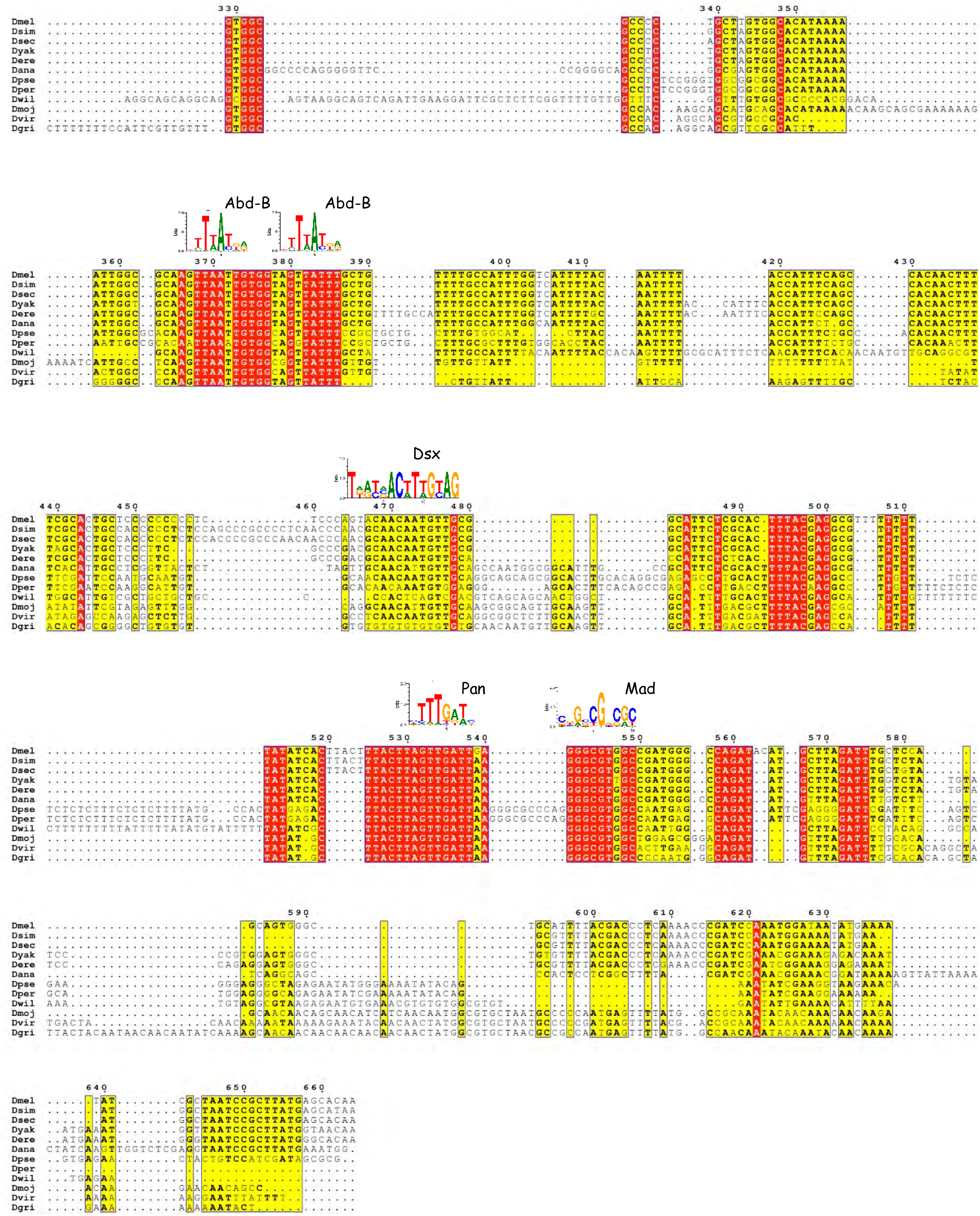

